# Transcriptome computational workbench (TCW): analysis of single and comparative transcriptomes

**DOI:** 10.1101/733311

**Authors:** Carol A. Soderlund

## Abstract

De novo transcriptome sequencing and analysis provides a way for researchers of non-model organisms to explore the differences between various conditions and species. The results are typically not definitive but will lead to new hypotheses to study. Therefore, it is important that the results be reproducible, extensible, queryable, and easily available to all members of the team. Towards this end, the Transcriptome Computational Workbench (TCW) is a software package to perform basic computations for transcriptome analysis (singleTCW) and comparative analysis (multiTCW). It is a Java-based desktop application that uses MySQL for the TCW database. The input to singleTCW is sequence and optional count files; the computations are sequence similarity, gene ontology (GO), open reading frame (ORF), and differential expression (DE). TCW provides support for searching with the super-fast DIAMOND program against UniProt taxonomic databases, though the user can provide other databases to search against. The ORF finder uses hit information, 5^th^-order Markov models and ORF length. For DE and GO enrichment, TCW interfaces with the R environment and an R script, where R scripts are provided for popular methods. The input to multiTCW is multiple singleTCW databases; the computations are homologous pair assignment, pairwise analysis (e.g. Ka/Ks) from codon-based alignments, clustering (bidirectional best hit, Closure, Best Hit, OrthoMCL, user-supplied), and cluster analysis and annotation. Both singleTCW and multiTCW provide a graphical interface for extensive query and display of the data and results. Example results are presented from two rhizome and one non-rhizome plant, where one of the rhizome plants has replicate count data from four tissues. The supplement describes how to reproduce all tables and figures. The TCW V4 software is freely available at https://github.com/csoderlund/TCW; the package contains the jar files, external software, and demo files.

## 1 Introduction

Given transcriptome sequences and read counts, there are various computations to be performed. There are also various computations to compare the transcriptomes of multiple species. These analyses are typically computed with various downloaded programs, web-based programs, spreadsheets and custom scripts. This ‘ad hoc’ style of analysis can lead to lack of reproducibility, human error, loss of data and results, and is inadequate for extensibility. The results are typically not definitive so a good graphical interface is necessary for exploring the data.

TCW is a desktop application that aids in creating reproducible and extensible results. It provides the standard computations for single transcriptome analysis (singleTCW) and for comparative analysis (multiTCW). TCW is written in Java and stores all data and results in a MySQL database. Both singleTCW and multiTCW provide interfaces for query and display, which allows the user to drill down to the details of the input data and results. TCW was first published in 2013 [1], but has been re-engineered and significantly enhanced since then. This publication discusses TCW V4.

For singleTCW, the input is sequence and optional replicate count files; the computations are sequence similarity annotation, GO assignment, ORF computation and differential expression. The user provides the annotation databases for the similarity search, where UniProt [2,3] is given special support for obtaining and viewing the results. With the freely available super-fast DIAMOND [4,5], the searching is no longer a bottleneck. The GO information, along with Interpro [6], KEGG [7], Pfam [8] and Enzyme numbers (EC) [9], are extracted from the UniProt files. GO names and relations are extracted from the go-basic.obo file. The ORF finder uses hit information, 5^th^-order Markov models and ORF length. For DE and GO enrichment, TCW interfaces with the R environment and an R script, where R scripts for edgeR [10], DEseq [11] and GOseq [12] are provided, or the user can supply their own.

For multiTCW, the input is one or more singleTCW databases; the computations are homologous pair assignment, clustering of homologous pairs, pairwise and cluster analysis. The clusters can be computed by best bidirectional hit (BBH), Closure, Best Hit, OrthoMCL [13], or loaded from a user-supplied file of clusters. For pairwise analysis, the codon-based alignment is computed for each pair in a cluster, from which statistics such as CpG, ts/tv, and Ka/Ks [14] are computed. For the cluster analysis, the multiple sequence alignments (MSA) is computed using MAFFT [15], from which two MSA scores are computed. Additionally, the majority annotation is assigned to each cluster.

To demonstrate the TCW analysis, the software is applied to the transcriptomes of two rhizome plant species (*Oryza longistaminata* and *Nelumbo nucifera*) and one non-rhizome species (*Oryza sativa)*. The red rice (*O. longistaminata*) contigs were de novo assembled and have replicate counts from rhizome, root, stem and leaf [16]. The transcript files for sacred lotus (*N. nucifera*) and the cultivated rice (*O. sativa*) were downloaded from NCBI.

## 2 Materials and Methods

### 2.1 TCW requirements and overview

TCW is available at https://github.com/CSoderlund/TCW. The package requires Java and MySQL (or MariaDB) to be installed. If differential expression will be computed, it requires R along with the necessary R packages for computing DE. The downloadable TCW package contains demo files and all necessary external software compiled for Linux and MacOS.

#### Input details

TCW takes as input one or more FASTA files of nucleotide or amino sequences, where each sequence file may have an associated tab-delimited file of counts. TCW can be used with transcriptomes with replicate count data, and can also be used with proteomes with replicate spectra (count) data. The ability to use TCW with both transcriptome and proteome data is useful when there are dual RNA-seq and peptide experiments, where the results can be equivalently analyzed with TCW, e.g. He et al. [17]. TCW has an assembly algorithm for Sanger ESTs [18], 454 data or a combination of these with pre-assembled contigs [1]; the input sequence files may have associated files of quality values, which are used in the assembly. TCW can take as input location information, which is useful with gene models. Due to its multiple types of input, TCW generically refers to the input as “sequences with optional counts”.

#### Test machines

TCW uses the BLAST[19], DIAMOND and MAFFT programs, which use multi-processors; all the TCW processing is single-process. TCW was tested on two machines: (i) Linux x86-64 24-core 2.3Ghz AMD with 128 gigabytes of RAM (purchased 2011), which used MariaDB 10.4.12. (ii) Mac 10.9.5 with 3.2Ghz 6-core and 64 gigabytes of RAM (purchased 2019), which used MySQL v1.8. Timing results are presented in the Results section; S1 Suppl §3 provides more details about the test machines, detailed timings and memory usage. The necessary size of the machine all depends on the number of input sequences and annotation databases. Once the databases are built, they can be queried on any machine that has Java and MySQL.

#### Demo files

The downloadable TCW package contains the following demo sets: (i) input sequences to be assembled, (ii) protein sequences with replicate counts, (iii) nucleotide sequences with replicate counts, and (iv) three datasets that have good homology for input to multiTCW. It also contains a subset of taxonomic UniProts and the go-basic.obo file to use for annotation. The TCW software can be experimented with by downloading the package, untar’ing it, entering the MySQL information in the HOSTs.cfg file, and running it on the demo datasets. Step-by-step instructions are provided at www.agcol.arizona.edu/software/tcw along with additional information.

#### Overview

TCW has four graphical interfaces to build the single database (runAS, runSingleTCW, runDE) and multiple database (runMultiTCW). Table 1 shows the salient terminology. Figure 1 shows the workflow.

**Table 1.** Terminology used by TCW.

| Term | Description |
| --- | --- |
| TCW-GOdb | MySQL database created by runAS, which contains all GO terms from the go-basic.obo file and associated information from the downloaded UniProt .dat files. |
| sTCWdb | MySQL database created by runSingleTCW, which contains the input sequences, optional counts and the analysis results. |
| mTCWdb | MySQL database created by runMultiTCW, which contains the data from multiple sTCWdbs and the comparative analysis results. |
| Sequence | The input sequences to TCW, e.g. transcripts, proteins, gene models. |
| annoDB | Annotation database, which is a file of protein or nucleotide sequences to search against. |
| SP-taxo | SP is SwissProt, taxo is the taxonomy, e.g. SP-plants is the SwissProt plants database. The same naming scheme is used for TR (TrEMBL). |
| Hit | A match between two sequences found by a search program (e.g. BLAST or DIAMOND). |
| Annotated sequence | A sequence that has one or more hits to any annoDB. |
| Best Bits | The annoDB hit for a sequence that has the best bit-score and E-value. |
| Best Anno | The annoDB best bit-score hit for a sequence where the hit description does not contain phrases such as “uncharacterized protein”. |
| Rank=1 | The annoDB best bit-score hit for a sequence to a given annoDB. |

**Figure 1.**
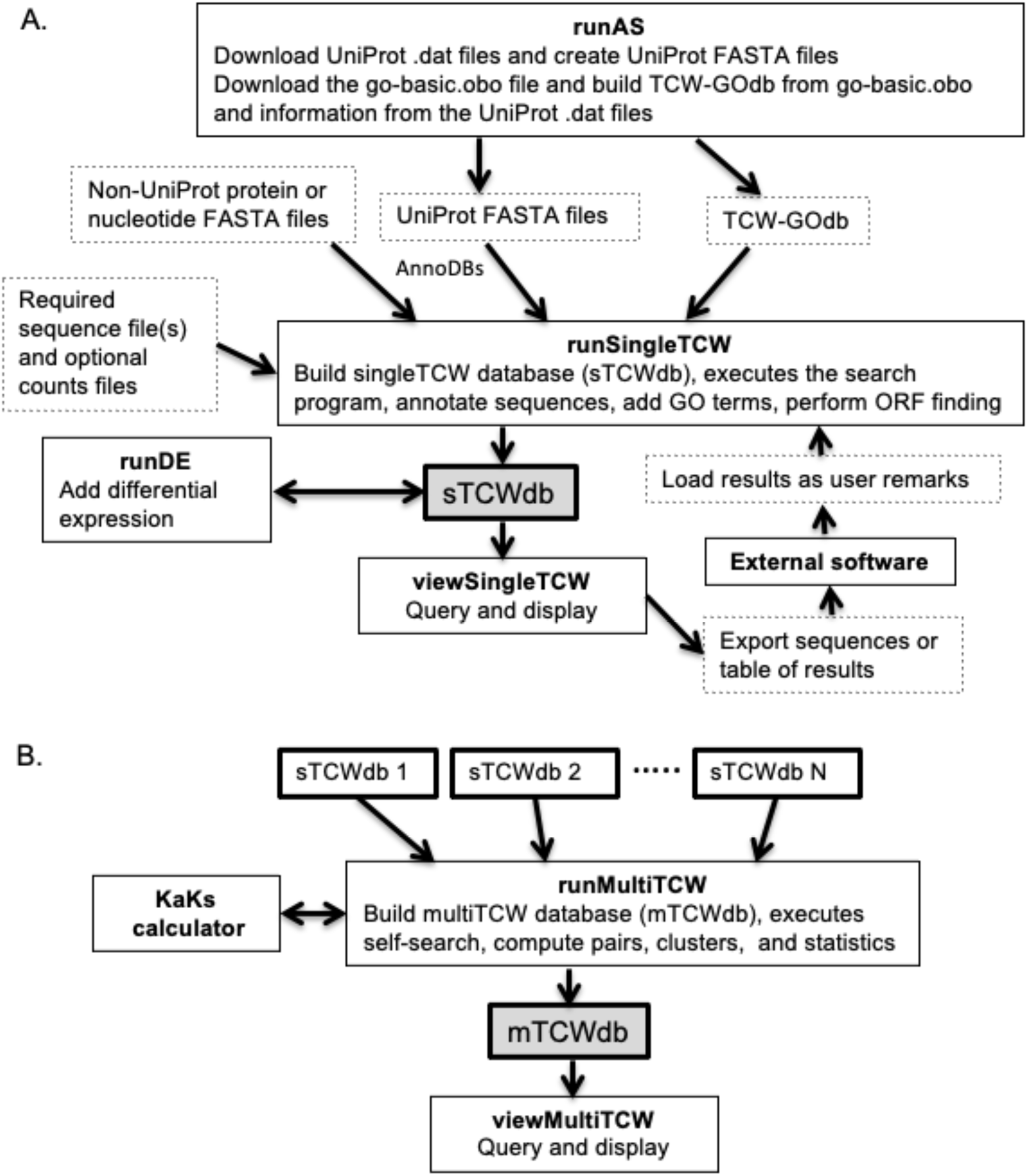
TCW programs and dataflow. (A) The steps for creating a singleTCW database. The only required input is the FASTA formatted sequence file(s). (B) The steps for creating a multiTCW database. The only required input is N (N≥2) sTCWdbs, where N is not a large number (recommended ≤ 4, though there is no TCW limit).

### 2.2 Create annoDBs and TCW-GOdb (runAS)

Multiple annotation databases (annoDBs) can be used for sequence similarity, where an annoDB is a protein or nucleotide FASTA formatted file. TCW offers specific support for the UniProt taxonomic databases with a graphical interface called runAS, which will download the desired UniProt .dat data files and create the corresponding FASTA file of sequences. RunAS will also create a subset SwissProt, which is the entire SwissProt minus all protein records from the downloaded taxonomic databases. The advantage of using the taxonomic databases is that the most relevant SwissProt and TrEMBL databases can be used and their hits can be queried by taxonomy using viewSingleTCW.

The runAS interface is used to create the TCW GO database (TCW-GOdb). It loads from the go-basic.obo file the GO terms, names, relations, and GO Slims, where it only considers ‘is_a’ and ‘is_part’ relations. RunAS also loads the relevant information from the downloaded UniProt databases. The UniProt .dat files contain for each record the GO “direct” assignments with their evidence code, InterPro [6], KEGG[7], Pfam [8] and Enzyme Commission (EC) [9], where these data items are loaded into the TCW-GOdb.

### 2.3 Build a singleTCW database (runSingleTCW)

The runSingleTCW program provides a graphical interface to build the MySQL database (sTCWdb). The first step is to load the sequences and optional counts into the database. By default, TCW assign names, though the user can request that the original names be used (if the names are assigned, the original name is stored for display in viewSingleTCW). The replicate counts are summed and the Transcripts Per Million (TPM) calculated. The rest of this section describes the annotation process where the details are provided in the S2 Suppl.

#### 2.3.1 Sequence similarity annotations

For similarity searching, TCW can use BLAST or DIAMOND. Given that DIAMOND is much faster than BLAST, it is the TCW default search program. From all hits for a sequence, a best bit-score hit (Best Bits) and best annotation hit (Best Anno) will be computed, where the Best Anno is the best hit with a meaningful description (e.g. not “Uncharacterized protein); see S2 Suppl §2.1.1 for details. Figure 2 shows an example where the Best Bits hit is ‘uncharacterized’ and the Best Anno is the SwissProt “Ureidoglycolate hydrolase”.

**Figure 2.**
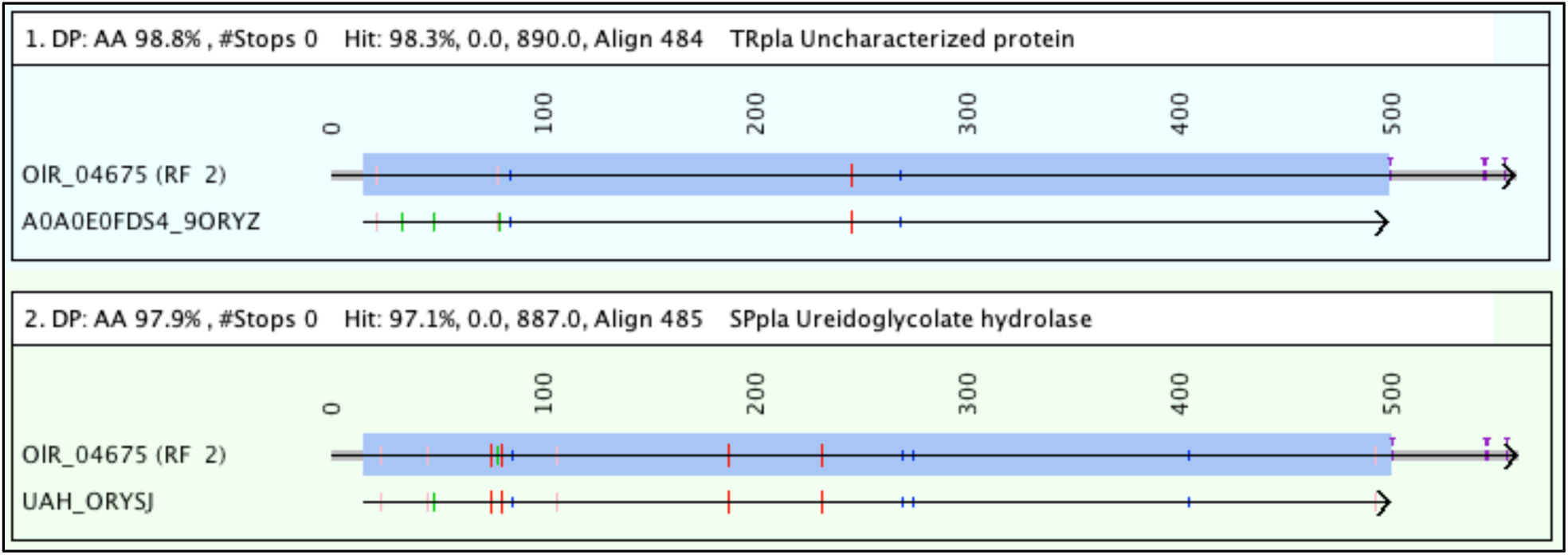
TCW alignment of Best Bits (bit-score) and Best Anno (annotation). A dynamic programming algorithm is used for the alignment, where the DIAMOND hit is highlighted in blue. The green marks are gaps, and the rest of the marks indicate different amino acids colored by their BLOSUM62 [20] score; red is BLOSUM < 0, pink is BLOSUM = 0, and dark blue is BLOSUM > 0. The purple T’s are stop codons. The gray areas at the ends are ‘overhangs’. The top alignment has the higher bit-score of 890 whereas the bottom alignment has a lower bit-score of 887 but the description is informative. The view can be changed to show the text alignment.

##### Prune hits

Given the growing size of the UniProt taxonomic databases, there can be many protein hits with the same alignments and/or descriptions. For most uses, it is only necessary to keep the best hit per annoDB with a given alignment or description; an exception would be if the goal was to obtain all species that hit each sequence. TCW has an option to prune hits based on the same alignment, where it arbitrarily retains one of the hits. There is also an option to prune hits based on similar descriptions, where it retains the hit with the best bit-score and E-value.

#### 2.3.2 GO annotation

For the UniProt hits in the sTCWdb, their associated data is copied from the TCW-GOdb to the sTCWdb, where the data is InterPro, KEGG, EC, Pfam and direct GO values. Each hit is assigned the direct Gos along with all inherited GOs. If a GO is not directly assigned or inherited by a hit, it will not be in the sTCWdb.

As discussed by Rhee et al. [21], assigning levels to the GO terms is problematic because the GO structure is not uniform and each GO term can exist at multiple levels. Nevertheless, it is common to use levels since it provides the scientist an indication of where a given GO term is in the directed acyclic graph; therefore, TCW assigns the maximum level to each GO term. TCW also assigns to each GO term the number of sequences with at least one hit that is assigned or inherited the GO.

#### 2.3.3 ORF finding and GC content

If a sequence has a best hit that passes the user-supplied E-value and similarity cutoff, the ORF finder will use the frame of the hit for the ORF. For the ORF coordinates, it uses the hit coordinates if the hit region ends with a valid start and stop codon, otherwise, it uses heuristics to extend the hit region to the best start and end for the ORF. If the sequence does not have a good hit, (i) the best ORF is found for each of the six reading frames and (ii) the best ORF from the six frames is selected. In both cases, the best ORF is selected based on the ORF length and Markov score. To compute the Markov score, the TransDecoder v5.5.0 [22] algorithm was translated from Perl to Java to be used in the TCW ORF finder. The 2000 longest sequence regions from the best hits are used to train the Markov model. ORF finding is complicated by transcripts with hits in multiple frames or with stop codons in the hit region, so the algorithm uses heuristics for these cases. These heuristics along with details of the algorithm are discussed in S2 Suppl §2.2.

The computation of the GC content is performed on each sequence. For the overview, the average GC and CpG content are computed for the coding sequence (CDS) and untranslated regions (UTRs) derived from the computed ORF.

#### 2.3.4 Other features

##### Self-search

There is an option to compare all input sequences, where the initial comparison uses BLASTn (NT), tBLASTx (6-frame), and/or BLASTp (translated ORFs); the highest scoring N pairs are aligned using dynamic programming, where N is a user-defined parameter. This feature is especially useful for evaluating de novo assembled contigs for highly similar sequences.

##### User-supplied remarks

Transcriptome and proteome studies generally have additional computations that are problem-specific; for example, transcripts are often analyzed for simple repeats. To use data and results from the TCW database, a file of results can be exported from viewSingleTCW for input to other programs, and then the external results can be imported into the sTCWdb as user remarks using runSingleTCW. The user remarks can be searched and viewed in viewSingleTCW. Additionally, location data can be entered into the database for display.

### 2.4 Differential expression (runDE)

The TCW runDE program is used to compute the differential expression from the replicate counts and enter the results into the database. The user can select to have the sequences pre-filtered with either of the following (N and M are user-supplied values): (i) the counts per million (CPM) filter removes sequences that do not have CPM > N for ≥ M samples, where CPM = (count/sample size) * 1E06, (ii) the count filter removes sequences that do not have any sample with count > N. To compute DE, runDE writes the necessary data to the R environment, runs an R script, and loads the results into the database. R scripts for edgeR [10] and DEseq [11] are provided; alternatively, the user can supply an R-script or a file of DE values. The results are entered into the sTCWdb with a user-supplied DE column name.

The runDE program provides the ability to compute enriched GOs for each DE column. Similar to running a DE script, it writes the necessary data to the R environment, runs the R script, and loads the results into the database. It allows any R script to be used, where by default it runs the GOseq [12] script. GOseq detects enriched GOs based on a binary vector representing the sequences with DE p-values < N (default 0.05) along with the sequence lengths.

### 2.5 Build a multiTCW database (runMultiTCW)

The runMultiTCW program takes as input two or more sTCWdbs and builds the multi-species database (mTCWdb) (Figure 1B). Though there is no upper limit on the number of sTCWdbs to be compared, it is not meant for a large number (recommended ≤ 4). The input sTCWdbs can be built from nucleotide or protein sequences. The following will discuss an mTCWdb built from nucleotide sTCWdbs. When the mTCWdb is built, the nucleotide sequences, ORF coordinates, translated ORFs, TPM, DE, and top hits are copied from the sTCWdbs to the mTCWdb. The GC and CpG content are computed for the nucleotide sequence, and CpG Obs/Exp [23] is computed for the CDS and UTRs, where the CDS and UTRs are derived from the TCW computed ORF. Optionally, the GOs for the hits can be copied to the mTCWdb. The rest of this section describes the clustering and annotation process where the details are provided in the S3 Suppl.

#### 2.5.1 Pairs and clustering

A self-search of all translated ORFs is performed along with an optional self-search of the nucleotide sequences. Either DIAMOND or BLAST can be used for the amino acid comparison; BLAST is required for the nucleotide comparison. The default TCW parameters are discussed in S3 Suppl §3.1.

The self-search tabular files are parsed and the pairs are loaded into the database. The pairs are used as input to the following cluster algorithms: (i) BBH (bidirectional best hit), (ii) Closure, (iii) Best Hit, (iv) OrthoMCL [13]. The first three options are implemented within TCW and have the following parameters: minimum percent similarity and minimum percent coverage over one or both sequences. The fourth option executes the OrthoMCL code from within runMultiTCW and loads the results; this option has the one OrthoMCL parameter of ‘inflation’. A fifth option is available, which allows the user to provide a file of clusters.

BBH is a common approach to use. Since more than two sTCWdbs may be compared, TCW provides N-way BBH, which first computes the 2-way BBH and then combines N-way BBH pairs. Alternatively, the user can select 2 datasets to use as input to the BBH algorithm. The Closure algorithm seeds the clusters with BBH hits, and then adds all sequences that (i) have a hit and (ii) pass the similarity and coverage rule with every other sequence in the cluster. The Best Hit option can form clusters on hitID or description substring; the similarity and coverage parameters are applied to both the hit and at least one sequence pair of the cluster. OrthoMCL builds a similarity matrix to normalize by species and uses Markov clustering.

#### 2.5.2 Cluster annotation and statistics

After creating clusters, runMultiTCW annotates the clusters with the majority hit; that is, it finds the common description substring among the best annotation hits of the sequences of the cluster and then finds the most common hit identifier for that description. Each cluster is assigned a %Hit, which is the percent of sequences in a cluster that have the description substring in any of its assigned hit descriptions.

The sequences of the clusters are aligned using the MSA program MAFFT [15] for N > 2 and the built-in dynamic programming alignment for N = 2. Each cluster is scored by sum-of-pairs and Wentropy [24]. Wentropy has been implemented within TCW, where it was translated from the mStatX program [25]. The user can request other scores from the mStatX program be computed in place of the sum-of-pairs and Wentropy.

#### 2.5.3 Pair annotation and statistics

First, given that the input sTCWdbs have the same conditions, the Pearson Correlation Coefficient (PCC) is calculated on the TPM values between each pair of sequences. Second, runMultiTCW provides statistics for the pairs found in clusters that have a hit. Each pair is aligned using a dynamic programming algorithm of the two translated ORFs and then maps the results to the corresponding codon-based ORFs, resulting in a codon-based alignment; see Wernersson and Pedersen [26] for a discussion on why this is important. The statistics detailed in Table 2 are computed from each pair alignment; the description column of the table states whether gaps are included, but no statistics include the overhangs (illustrated in Figure 2). The aligned pairs are written to file and a shell script is written for the user to run the KaKs_Calculator on the pairs. The shell script specifies the method name YN [27]; the reason for making YN the default is strictly because it is the fastest. By editing the shell script, the user can change to another method provided by the KaKs_Calculator, or provide results from alternative software. The Ka/Ks results are read into the mTCWdb.

**Table 2.**
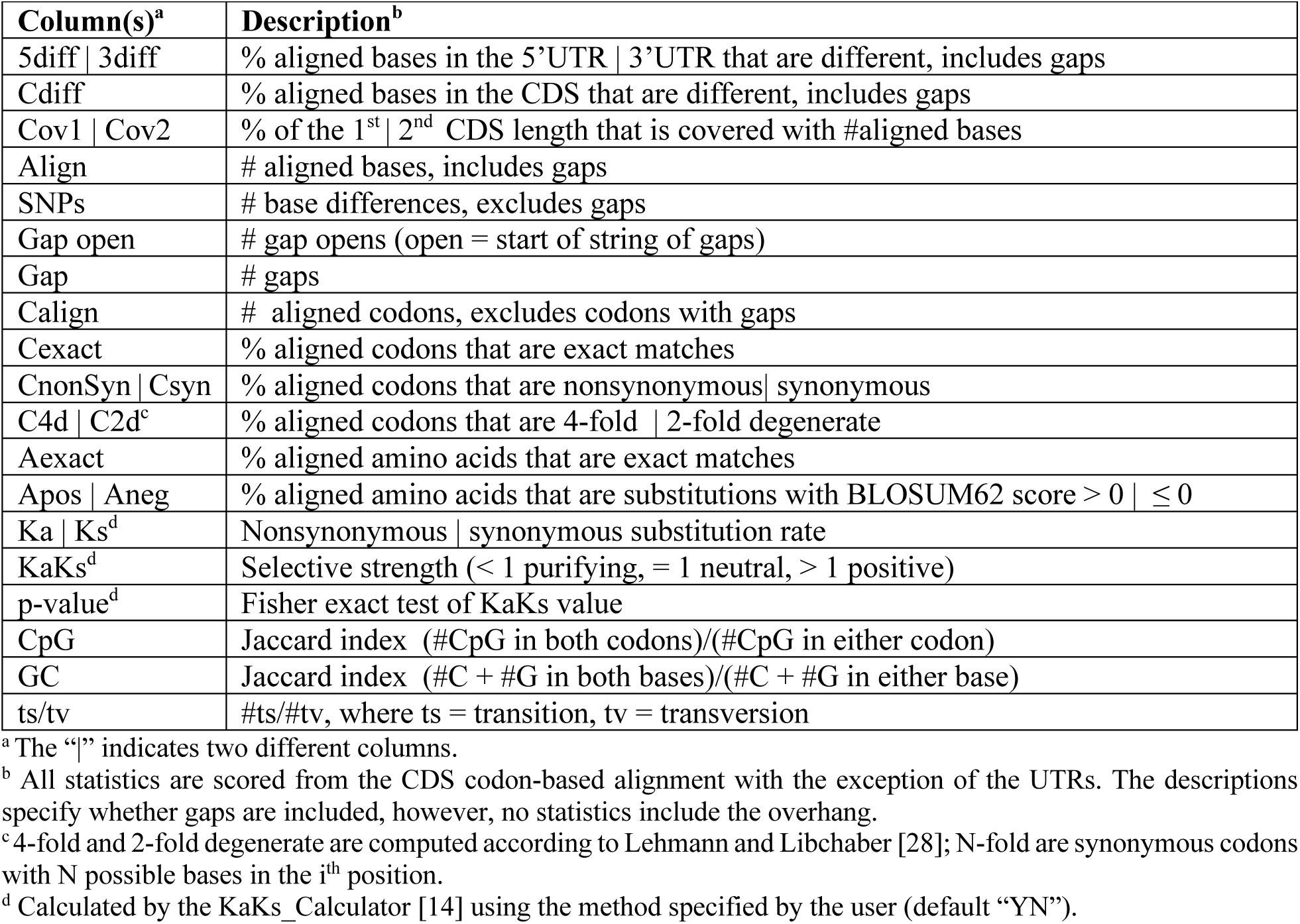
Pair statistics. The following statistics are computed for each pair that is in a cluster.

### 2.6 Query and display (viewSingleTCW and viewMultiTCW)

ViewSingleTCW provides filters and displays of the sTCWdb content. Briefly, the user can filter on the data associated with the sequences (e.g. Best Bits, DE, etc.), which results in a table of sequences. The hits can be filtered, which results in a table of hits. The GOs can be filtered, which results in a table of GOs; the relations between the GOs, hits and sequences are complicated; to aid in understanding the relations, there are multiple ways to view these relations. For both the hit and GO tables, the associated sequences can be view in the sequence table. From the sequence table, a sequence can be selected to show all information associated with it along with panels of the frame, GOs, and aligned hits. S1 and S2 Suppl provide snapshots of most of these tables, filters and panels.

ViewMultiTCW provides filters and displays of the mTCWdb content. Briefly, the user can filter on sequences, pairs, clusters and hits which results in a table of the corresponding results. From the cluster table, a selected cluster can be viewed by its sequences, pairs or MSA. From the pairs table, the selected pair can be viewed by its sequences, clusters or pairwise alignment. From the sequence table, multiple selected sequences can be viewed by their clusters, pairs or MSA, or a single selected sequence can display its details. From the sequence detail panel, the frame, GOs, or pairwise alignment of it hits and pairs can be viewed. The pairwise alignment can be by nucleotide (full sequence, CDS, 5’UTR, 3’UTR) or amino acid.

From the CDS alignment panel, the text alignment can be viewed with any of the following annotations: (i) synonymous/nonsynymous, (ii) amino acid, (iii) degenerate, (iv) CpG, or (v) ts/tv. S1 and S3 Suppl provide snapshots of most of these tables, filters and panels.

Both viewSingleTCW and viewMultiTCW produce overviews of their results and processing information (e.g. the dates of the GO tables and UniProts used), which is the initial view when starting either program. For all tables of results described for both viewSingleTCW and viewMultiTCW, the user can select the columns to view, move columns, and sort columns. All tables provide statistics on the selected numeric columns. All tables can be copied and exported in various formats. TCW does not provide overview graphical plots since there are many existing programs to create graphs, e.g. the data can be exported to a tab-delimited file for input into Excel. Instead, TCW graphics specializes in the details of the results such as alignments.

## 3 Results

The results use real data, but the analysis is strictly to demonstrate the TCW capabilities and does not attempt to answer any biological questions. Detailed descriptions on how to reproduce the rhizome study results are given in the S1 Suppl.

### 3.1 Rhizome study databases

To demonstrate the TCW analysis, the software is applied to the transcriptomes of two rhizome and one non-rhizome plant species. The rhizome *O. longistaminata* is a published dataset that is de novo assembled from Illumina reads with 5 replicates from rhizome, root, stem and leaf [16]; for this example study, the number of sequences was reduced from 143k to 48k (see S1 Suppl §1). The rhizome *N. nucifera* and non-rhizome *O. sativa* transcripts were downloaded from NCBI where they both have genome sequences [29] and [30], respectively. The three corresponding databases were named sTCW_OlR (*O. longistaminata* Rhizome), sTCW_NnR (*N. nucifera* Rhizome) and sTCW_Osj (*O. sativa* Japonica).

The TCW databases were built on the Linux machine specified in §2.1. The UniProt databases were downloaded on 21-Dec-2021; the resulting annoDBs are shown in Figure 3, where SP-full_BFIPV is the full SwissProt minus the records from the 5 taxonomic databases. The downloaded go-basic.obo was dated Nov-2021. The runSingleTCW defaults were used for building all three sTCWdbs. The DE was computed for sTCW_OlR with the edgeR script and the CpM filtering defaults. The runMultiTCW program was run to build mTCW_pl from the three sTCWdbs. Clusters were created using Closure, OrthoMCL and BBH, where TCW defaults were used for all computations.

**Figure 3.**
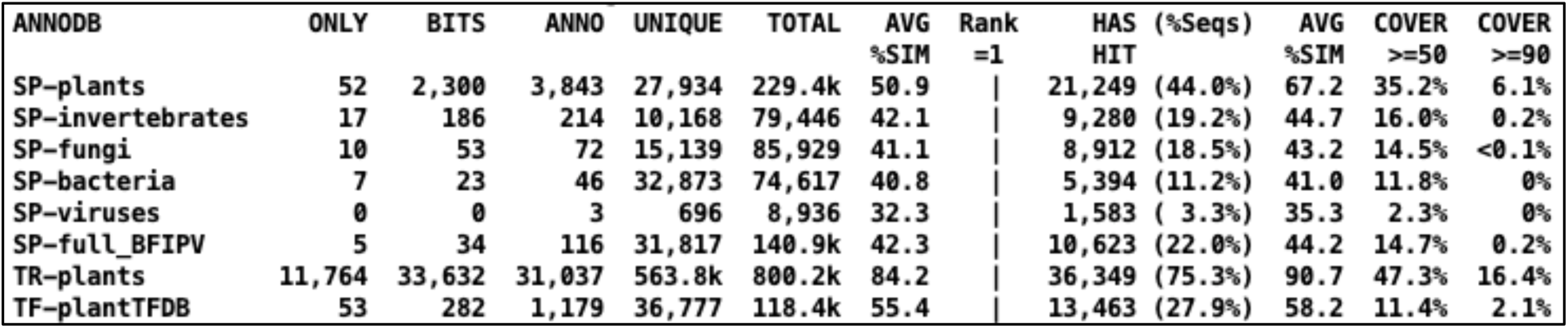
Overview of the annoDBs used to annotate the 48,272 OlR transcripts. The following describes the columns. ANNODB: the first part is defined by the input file, for example, “SP” and “TR” stand for SwissProt and TrEMBL, respectively, and “TF” refers to the transcription factor database PlantTFDB [31]; the second part is the taxonomy or source (named by the user). ONLY: the number of sequences that were only hit by the annoDB. BITS and ANNO: the number of sequences from the annoDB assigned the Best Bits and Best Anno, respectively. UNIQUE: the number of unique identifiers for the annoDB. TOTAL: the number of sequence-hit pairs from the annoDB. AVG %SIM: the average percent similarity for the total sequence-hit pairs. Rank=1: The four numbers following this heading refer to the best hit to the annoDB for each sequence. HAS HIT: the number and percent of sequences with at least one hit to the annoDB. AVG %SIM: the average percent similarity of the Rank=1 hits. COVER ≥ N: the percent of Rank=1 hits that have similarity ≥ N% and hit coverage ≥ N%.

The singleTCW results below use unpruned hits. However, when the OlR hits were pruned by alignment, the results had 11% reduction in sequence hits and 12% reduction in unique hits. When they were pruned by description, the results had 65% reduction in sequence hits and a 76% reduction in unique hits, yet all statistics shown in Figure 3 are similar except for the UNIQUE and TOTAL for the total hits (see S2 Suppl §2.1.2 for more details).

### 3.2 Transcriptome analysis for rhizome study

Table 3 provides summary statistics of the three datasets. Since *N. nucifera* (NnR) and *O. sativa* (Osj) are transcripts from genome sequence, it is expected that they would have transcripts that align well with the annoDB hits. Indeed, NnR has 99.2% and Osj has 99.0 transcripts with hits. *O. longistaminata* (OlR) had only 75.6% hit transcripts even though it is closely related to *O. sativa;* the lower number is partially due to them being assembled transcripts. Considering the species with the most hits, the top 8 species for the two Oryza databases were to Oryza species whereas none of the NnR top 8 species were to Oryza species. All three sTCWdbs had Best Bits hits to the transcription factor sequences in PlantTFDB, where OlR had 282, NnR had 118, and Osj had 222.

**Table 3.**
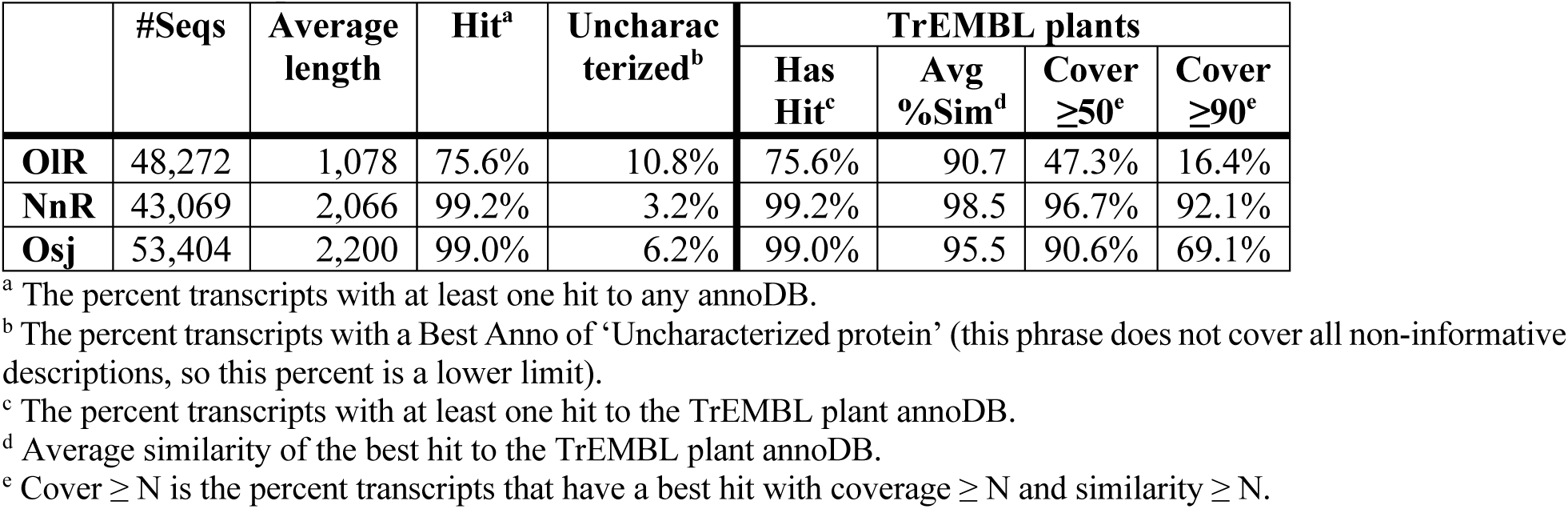
Transcript and hit statistics.

|  | #Seqs | Average length | Hit<sup>a</sup> | Uncharacterized<sup>b</sup> | TrEMBL plants |  |  |  |
| --- | --- | --- | --- | --- | --- | --- | --- | --- |
|  |  |  |  |  | Has Hit<sup>c</sup> | Avg %Sim<sup>d</sup> | Cover ≥50<sup>e</sup> | Cover ≥90<sup>e</sup> |
| **OIR** | 48,272 | 1,078 | 75.6% | 10.8% | 75.6% | 90.7 | 47.3% | 16.4% |
| **NnR** | 43,069 | 2,066 | 99.2% | 3.2% | 99.2% | 98.5 | 96.7% | 92.1% |
| **Osj** | 53,404 | 2,200 | 99.0% | 6.2% | 99.0% | 95.5 | 90.6% | 69.1% |

<sup>a</sup> The percent transcripts with at least one hit to any annoDB.
<sup>b</sup> The percent transcripts with a Best Anno of 'Uncharacterized protein' (this phrase does not cover all non-informative descriptions, so this percent is a lower limit).
<sup>c</sup> The percent transcripts with at least one hit to the TrEMBL plant annoDB.
<sup>d</sup> Average similarity of the best hit to the TrEMBL plant annoDB.
<sup>e</sup> Cover ≥ N is the percent transcripts that have a best hit with coverage ≥ N and similarity ≥ N.

#### 3.2.1 Open reading frames

The average length of the ORFs for OlR, NnR and Osj was 578, 1375 and 1269 nucleotides, respectively. Table 4 provides a summary of their features.

**Table 4.** Summary of ORFs.

| Percent of transcripts | OIR | NnR | Osj | Exact hit<sup>a</sup> | OIR | NnR | Osj |
| --- | --- | --- | --- | --- | --- | --- | --- |
| %Hit & Longest & Markov | 43. | 86.8 | 73.4 | % Sim ≥ 90 | 92.9 | 99.9 | 98.2 |
| % Has hit | 75.6 | 99.2 | 99.0 | % Longest & Markov | 73.7 | 90.6 | 80.2 |
| % Longest ORF | 61.0 | 93.2 | 79.1 | % Longest ORF | 76.1 | 96.6 | 82.1 |
| % Markov best score | 71.5 | 89.8 | 86.4 | % Markov best score | 92.8 | 92.2 | 94.8 |
| % ORF ≥ 300 | 50.1 | 92.8 | 91.2 | % ORF ≥ 300 | 90.3 | 96.7 | 96.9 |
| % Exact hits<sup>a</sup> | 7.1 | 86.6 | 54.2 |  |  |  |  |

<sup>a</sup> Exact Hit: An alignment to the entire protein ending with a ATG and stop codon; there may be mismatches or gaps (e.g. the hits in Figure 2 are exact). The exact hit counts are OIR = 3446, NnR = 37,294, Osj = 28,922.

For the transcripts with genome sequence, the percentage of all transcripts with exact hits for NnR was 86.6% and Osj was 54.2%. When the Osj ORFs were computed allowing alternative start sites, the percentage of exact hits only increased to 54.7%. Computing the longest ORF often finds the correct ORF, but not always; for example, considering the NnR 37,294 exact hits, 96.6% were the longest. The Markov score [22] is a excellent indication of coding, where 92.2% of the NnR exact hit ORFs had the best score. Putting the two features together, 90.6% were the longest with the best Markov score. The Osj had a much lower percentage for the longest (82.1%) and higher percentage for the Markov score (94.8%).

ORF finding is complicated when there are multiple hit frames or stop codons within the hit region; the percentages of these two cases (multi-frame, stop-in-hit) was OlR (15.7%, 9.7%), NnR (8.1%, 5.4%) and Osj (24.3%, 14.7%). The sequences with multiple frames and stop codons within hits can be viewed in viewSingleTCW, which is demostrated in S2 Suppl §2.1.3.

##### Comparison with TransDecoder v5.5.0

The TransDecoder software [22] is often used for finding ORFs, which also uses information from hits, length and the Markov score. S2 Suppl §2.2.1 compares results between TCW and TransDecoder (TD) for the task of finding the best ORF per sequence, where the results are summarized as follows: 15,000 sequences with a perfect hit (exact hit with 100% similarity) to TR-plants were used, of which 9000 had hits to SP-plants. The TCW and TD ORFs were computed using the SP-plants hits and the results compared to the TR-plant perfect hits. TCW produced an ORF for all sequences and TD produced results for all but 192 sequences. Of the 14,808 that had a TD results: TCW had 35 and TD had 50 wrong frames, TCW had 9 and TD had 3 non-overlapping coordinates, TCW had 639 and TD had 2022 wrong starts. The greatest difference was with the start coordinate, where TCW performs much better because it directly uses the start coordinates of the SP-plant hit when it is a good hit.

#### 3.2.2 Differential expression

To explore the differential expression, the OlR database was queried for transcripts that were preferentially expressed in the rhizome compared to the other tissues. Comparing rhizomes to root, stem and leaf using p-value < 1E-04, there were 483 down-regulated transcripts of which 21 had TPM ≥ 50 for rhizomes. There were 863 up-regulated transcripts of which 151 had TPM ≥ 50 for rhizome. Of the 151 transcripts, 9 were not annotated, which are shown in Figure 4.

**Figure 4.**
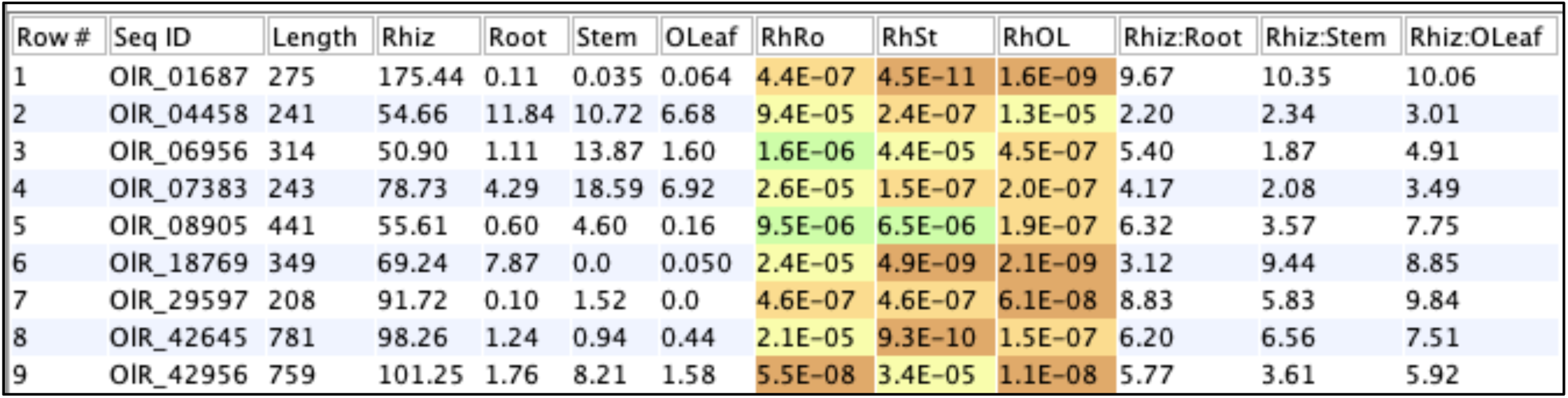
Unique rhizome transcripts that are up-regulated for rhizomes. Transcripts with no annotation, rhizome TPM ≥ 50 and up-regulated for rhizome compared to root, stem and leaf at p-value < 1E-04. The columns Rhiz, Root, Stem and OLeaf are the TPM values, the middle three are the DE columns, and the last three columns are the fold-change.

#### 3.2.3 Gene ontology

To explore the gene ontology, the OlR database was queried on Plant GO Slims, which resulted in 97 GO terms. This set was analyzed for enrichment of rhizome to root (RhRo), stem (RhSt) or leaf (RhOL), where 32 GO terms were enriched using GOseq p-value < 1E-06; the graph is shown in Figure 5.

**Figure 5.**
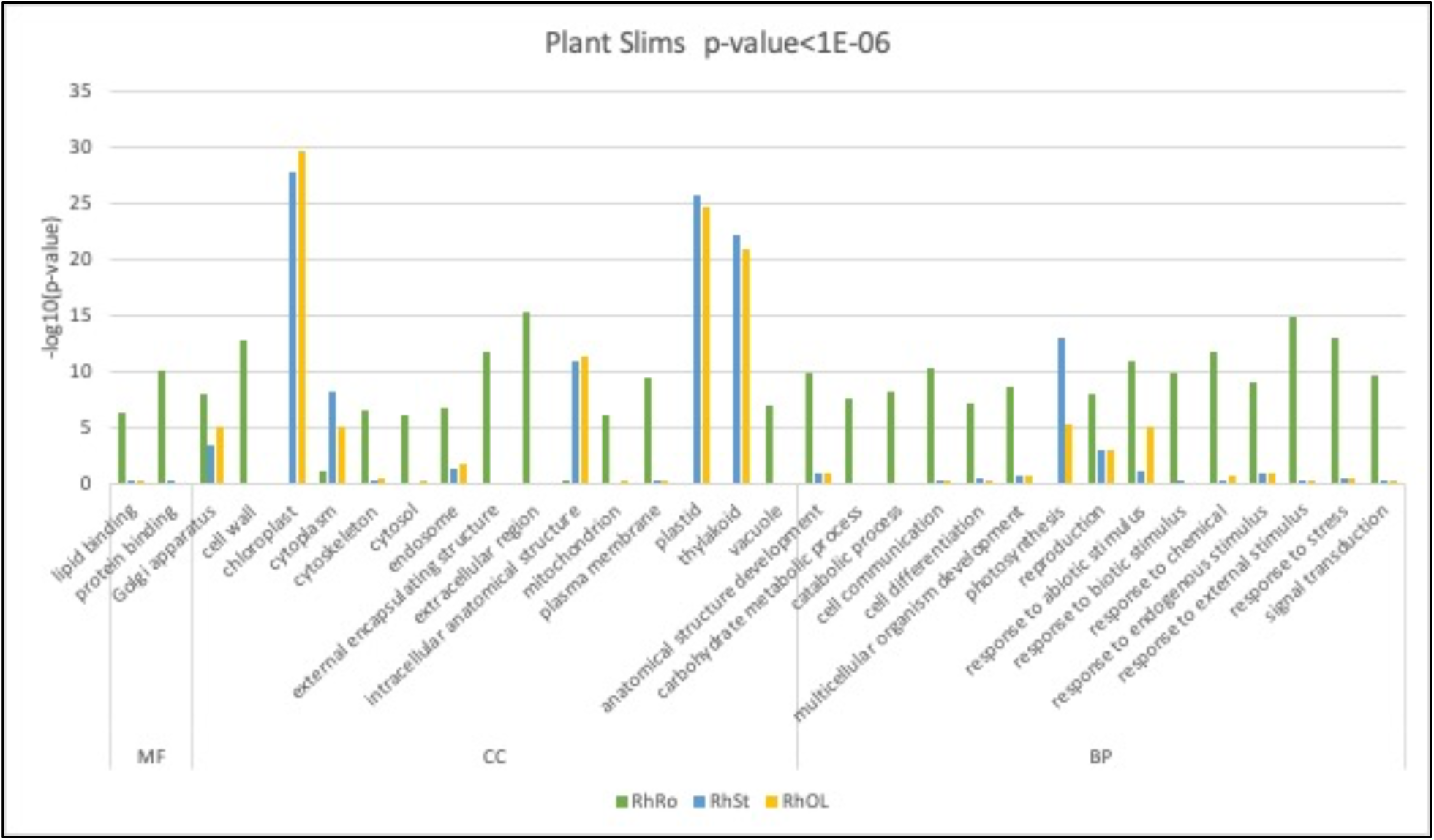
GO enrichment of Plant GO Slims for OlR rhizomes. Rhizomes were compared to root (RhRo), stem (RhSt) and leaf (RhOL) using p-value < 1E-06 for any of the three.

The following demonstrates using TCW results with other programs. There were 162 total GOs enriched for rhizomes at p-value < 1E-06. The GO terms were from levels 2-10, with 100 biological process, 57 cellular component and 5 molecular function. The data was input into REVIGO [32] where the values associated with each GO term were the number of DE transcripts; the 57 cellular component GOs were reduced to 30 and shown graphically (see S2 Suppl §4). WEGO [33] reduced the entire set to 29 GO terms from WEGO’s level 3 and shown graphically (see S2 Suppl §4).

#### 3.2.4 Timing results

The following execution times are from the Linux machine described in §2.1. It took 7 minutes to build the sTCW_OlR database of 48,272 sequences with 20 replicate counts per sequence. Table 5 shows the annotation times, where the second row has 4 additional TrEMBL taxonomic database (invertebrate, fungi, virus, bacteria). The execution time for row 1 is further broken down in S1 Suppl §3.2 along with times for description pruning annotation (Total 2h:09m on Linux and 51m on MacOS).

**Table 5.** SingleTCW annotation times on Linux for 48,272 sequences.

| Row | AnnoDBs<sup>a</sup> | Largest AnnoDB | runAS<sup>b</sup> | runSingleTCW Annotate |  |  |
| --- | --- | --- | --- | --- | --- | --- |
|  |  |  |  | Add Anno<sup>c</sup> | Add GO | Total<sup>d</sup> |
| 1 | 8: 6 SP, 1 TR, 1 Tf | 5.1Gb TR-plants | 1h:17m | 1h:06m | 1h:03m | 2h:28m |
| 2 | 12: 6 SP, 5 TR, 1 Tf | 32.5Gb TR-bacteria | 22h:23m<sup>e</sup> | 4h:22m | 2h:02m | 6h:37m |

<sup>a</sup> SwissProt (SP), TrEMBL (TR), and PlantTFDB (Tf)
<sup>b</sup> Includes download times.
<sup>c</sup> Includes search times, but not the times to build the DIAMOND databases.
<sup>d</sup> Includes Add Anno, Add GO and ORF-finding.
<sup>e</sup> 17h:20m was for TR-bacteria (6h:36m download, 2h:21m create FASTA, 8h:23m load GObd)

There were 75.6% annotated sequences using the 8 annoDBs and 75.8% using the 12 annoDBs. Given that the difference in annotated sequences is small, it is generally not worth including the large TrEMBL databases except for the most relevant.

### 3.3 Comparative analysis for rhizome study

The mTCW_pl comparative database was created from sTCW_OlR, sTCW_NnR, and sTCW_Osj. Table 6 displays the number of clusters for BBH, Closure and OrthoMCL.

**Table 6.** Summary of clusters.

| Method<sup>a</sup> | Clusters |  |  |  |  |  |  | Cluster pairs<sup>d</sup> |  |
| --- | --- | --- | --- | --- | --- | --- | --- | --- | --- |
|  | =2 | 3-5 | 6-15 | >15 | #Seqs | Shared Desc<sup>b</sup> | Avg Went<sup>c</sup> | Avg Ka | Avg Ks |
| **NnR-OIR (Bnl)** | 5,056 | 0 | 0 | 0 | 7.0% | 90.5% | 0.58 | 0.188 | 6.928 |
| **NnR-Osj (Bns)** | 6,473 | 0 | 0 | 0 | 8.9% | 91.1% | 0.65 | 0.198 | 7.009 |
| **OIR-Osj (Bls)** | 13,424 | 0 | 0 | 0 | 18.5% | 96.4% | 0.77 | 0.024 | 0.271 |
| **CL** | 11,770 | 10,322 | 4,181 | 284 | 70.2% | 90.0% | 0.76 | 0.125 | 4.118 |
| **OM** | 5,692 | 7,936 | 5,509 | 1,217 | 80.0% | 80.0% | 0.69 | 0.260 | 6.044 |

<sup>a</sup> The first three methods are BBH; CL is Closure; OM is OrthoMCL.
<sup>b</sup> The percentage of the clusters where all sequence share an annotation description.
<sup>c</sup> Wentrophy scores are between 0 and 1 where 1 is the most conserved.
<sup>d</sup> Average Ka (nonsynonymous rate) and Ks (synonymous rate).

CL has more total clusters than OM (26,557 versus 20,558), yet it uses less sequences (70.2% versus 80.0%). As shown in Table 6, CL has a higher number of clusters where all sequences share an annotation hit compared to OM, higher Wentrophy and lower Ka and Ks. This would imply that the CL clusters are more conservative. If OM was run with the less stringent inflation of 1.5, the differences would be greater.

TCW specializes in exploratory graphics, where the scientist can view the details of a cluster. For example, if the scientist was interested in the “HORMA domain-containing protein”, Figure 6A shows that the two Orzya species have perfectly aligned amino acid sequences and NnR has some substitutions. Figure 6B shows that the two Oryza sequences have four mismatches for the nucleotide sequence that result in synonymous codons, as shown in Figure 6C.

**Figure 6.**
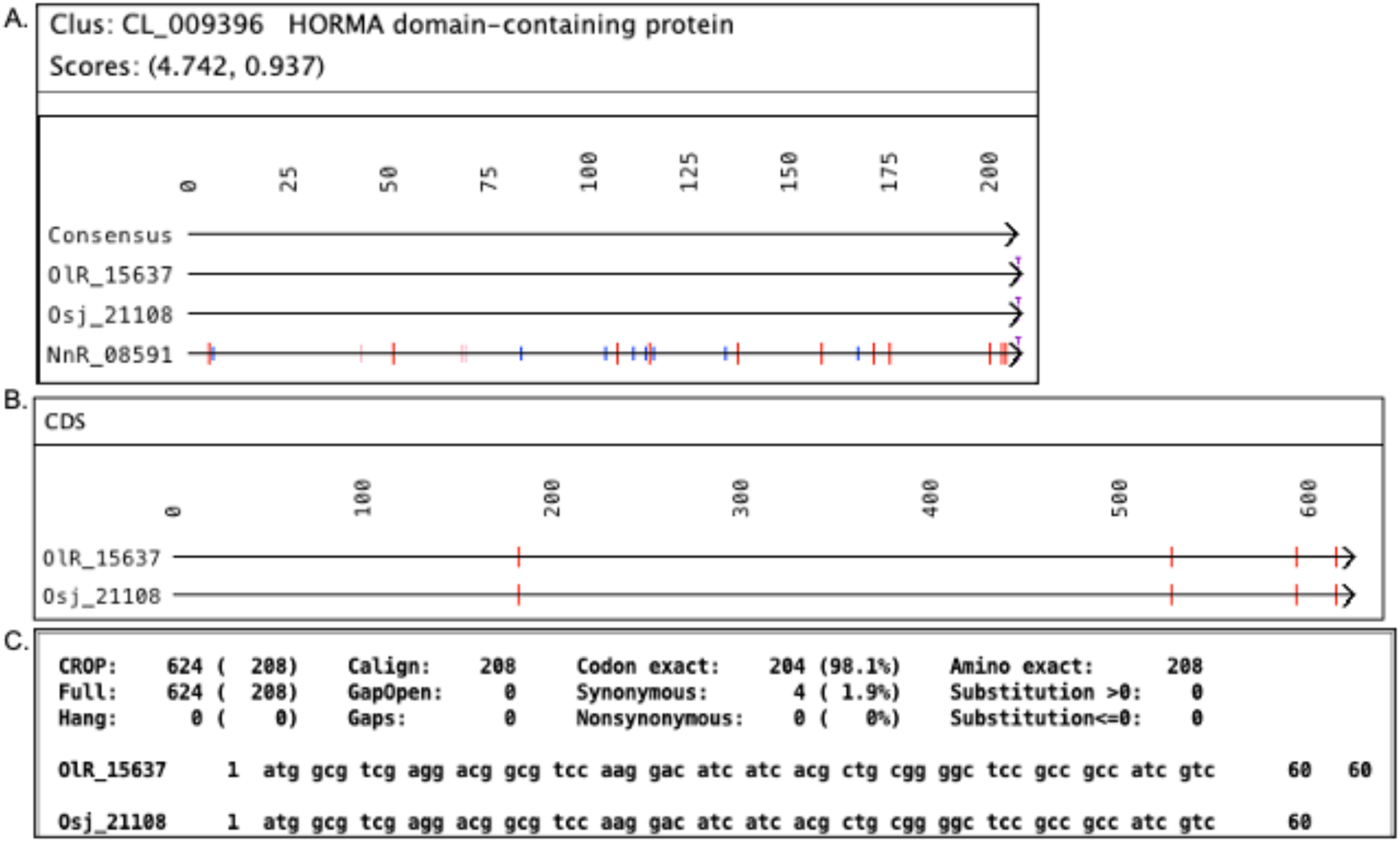
MSA and Pairwise alignments. (A) The MSA of a cluster where the first two amino acid sequences are identical, the third one has some differences. The green marks are gaps, and the rest of the marks indicate a difference in amino acid, where blue marks are BLOSUM62 > 0, pink are BLOSUM62 = 0, red are BLOSUM62 < 0. (B) For the identical amino acid sequences, the CDS nucleotide alignment shows 4 mismatches (red marks). (C) The top portion of the CDS text alignment states that the mismatches correspond to synonymous codons (the 4 mismatches are shown on the full text alignment).

#### 3.3.1 Rhizome specific clusters

Clusters with at least one transcript from the two rhizome datasets OlR and NnR, and no transcripts from the non-rhizome Osj dataset are potential rhizome-specific proteins. From the Closure clusters, there were 243 candidate rhizome clusters where all of them were annotated, one had the description “uncharacterized” and 201 (82%) shared the same annotation description. Figure 7 shows the top 10 clusters sorted by Wentrophy (Score2) score. From this cluster set, NnR had an average of 1.64 transcripts per cluster and OlR has 1.11 transcripts per cluster. Further analysis of these clusters would be of interest to a scientist studying rhizomes, where the descriptions and GO assignments could elucidate important functionality of rhizomes.

**Figure 7.**
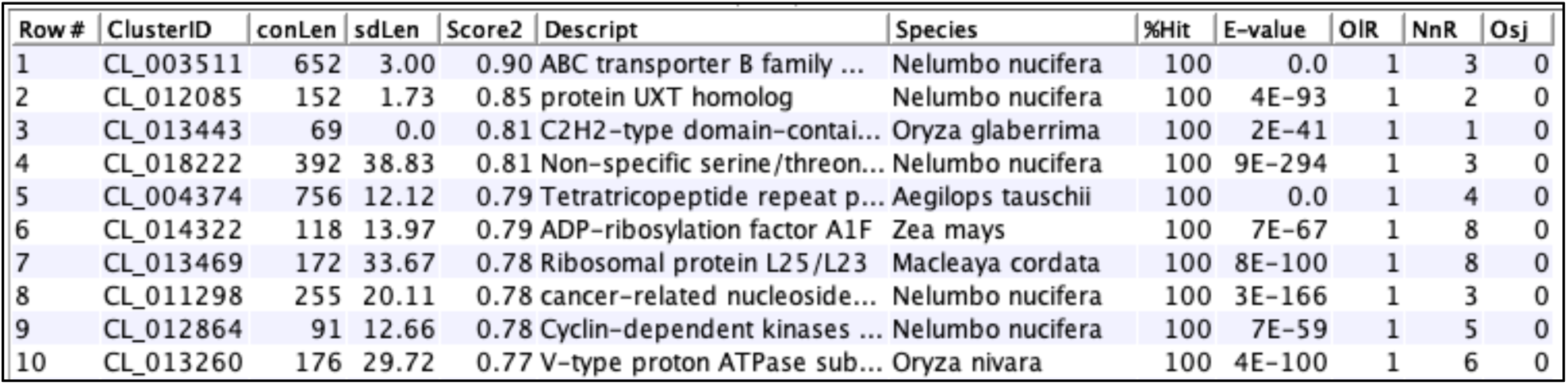
Top ten annotated rhizome Closure cluster. Only clusters where all members have the same shared annotation description (%Hit>=100) are shown, sorted on Score2. The conLen column is the consensus length and sdLen is the standard deviation of the aligned sequences lengths. Score2 is Wentrophy. The E-value is from the best hit. The OlR, NnR and Osj columns are the number of transcripts from each respective dataset.

#### 3.3.2 BBH clusters

Figure 8 shows the BBH statistics from the related OlR-Osj pairs and the more distantly related OlR-NnR. The 94.9% exact codons for OlR-Osj elucidates how closely related they are compared to the 33.9% for OlR-NnR. The BBH NnR-Osj statistics (not shown) have very similar values to the NnR-OlR.

**Figure 8.**
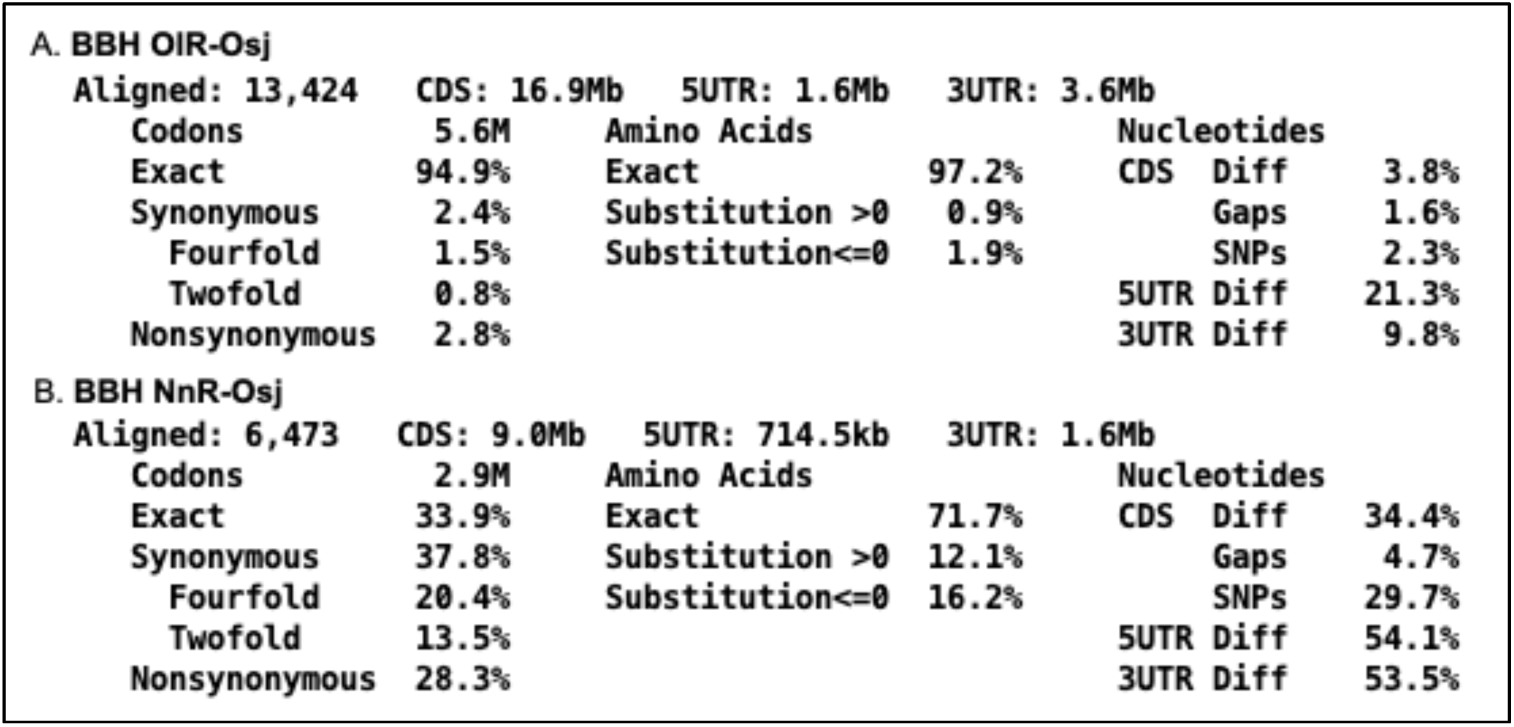
Pair statistics for BBH OlR-Osj and NnR-Osj. The explanation of each statistic is given in Table 2.

##### Comparison with Galaxy BBH

S3 Suppl §4.1.1 has a comparison between TCW and Galaxy BBH, summarized as follows. The following are handled differently in the two programs: (i) filter with or without rounding, (ii) filter before or after creating possible BBH pairs, (iii) handling of a tie between two pairs, (iv) how the self-search is performed. TCW and Galaxy agreed on 800 pairs, where TCW had 853 and Galaxy had 850 pairs. The difference are typically from the situation where A’s best hit is B, whose best hit is C, who has a bi-directional hit with D; the differences in handling can change this relation. The benefit of the Closure method is that it brings ABCD into the same cluster.

#### 3.3.3 Ka/Ks results

Figure 9A shows the Ka/Ks summary for the BBH OlR-Osj pairs. As stated by Zhang et al. [14], the selection is neutral if Ka = Ka, purifying if Ka < Ks, and positive (diversifying) if Ka > Ks. Either Ka or Ks may be NA (if Ka = NA, then KaKs = NA; if Ks = NA, then KaKs = 0). Since it is rare for Ka to be exactly equal to Ks (mTCW_pl has 1pair), the TCW summary uses the following: Ka/Ks ∼ 1 is (0.995 ≤ KaKs < 1.006), Ka/Ks < 1 is (KaKs < 0.995) and Ka/Ks > 1 is (KaKs ≥ 1.006).

**Figure 9.**
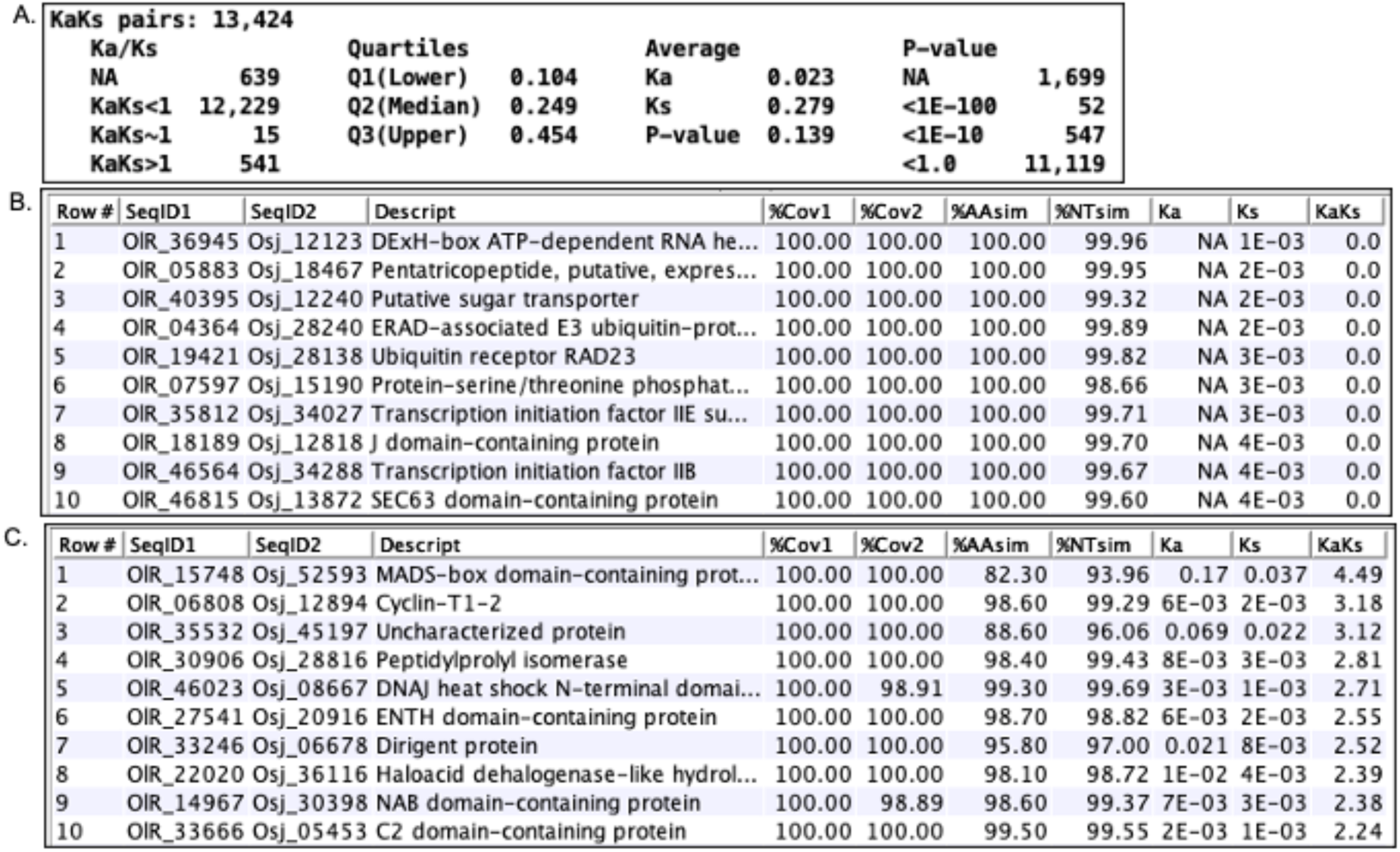
Ka/Ks for BBH OlR-Osj top purified and diversified pairs. (A) The overview for the Ka/Ks results for the BBH OlR-Osj pairs. (B) Purifying: The ten pairs with the lowest Ks scores, coverage ≥ 98% and no gaps. (C) Diversifying: The ten pairs with the highest Ka/Ks scores, coverage ≥ 98% and no gaps.

Filtering on coverage ≥ 98% for both sequences, no gaps and Ka/Ks < 1 results in 1964 pairs, where Figure 9B shows the the top 10 with the lowest Ks value. Using the same filter but with KaKs > 1 results in 73 pairs, where Figure 9C shows the top ten.

#### 3.3.4 Timing results

Table 7 shows the execution time on the Linux machine to build two multiTCW databases. The mTCW_rhi has two input sTCWdbs that were description pruned. S1 Suppl §3.3 shows a breakdown of times for the different processing steps on Linux and Mac.

**Table 7.**
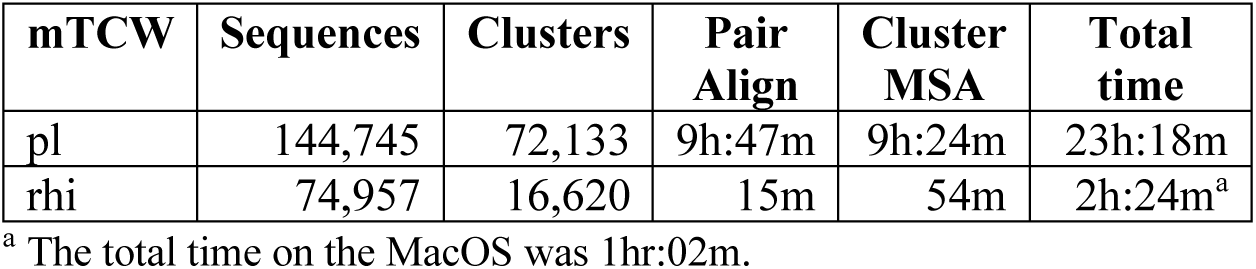
MultiTCW build times on Linux.

| mTCW | Sequences | Clusters | Pair Align | Cluster MSA | Total time |
| --- | --- | --- | --- | --- | --- |
| pl | 144,745 | 72,133 | 9h:47m | 9h:24m | 23h:18m |
| rhi | 74,957 | 16,620 | 15m | 54m | 2h:24m<sup>a</sup> |

<sup>a</sup> The total time on the MacOS was 1hr:02m.

The step to add GOs can be omitted if they are not going to be used (viewMultiTCW has limited support for GOs); the time to add GOs for mTCW_pl was 1h:36m, which is part of the total time in Table 7. The MSA step can be omitted if it is sufficient to compute the MSA and scores per cluster on-the-fly in viewSingleTCW, but it is generally worth taking the compute time to have them pre-computed.

## 4 Discussion

The topic of reproducibility has been addressed in numerous publications. Peng [34] discusses how the introduction of computers to the analysis of biological data has introduced published computational results that are not reproducible. Stodden et al. [35] reviewed 204 *Science* publications and were able to reproduce the findings of only 26% of them. Garijo [36] provided reproducibility guidelines for authors, one of which was the importance of using published open source software whenever possible. TCW provides easily reproducible results and is open source. The supplements provide instructions on how to reproduce the results in the manuscript and the supplements. A significant benefit is that researchers can produce the same type of TCW results for their data; this saves time in figuring out how to generate the results and for writing detailed methods on how they were produced.

The topic of reproducibility has drawn attention, but there are also the problems of loss of results, extensibility and accessibility. Bioinformatics often utilize many flat files (plain text files such as a FASTA file) of data and results, and it is complicated to keep such files organized. As discussed, TCW stores all data in a MySQL database and allows the input of external results, hence, keeping all results in one place and is easily accessible to all members of the team.

Scientists explore transcriptomes through ingenious wet lab experiments; however, with large-scale data they must also be proficient at exploring the transcriptomes through computational approaches. The scientist should be able to have at their workbench a computer with multiple applications to aid in exploring this wealth of information. Since computational analysis is permanently part of the biology world, it should be routine for any lab that analyzes large-scale data to have a bioinformatics scientist as a member of the team, who can install the basic dependencies used by software such as TCW.

TCW allows the viewing of all data and results, nothing is hidden, and the graphics allow the user to verify and understand the results. Some examples are as follows: Using viewSingleTCW, the user can view over-expressed GOs for a given p-value, and can then drill down to view all sequences that have a hit with a given GO term. A hit E-value of 1E-100 could be a long match with mismatches and gaps or a short exact match, where observing the actual TCW alignment aids in understanding the characteristics of the match. With viewMultiTCW, the user can view the text alignment of a homologous pair along with the location of the different statistics listed in Table 2. In summary, allowing the scientist to view the details of the computation aids in clearly understanding the results. It identifies the ambiguities and difficulties and demonstrates why there is not a perfect algorithm to solve many of these problems. This in turn could lead to refinement of the wet lab experiments.

TCW includes many different types of computations, and the objective was to use the most state-of-the-art approach for each computation. New methods and algorithms will be added in the future along with enhanced graphics and queries. Computer scientists, biologists and bioinformaticians are welcome to directly add or request help in adding their methods to TCW.

## 5 Supporting information

### S1 Suppl. Datasets, reproduce results, timings

§1. Details of the datasets used in the manuscript and supplements. §2. Instructions on how to reproduce the single and multi TCW results along with TCW snapshots. §3. Timing results for both single and multi TCW on a Linux and Mac machine. §4. External software and databases used by TCW.

### S2 Suppl. Build a singleTCW database

§1. Using runAS for downloading UniProts and GO for TCW annotation. §2. Using runSingleTCW, which includes *N. nucifera* multi-frame hits and hits with stop codons, the ORF finding algorithm with comparison to TransDecoder, GO levels, adding external data. §3. RunDE for differential expression. §4. GO results using REVIGO and WEGO.

### S3 Suppl. Build a multiTCW database

§1. The runMultiTCW interface. §2. Building the database and adding GOs. §3. Search parameters and adding pairs. §4. Computing clusters, the BBH algorithm with comparison to Galaxy BBH, assigning annotation to cluster. §5. Run Stats: statistics for pairs and clusters. §6. GC, CpG, Ts/Tv computations. §7. TPM, DE and PCC.

## 6 Acknowledgments

The 24-CPU Linux machine used for building the databases is housed at the BIO5 Institute and Lomax Boyd of BIO5 provided the system maintenance.

