## Supplementary material for "Transcriptome computational workbench (TCW): analysis of single and comparative transcriptomes": S1 Suppl

### Datasets, reproduce results, timings

**Sections**

1 Datasets ... 1
2 Reproduce manuscript results ... 2
2.1 SingleTCW results ... 2
2.1.1 Figure 2. Alignment of Best Bits (bit-score) and Best Anno (annotation) ... 2
2.1.2 Figure 3. Overview of annoDBs ... 2
2.1.3 Table 3. Transcript and hit statistics ... 3
2.1.4 Table 4. Summary of ORFs; longest ORF; multi-frame and stop-in-hit results ... 3
2.1.5 Figure 4. Unique rhizome transcripts ... 4
2.1.6 Figure 5. GO enrichment of plant slims for OlR rhizomes ... 5
2.2 MultiTCW results ... 6
2.2.1 Table 6: Summary of clusters ... 6
2.2.2 Figure 6. MSA and Pairwise ... 7
2.2.3 Figure 7. Top ten annotated rhizome Closure clusters ... 8
2.2.4 Figure 8. Pair statistics for BBH OlR-Osj (Bls) and NnR-Osj (Bns) ... 10
2.2.5 Figure 9. Ka/Ks for BBH OlR-Osj, top 10 conserved/diverged pairs ... 10
3 Timing results of builds ... 10
3.1 Test machines ... 10
3.2 Build single database ... 10
3.3 Build multi database ... 11
3.4 Memory ... 12
3.5 Timing functions ... 12
4 External software and databases ... 13
5 References ... 13

### 1 Datasets

The following datasets were used for the results and supplements:
sTCW_OlR*: Oryza longistaminata* assembled transcripts and 5 replicates for rhizome, root, stem and oleaf [1].

- This dataset contained 143k transcripts. It was reduced to 48k by only using the ones that have TPM ≥ 3 for rhizome, root, stem or old leaf.
- The single replicate tip and zone samples were not used.
- The original dataset [1] was deposited to the Short Read Archive (SRA) in Genbank under BioProject accession PRJNA196977.

sTCW_Osj: *Oryza sativa* subsp. Japonica [2] from www.ncbi.nlm.nih.gov/genome/?term=Oryza+Sativa

- GCF_001433935.1_IRGSP-1.0_rna.fna.gz (June 2021)

sTCW_NnR*: Nelumbo nucifera* [3] from https://www.ncbi.nlm.nih.gov/genome/14095

- GCF_000365185.1_Chinese_Lotus_1.1_rna.fna.gz (June 2021)

The following were used for the supplements only:
sTCW_NNU*: N. nucifera* from publication [4]

- 26,685 sequences and 5 replicates for rhizome, root, stem and old leaf.

### 2 Reproduce manuscript results

The following subsections describe how the results in the manuscript were derived using TCW. References to viewSingleTCW and viewMultiTCW panels and options are in Arial Narrow text.

#### 2.1 SingleTCW results

##### 2.1.1 Figure 2. Alignment of Best Bits (bit-score) and Best Anno (annotation)

- Basic Sequences (Figure S1): Enter 04675 in the text box followed by BUILD. On the resulting table, select the sequence followed by Seq Detail.
- On the resulting panel (Figure S2), select Best Hits from Align Hits…

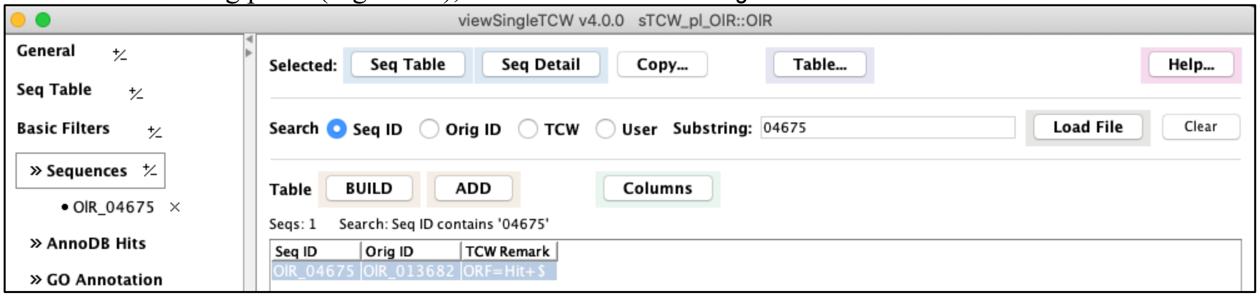


**Figure S1. Basic Sequences for finding a specific sequence.**

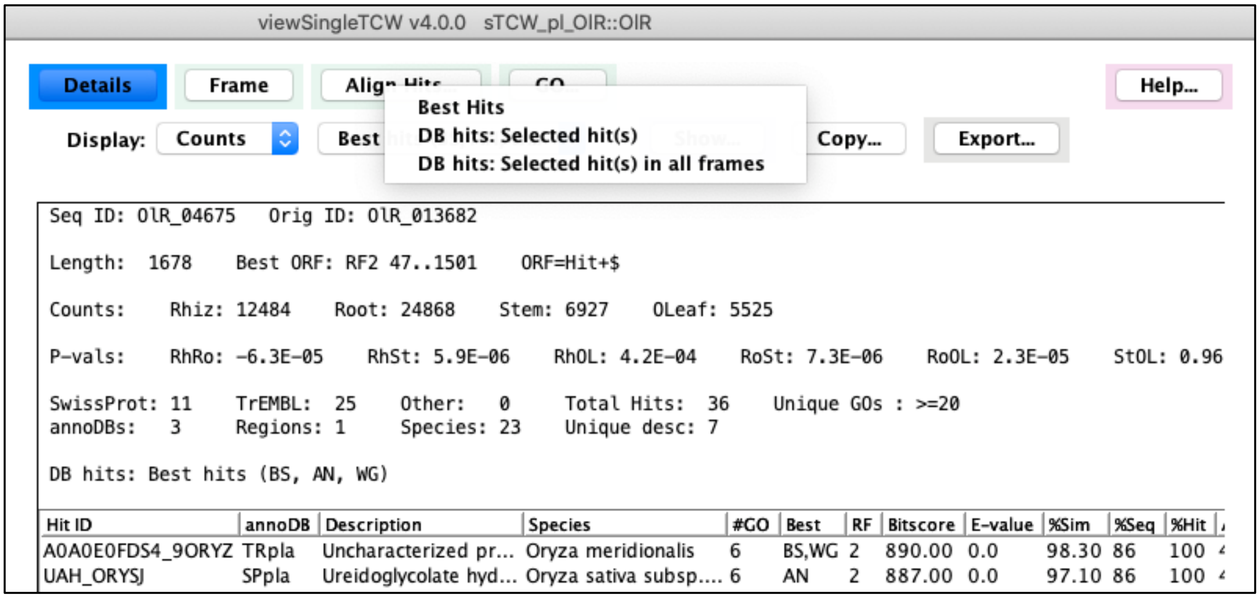


**Figure S2. Sequence Details of OIR_04675.**

##### 2.1.2 Figure 3. Overview of annoDBs

Figure 3 is a snapshot of the “annoDB” section from the sTCW overview, as shown in Figure S3. At the bottom of the Overview is a Reproduce button, which popups a panel that describes how to reproduce the numbers in the Overview.

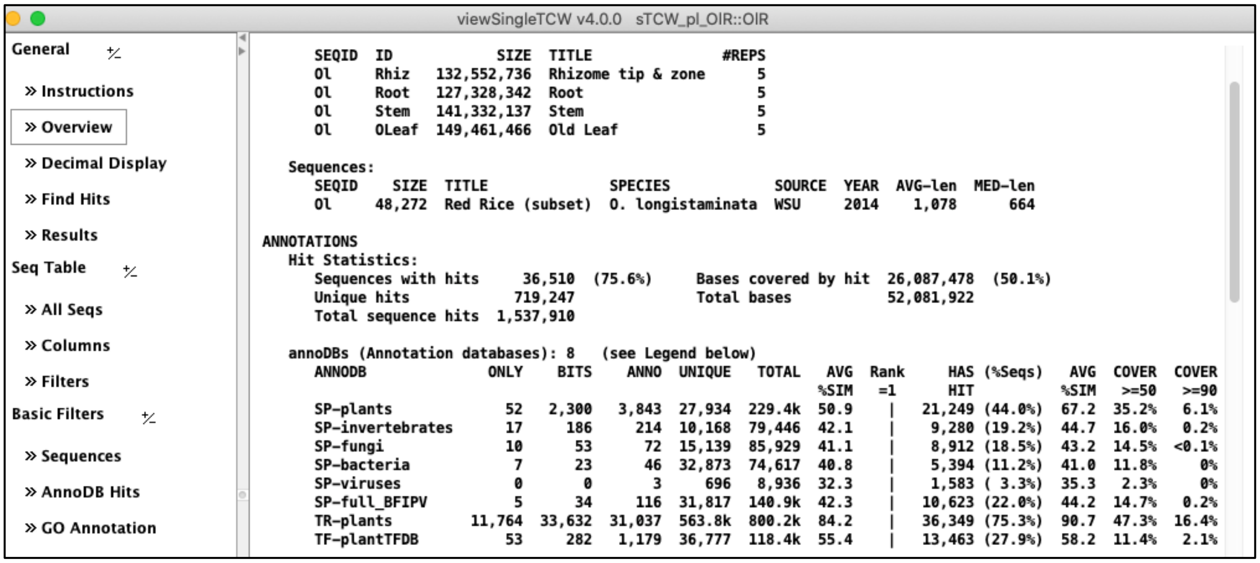

**Figure S3. SingleTCW overview showing the annoDBs.**

#### 2.1.3 Table 3. Transcript and hit statistics

- Columns 1-3: The values from the Overview (Figure S3 above) of SIZE, AVG-len and Sequences with hit.
- Columns 5-9: Last 4 columns of the second to last line of Figure S3.
- Column 4: For "Uncharacterized":
    a. Basic AnnoDB Hits (Figure S4): Set the filters as shown.
    b. Compute the percent of annotated hits that are "Unchar" by using the number beside the Seqs: (e.g. 3,930) over the Sequences with hits number in the Overview (e.g. 36,510).

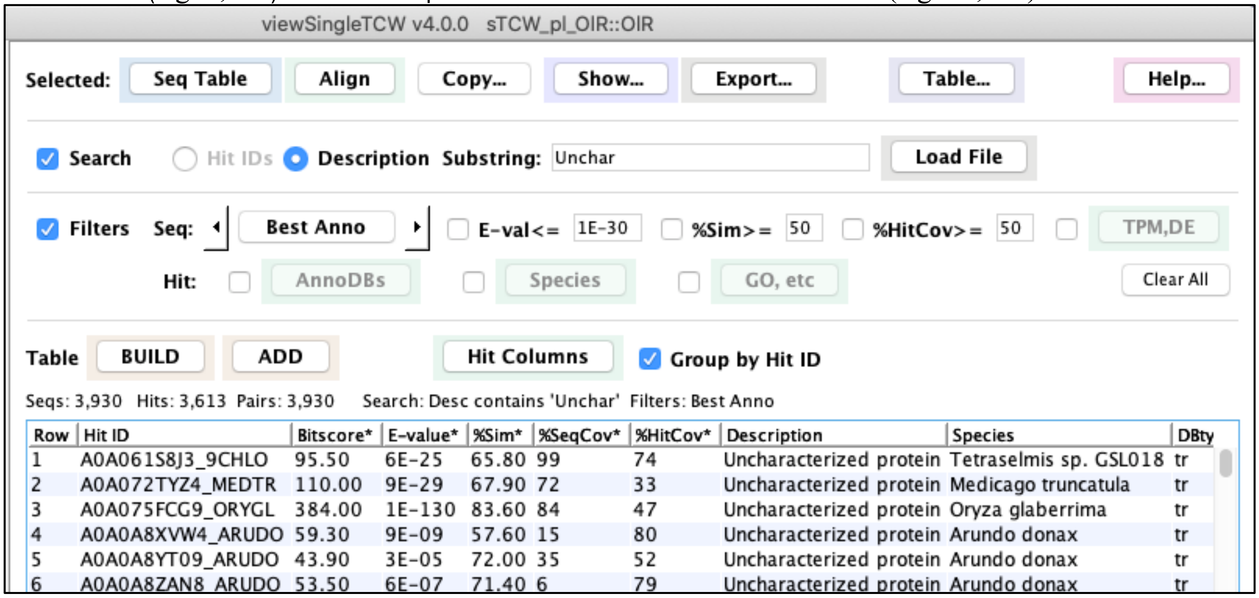

**Figure S4. Basic AnnoDB Hits for finding "Uncharacterized protein".**

#### 2.1.4 Table 4. Summary of ORFs; longest ORF; multi-frame and stop-in-hit results

All statistics for Table 4 were from the Overview and the ORF finder log file (projects/<project-name>/logs/anno.log), where the <project-name> was OlR, NnR and Osj (see S2 Suppl §2.2 Figure S12 for an example of the OlR anno.log file). The percent of sequences with multi-frame hits or stop codons in hit regions is also shown in this file and the Overview; see S2 Suppl §2.1.3 for an in-depth discussion on these sequences.

##### 2.1.5 Figure 4. Unique rhizome transcripts

- Sequence Filter (Figure S5): Set the filters as shown.
- The Filter panel has a button called Filtered Seq Table, which results in the Sequence Table (Figure S6).
- The column were set with the Columns tab (the tab is shown in Figure S1 above).
- The color scheme can be changed with the Decimal Display panel (Figure S6).

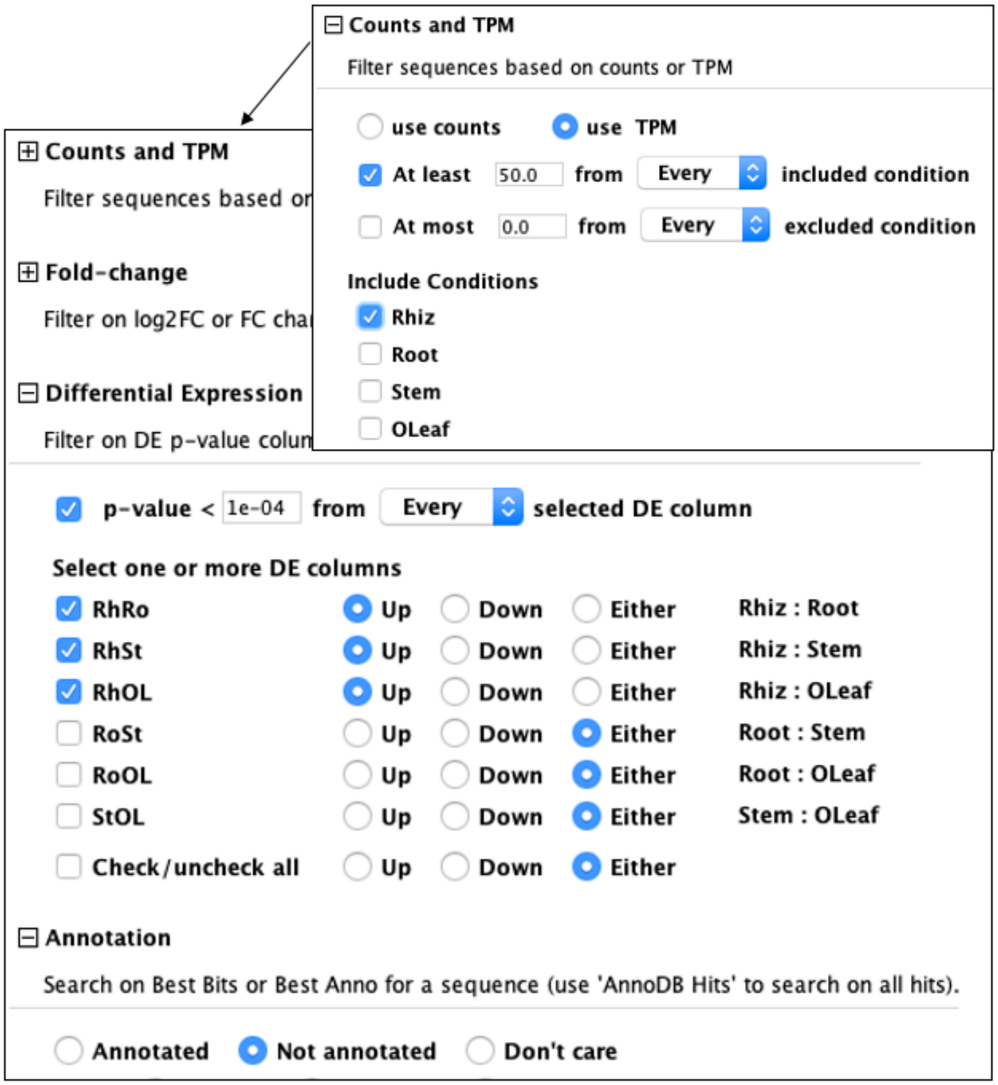


**Figure S5. Sequence Filters for TPM, DE and annotation.**

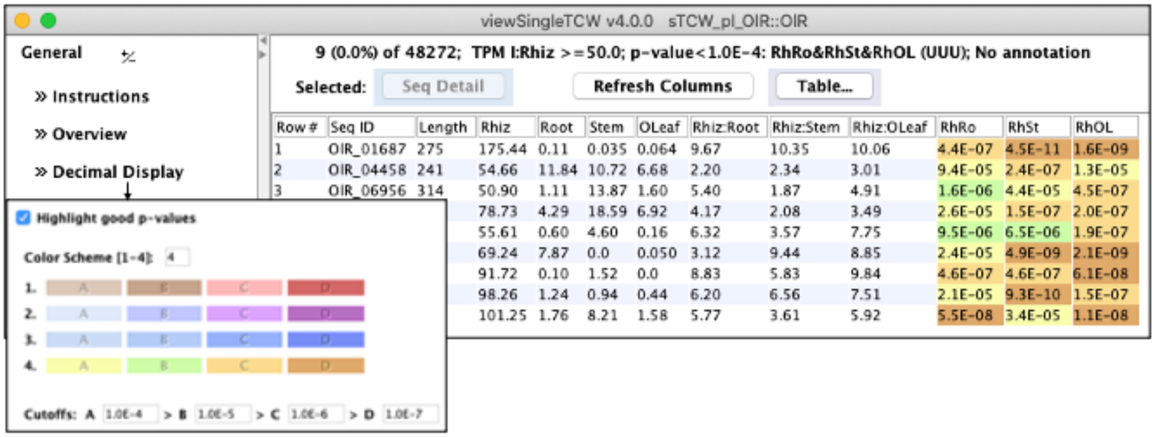


**Figure S6. Seq Table and Decimal Display.** The table is from the filters shown in Figure S5.

#### 2.1.6 Figure 5. GO enrichment of plant slims for OIR rhizomes

- Basic GO Annotation (Figure S7): Set the filters as shown and BUILD.
- Select Columns, select GOid, Ont, GO Name, RhRo, RhSt, RhOL. Sort on column Ont.
- Select Export..., option Table columns with -log10(Pval).
- Open the file with Excel to create the graph.

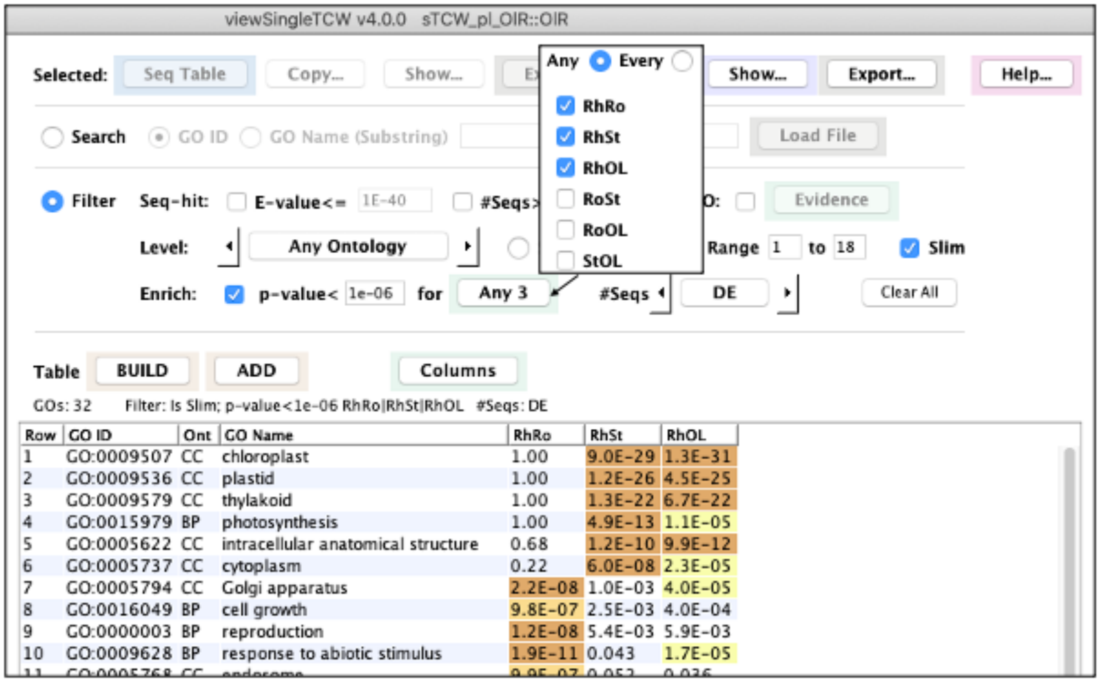


**Figure S7. Basic GO Annotation.** Plant Slims enriched for rhizome compared root (RhRo), stem (RhSt), and old leaf (RhOL).

For the 162 total GOs used for REVIGO and WEGO, start with the filters above, then change:
Uncheck Slim, select #Seq: DE, BUILD, Select All (bottom of table, not shown), Seq Table.
The #Seq: DE was set for input to REVIGO, as it causes the #Seq column (not shown) to provide the number of transcripts with DE<0.05, which is input to GOseq. If you view the Seq Table with the RhRo, RhSt and RhOL columns, you will notice that all sequences have at least one value > 0.05 . If the Basic GO #Seq: is set to All (instead of DE), then all sequences with at least one of the GOs will be shown, even if they do not have a value > 0.05.

#### 2.2 MultiTCW results

##### 2.2.1 Table 6: Summary of clusters

- Columns 2-5: Overview (Figure S8): All columns under Sizes, where some are combined, e.g. “=3” and “4-5” for the “3-5” column in manuscript.
- Column 6, 8: Overview (Figure S8): Use the #Seqs and Score2 column, respectively.
- Column 7: Cluster Filter (Figure S5 above):
  - Set the Cluster Set and set %Hit>=100 under Basic, then View Filtered Clusters.
  - Divide the number of rows by the cluster’s Total in the Overview.
- Columns 9-10: The average Ka and Ks columns were found as follows:
  - Pairs Filter (Figure S9): Select a Cluster Set, then View Filtered Pairs.
  - From the resulting Pairs Table (Figure S10), select the Show Table Stats option from the Table... pull-down. A pop-up of the statistics for the cluster will show the the average Ka and Ks.

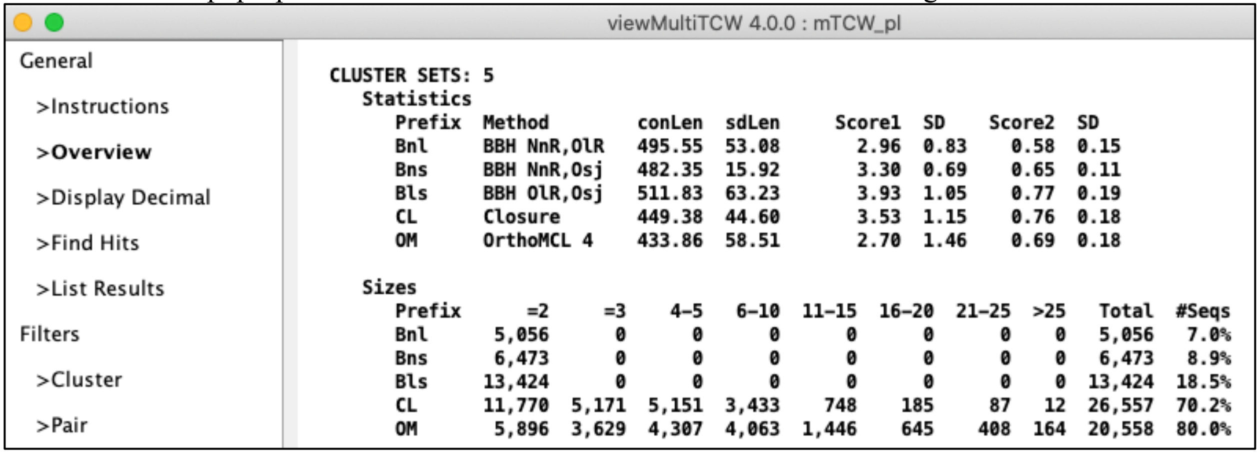


**Figure S8. MultiTCW overview of cluster section.**

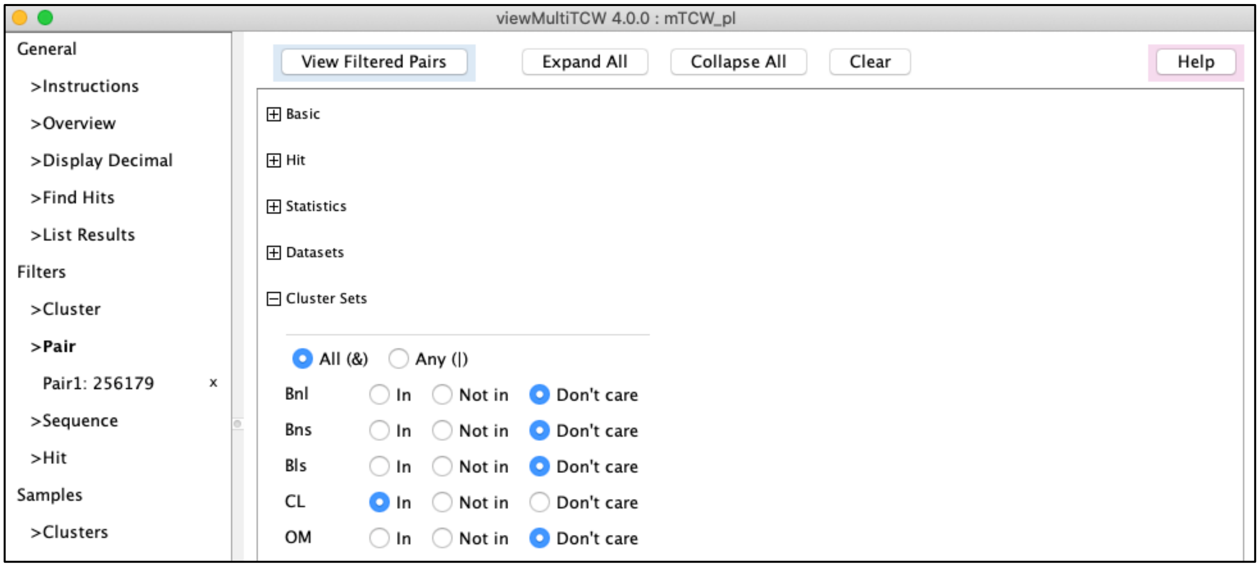


**Figure S9. Pairs Filter of the cluster set CL.**

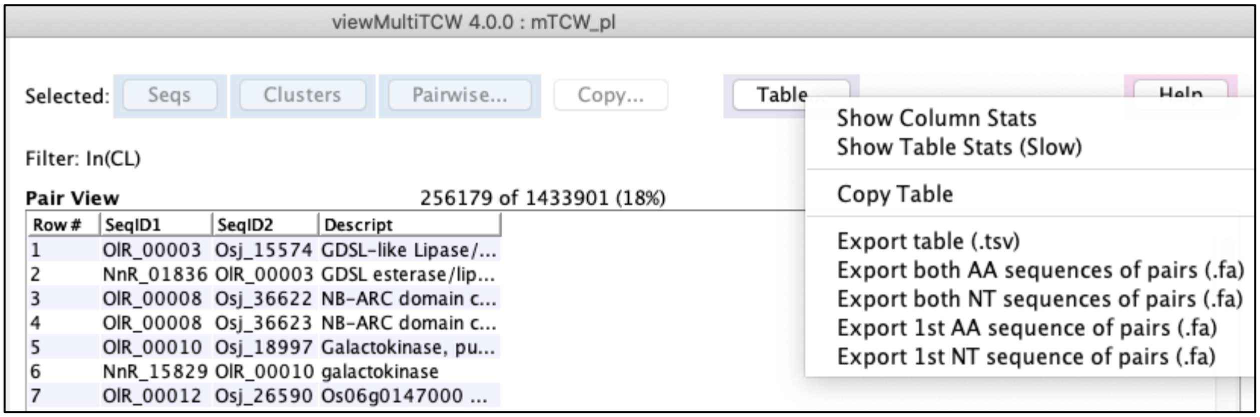

**Figure S10. Pairs Table of Closure pairs.**

#### 2.2.2 Figure 6. MSA and Pairwise

- Cluster Filter (Figure S11): Enter CL_009396.
- Figure 6A: From the Cluster Table (Figure S12), select MSAdb.
- Figure 6B: From the Cluster Table, select Pairs. From the resulting Pairs Table (Figure S13), select the first pair followed by the AA,CDS,NT option of the Pairwise dropdown.
- Figure 6C: Select the CDS option (Figure S14A) at the top of the pairwise alignment.

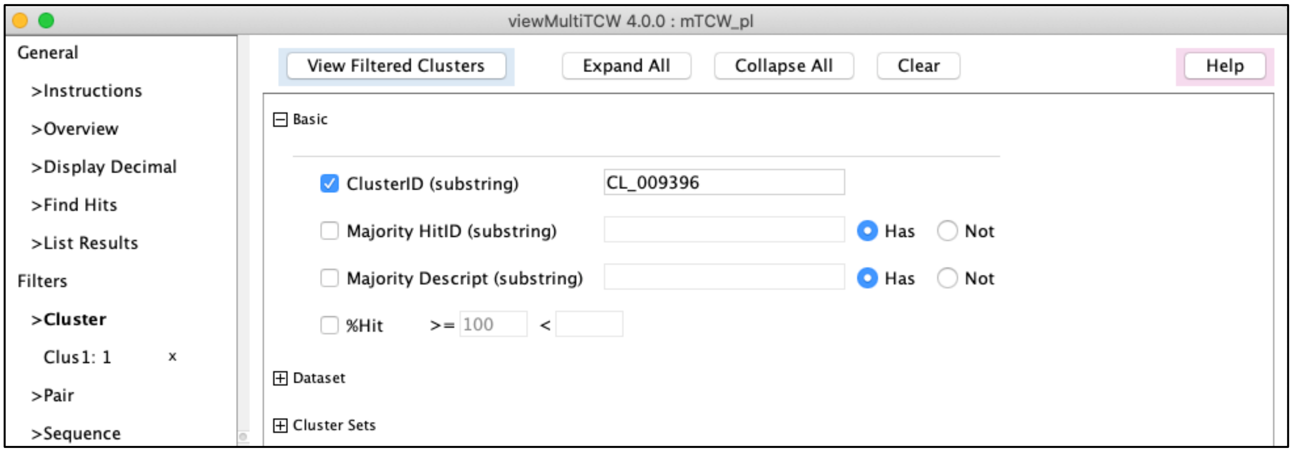

**Figure S11. Cluster Filter for specific cluster.**

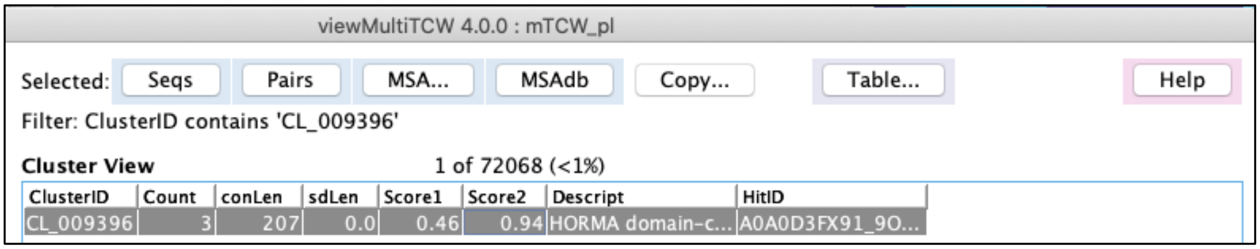

**Figure S12. Cluster Table showing the MSAdb option.**

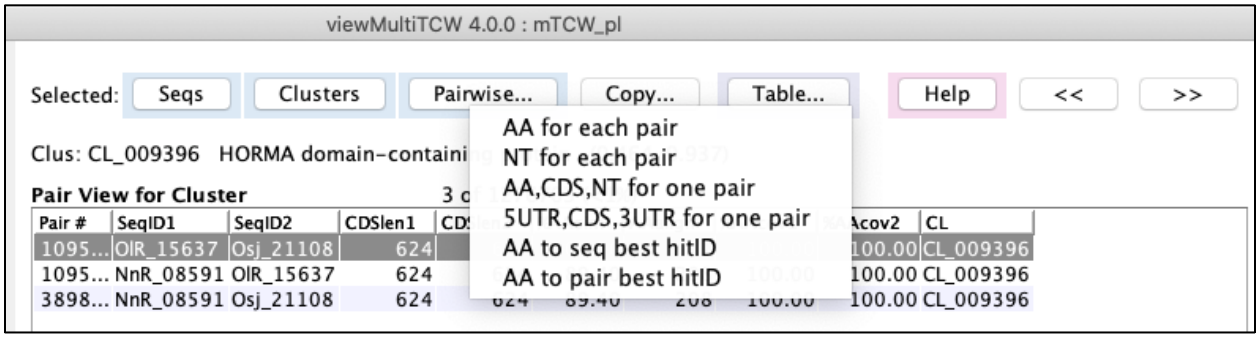


**Figure S13. Pairs Table for cluster.**

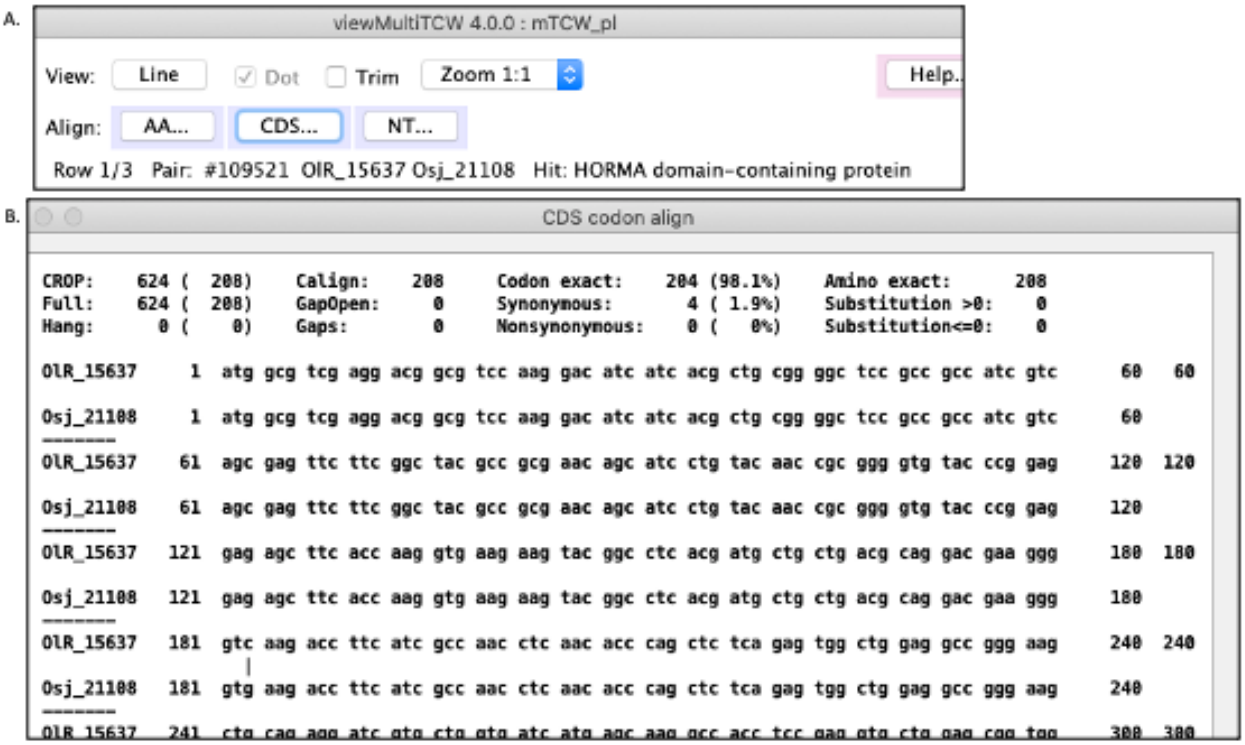


**Figure S14. Pairs alignment details.** (A) The pairwise alignment of AA,CDS,NT has options to show the text of the CDS. (B) The text alignment which shows the 4 synonymous codons (the full alignment is not shown, so only one synonymous codon is shown at "|" in row 181).

#### 2.2.3 Figure 7. Top ten annotated rhizome Closure clusters

- Cluster Filter (Figure S15): Set the filters as shown.
- Cluster Table (Figure S16):
  - Select Show Columns, select the columns shown. Sort the Wentroy (Score2) column by right clicking the column head.
  - For the average number of NnR and OlR sequences, use the Column Stats from the drop-down.

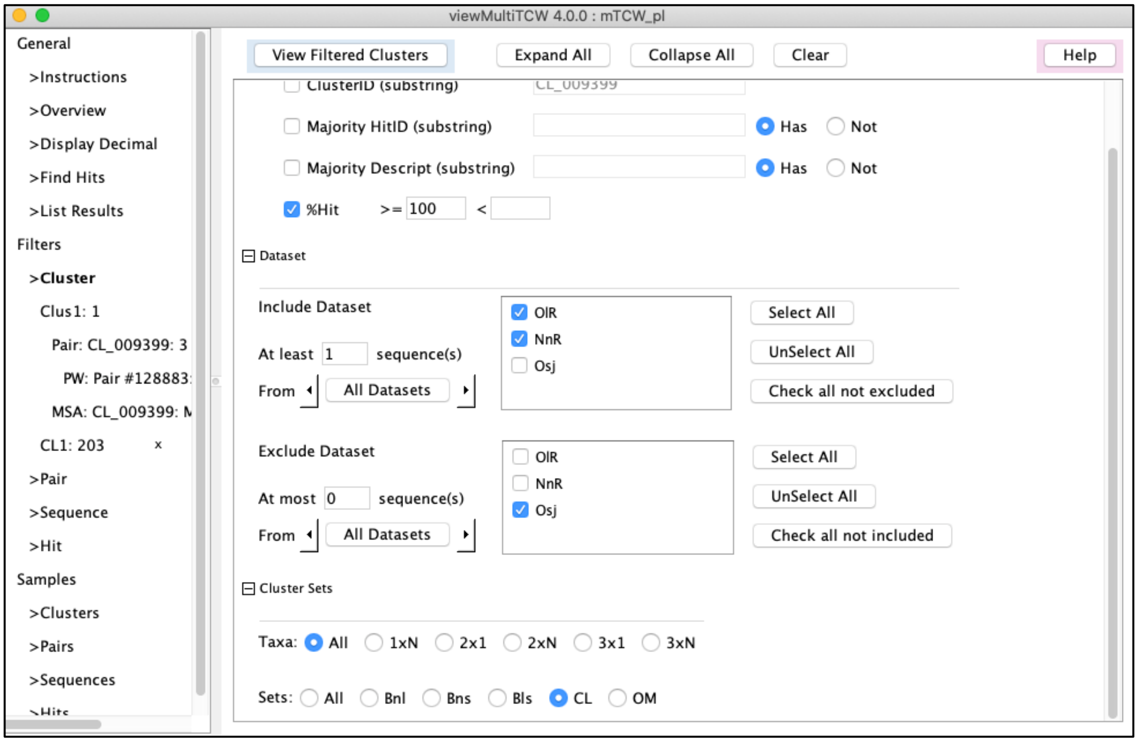


**Figure S15. Cluster Filter for rhizome only and shared annotation.**

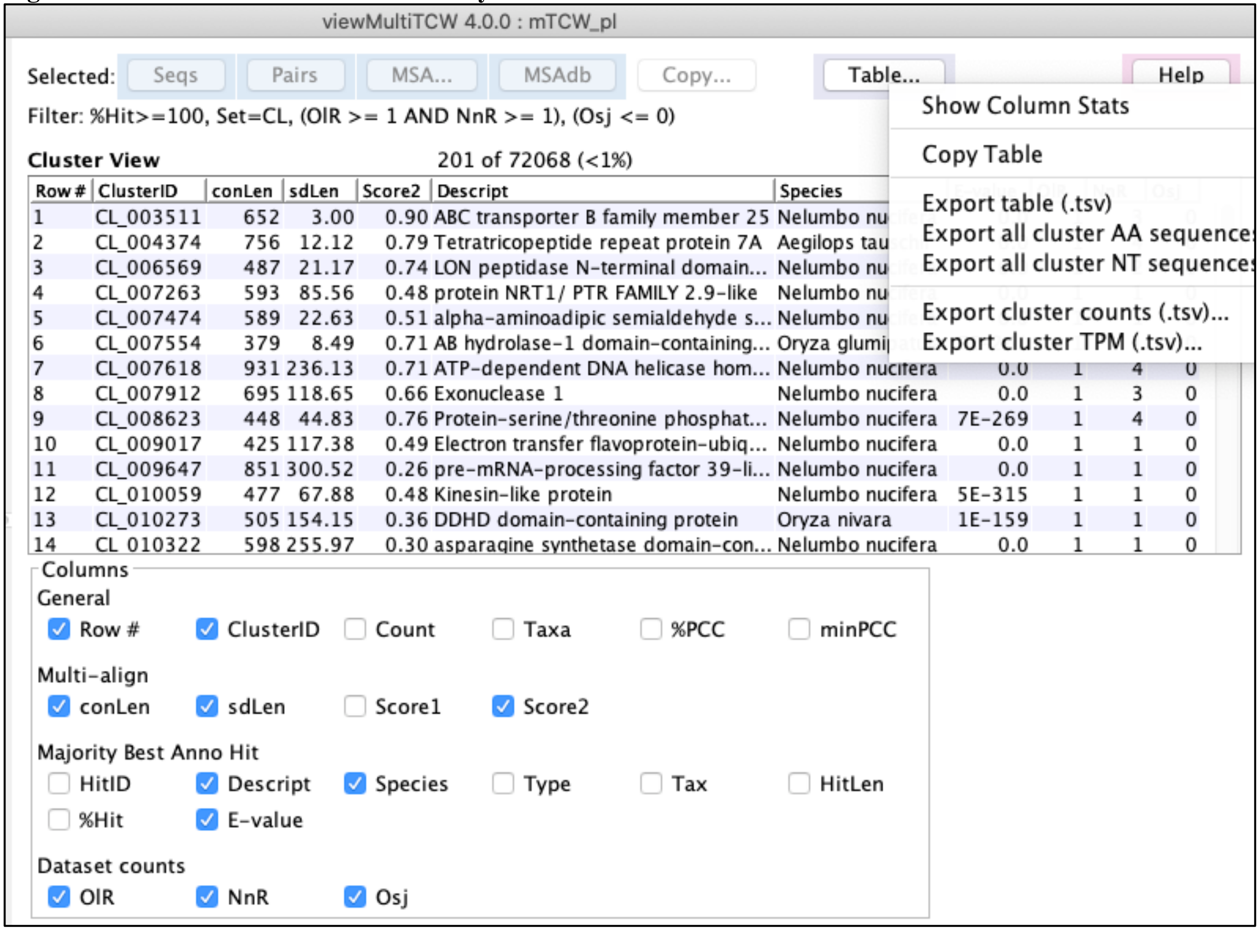


**Figure S16. Cluster Table for rhizome only with the column selection displayed.**

##### 2.2.4 Figure 8. Pair statistics for BBH OlR-Osj (Bls) and NnR-Osj (Bns)

- Pairs Filter (Figure S9 above): Select the Cluster Set.
- From the resulting Pairs Table (Figure S10 above), select the Show Table Stats option from Table....

Figure 8A is from the top part of the Bls stats and Figure 8B is the top part of the Bns stats.

##### 2.2.5 Figure 9. Ka/Ks for BBH OlR-Osj, top 10 conserved/diverged pairs

- Figure 9A: Pairs Filter (Figure S9 above): Select Cluster Set Bls (BBH OlR-Osj).
  From the resulting Pairs Table (Figure S10 above), select the Show Table Stats option.
- Figure 9B-C. Select the filters as shown in Figure S17 (make sure to have 0 in both boxes for Gap). Select KaKs < 1 for Figure 9B and KaKs ≥ 1.001 for Figure 9C (there is no way to request KaKs>1, hence, the 1.001).
- From the Pairs Table, select Show Columns to select the columns shown.

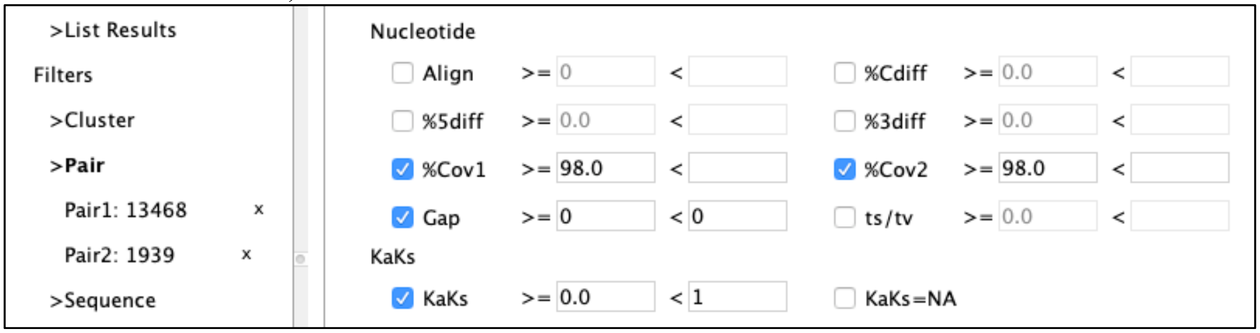


**Figure S17. Cluster Filter for BBH OlR-Osj conserved pairs .**

### 3 Timing results of builds

#### 3.1 Test machines

The builds were timed on a Linux and Mac machine, as specified in Table S1.

**Table S1: Linux and Mac**

| Machine | Purchase | Cores | RAM | SQL | Download |
|---|---|---|---|---|---|
| Linux x86-64 | 2011 Centos | 2.3Ghz 24-core AMD | 128 GB SSD | MariaDB 10.4.12 | 500 Mbsp |
| Mac 10.15.7 | 2019 Catalina | 3.2Ghz 6-core Intel i7 | 64 GB DDR4 | MySQL 1.8 | 150 Mbsp |

DIAMOND, BLAST and MAFFT use multi-processors, whereas all the TCW processing is single process. Therefore, it is advantageous to have a multi-processor for the searching large datasets.

#### 3.2 Build single database

Table S2 shows the timing of all steps for building sTCW_OlR, where the times for the no-pruned and description-pruned annotation are shown on Linux, and only the description-pruned annotation on Mac. S2 Suppl §2.1.2 shows that the un-pruned OlR has 1030k unique hits with GOs whereas the description pruned OlR has 134k unique hits with GOs, which reduced the time for adding GOs.

**Table S2: Linux and Mac timing results for building sTCW_OlR**

| | No Prune | Description Prune | |
|---|---|---|---|
| | Linux | Linux[a] | Mac |
| **runAS** | | | |
| Download[b] | | | |
| 5 uniprot_sprot_*taxo*[c].dat.gz (Max 214.4Mb) | 1m:10s | 1m:10s | 1m:00s |
| 1 uniprot_sprot.dat.gz (625.8Mb) | 1m:11s | 1m:11s | 0m:53s |
| 1 uniprot_trembl_plants.dat.gz (12.2Gb) | 15m:56s | 15m:56s | 15m:33s |
| Create UniProt 5 SP and 1 TR FASTA | 15m:39s | 15m:39s | 5m:39s |
| Create SwissProt Subset FASTA | 0m:58s | 0m:58s | 0m:16s |
| Build GO database | 42m:54s | 42m:54s | 21m:36s |
| *TOTAL* | *1h:17m:48s* | *1h:17m:48s* | *44s:57s* |
| **runSingleTCW** | | | |
| Build with 48,272 transcripts | 7m:05s | 7m:05s | 4m:29s |
| Annotate: | | | |
| Search and add annotations[d] | 1h:06m:53s | 1h:10m:08s | 50m:17s |
| ORF-finding | 17m:54s | 17m:45s | 6m:17s |
| Add GO annotations | 1h:03m:28s | 41m:35s | 22m:32s |
| *TOTAL* | *2h:28m:15s* | *2h:09m:28s* | *1h:19m:06s* |
| **runDE** | | | |
| edgeR for RhRo (rhizome-root)[e] | 1m:16s | 1m:16s | 0m:21s |
| GOseq for RhRo (rhizome-root)[e] | 3m:00s | 3m:00s | 0m:51s |

[a] All the times are the same as the "No Prune" Linux except for the 4 annotation times.
[b] There can be considerable variation in download times.
[c] *Taxo* = plants, fungi, virus, bacteria, invertebrate
[d] This did not include for formatting the DIAMOND database but did include the time for searching.
[e] The time for executing the R code and loading the results for one DE column.

#### 3.3 Build multi database

Table S3 has timing for the database mTCW_pl, which has input sTCW_OlR, sTCW_NnR and sTCW_Osj.

**Table S3: mTCW_pl – input three non-pruned sTCWdbs**

| Step | | Linux |
|---|---|---|
| Build mTCW_pl | 144,745 transcripts | 31m:04s |
| AA self-diamond and NT self-blast | | 13m:11s |
| Add all pairs from AA&NT files | 1,212,949 all pairs | 1h:01m:34s |
| Build Clusters (3 BBH, Closure, OrthoMCL[a]) | 5 sets | 36m:58s |
| Compute PCC for all pairs | | 7m:41s |
| Align and analyze cluster pairs[b] | 516,898 pairs | 9h:46m:47s |
| Run MAFFT and analyze MSA[c] | 72,133 clusters | 9h:23m:39s |
| Add KaKs values for cluster pairs | | 2m:54s |
| Add GO annotation[d] | 24,784 unique GOs | 1h:36m:02s |
| *TOTAL* | | *23h:18m:56s* |

[a] OrthoMCL took 15m:17s of the 36m:58s.
[b] Only pairs in clusters are aligned. A future release will parallelize this step.
[c] This pre-computes the MSA and scores to store for display, but can be computed on-the-fly in viewMultiTCW.
[d] The GO annotation is not necessary; however, viewMultiTCW does have some queries for it.

Table S4 is timing for the database mTCW_rhi, which has input sTCW_OlR and sTCW_NNU, where the sTCW_NNU has the input sequences as specified in §1 of this supplement. They both were pruned (hence, less descriptions) and have counts.

**Table S4: mTCW_rhi – input two "description pruned" sTCWdbs**

| **Step** | | **Linux** | **Mac** |
|---|---|---|---|
| Build mTCW_rhi | 74,957 transcripts | 9m:11s | 5m:31s |
| AA self-diamond and NT self-blast | | 2m:54s | 2m:00s |
| Add all pairs from AA&NT file | 389,983 all pairs | 16m:15s | 5m:27s |
| Build BBH and Closure clusters | 2 sets | 2m:27s | 1m:13s |
| Compute PCC for all pairs | 389,983 all pairs | 2m:04s | 1m:26s |
| Align and analyze cluster pairs | 31,929 cluster pairs | 15m:18s | 6m:16s |
| Run MAFFT and analyze MSA of | 16,620 clusters | 53m:53s | 15m:57s |
| Add KaKs values for cluster pairs | 31,929 cluster pairs | 0m:16s | 0m:15s |
| Add GO annotation | 24,218 unique GOs | 42m:33s | 24m:47s |
| *TOTAL* | | *2h:24m:51s* | *1h:02m:52s* |

#### 3.4 Memory

The maximum memory used for building sTCW_OlR (no prune) was for adding GOs at 1442Mb. The maximum for the description pruned sTCW_OlR was 631Gb on both Linux and MacOS.

The MySQL (MariaDB 10.4.7) database size for sTCW_OlR was 2.2Gb and mTCW_pl was 4.3Gb. The pruned sTCW_OlR was 1.3Gb and the smaller mTCW_rhi was 1.4Gb.

#### 3.5 Timing functions

The timings used the following Java functions:

```
long startTime  = System.nanoTime();
long elapseTime = System.nanoTime() – startTime;
```

Approximate memory usage were computed by:

```
Runtime runtime = Runtime.getRuntime();
runtime.gc();
double memory   = runtime.totalMemory() – runtime.freeMemory();
```

This is the total memory used, not just the memory used for the given method, so may vary by machine.

The Linux database build and annotate functions were run multiple times and gave similar timing results. The download times on both Linux and Mac can vary considerably.

### 4 External software and databases

With the exception of the R packages (edgeR, DEseq, GOseq), all external software comes with the TCW package (rows 1-8 of Table S5). The version numbers apply to the packages provided with TCW V4.

Table S5. External software and databases used by TCW.

| | | |
|---|---|---|
| 1 | DIAMOND v2.0.11.149 [5,6] | Used for building the sTCWdb and mTCWdb. Also used for searching in both viewSingleTCW and viewMultiTCW. |
| 2 | BLAST 2.12.0+ [7] | Same as above |
| 3 | CAP3 [8] | Used for assembling ESTs or transcripts for runSingleTCW. |
| 4 | OrthoMCL v2.0.9 [9] | Used for building clusters in runMultiTCW. |
| 5 | MAFFTA v7.407 [10] | Used for computing MSA in runMultiTCW and viewMultiTCW. |
| 6 | MUSCLE v3.8.31 [11] | Same as above. |
| 7 | MstatX [12] | Used for scoring MSAs. |
| 8 | KaKs_calculator 1.2 [13] | Computing Ka/Ks for pairs for runMultiTCW. |
| 9 | edgeR [14], DEseq [15], GOseq [16] | R packages that must be installed by the user. TCW provides scripts to run them from runDE. |
| 10 | UniProt [17] | runAS provides an interface to download and format SwissProt and TrEMBL taxonomic database for input to runSingleTCW. |
| 11 | Gene Ontology [18] | runAS provides an interface to download and build the GO database for input to runSingleTCW. |
