## Supplementary material for "Transcriptome computational workbench (TCW): analysis of single and comparative transcriptomes": S2 Suppl

### Build a singleTCW database

**Sections**

The purpose of this supplement is to provide additional information about the algorithms and parameters. ViewSingleTCW is used to demonstrate the results, where Figure S1 shows the interface features.

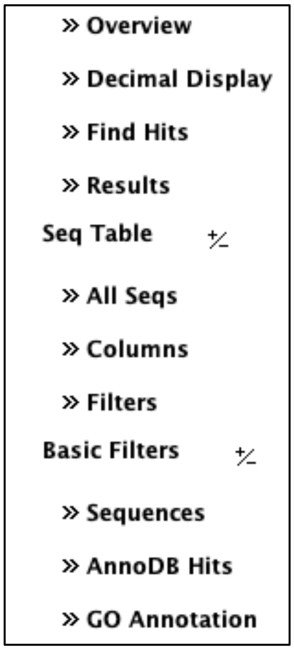

**Figure S1. The panels shown on the left of viewSingleTCW**

Decimal Display changes the formatting of decimal numbers and the highlighting of p-values.

Find Hits allows BLAST or DIAMOND searching against the sequences in the database or an external database.

Seq Table
Columns sets the columns for the Seq Table.
Filters provides filters on the sequences, resulting in a Seq Table.

Basic Filters
Sequences provides some specific filters for sequences.
AnnoDB Hits provides filter and display for the hits.
GO Annotations provides filter and display for the GOs.

From the Seq Table, a row can be selected to view the Seq Detail.
From the Seq Detail, the Frame, GOs and Hit Alignment can be viewed.

### 1 runAS: annotation setup

The runAS program provides a graphical interface (Figure S2) that creates the necessary UniProt files and TCW GO database. First, all desired taxonomic UniProt SwissProt and TrEMBL .dat files are downloaded and a corresponding FASTA file of the sequences is created for each. After all taxonomic files have been downloaded, the full SwissProt can be downloaded and a subset FASTA file is created by removing all records found in the downloaded taxonomic files. This can also be done for the full TrEMBL, but it has gotten so big that it requires a tremendous amount of memory to process and typically is not worth it; that is, by selecting all relevant SwissProt and TrEMBL taxonomic databases and the full SwissProt, the best and most relevant protein hits will be obtained.

The runAS program creates the TCW-GOdb by first downloading the latest go-basic.obo file from archive.geneontology.org, where the file contains the GO terms, descriptions, relations and GO Slim subsets. Then runAS creates the TCW-GOdb MySQL database and populates it with the data from go-basic.obo. Three additional tables are created in the database: the first contains the name of the GO file downloaded, the second contains GO levels, and the third contains all the relevant UniProt information from the downloaded .dat files, which is the direct GO terms, InterPro, KEGG, EC and Pfam assignments for each protein. The GO levels and ancestors are computed and entered into the GOdb.

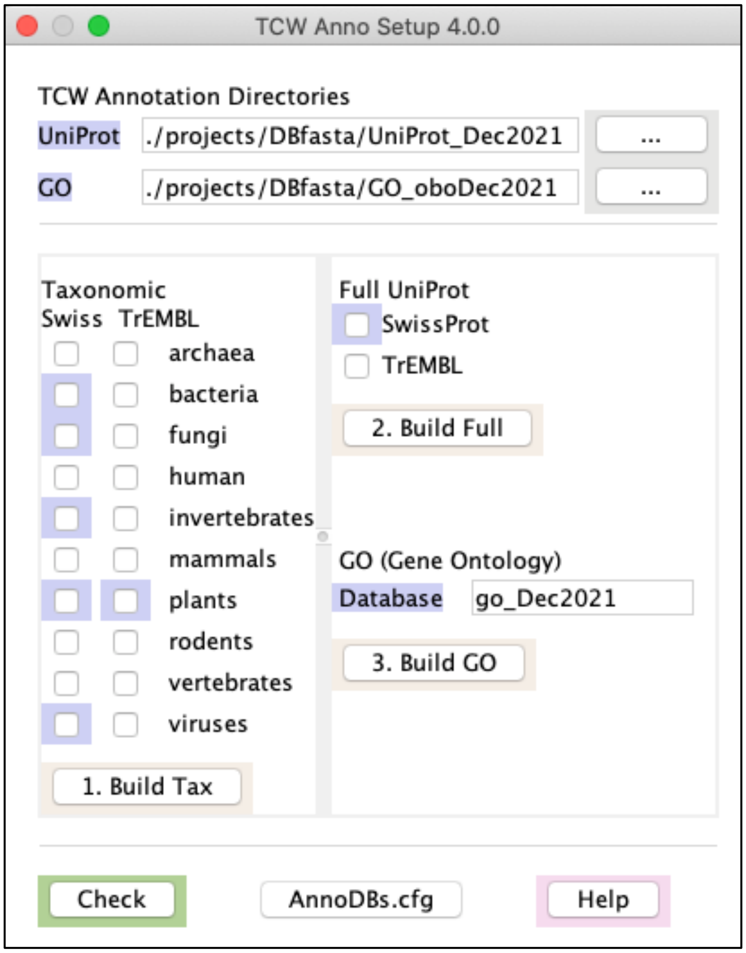

**Figure S2. runAS: annotation setup.** The purple highlights indicate completed downloads and processing.

#### 2 runSingleTCW: annotation

The runSingleTCW interface is shown in Figure S3. Step1 loads the data. Step2 assembles or instantiates (finalizes) the input sequences. Step 3 annotates the sequences with hits, assigns GOs, and compute ORFs.

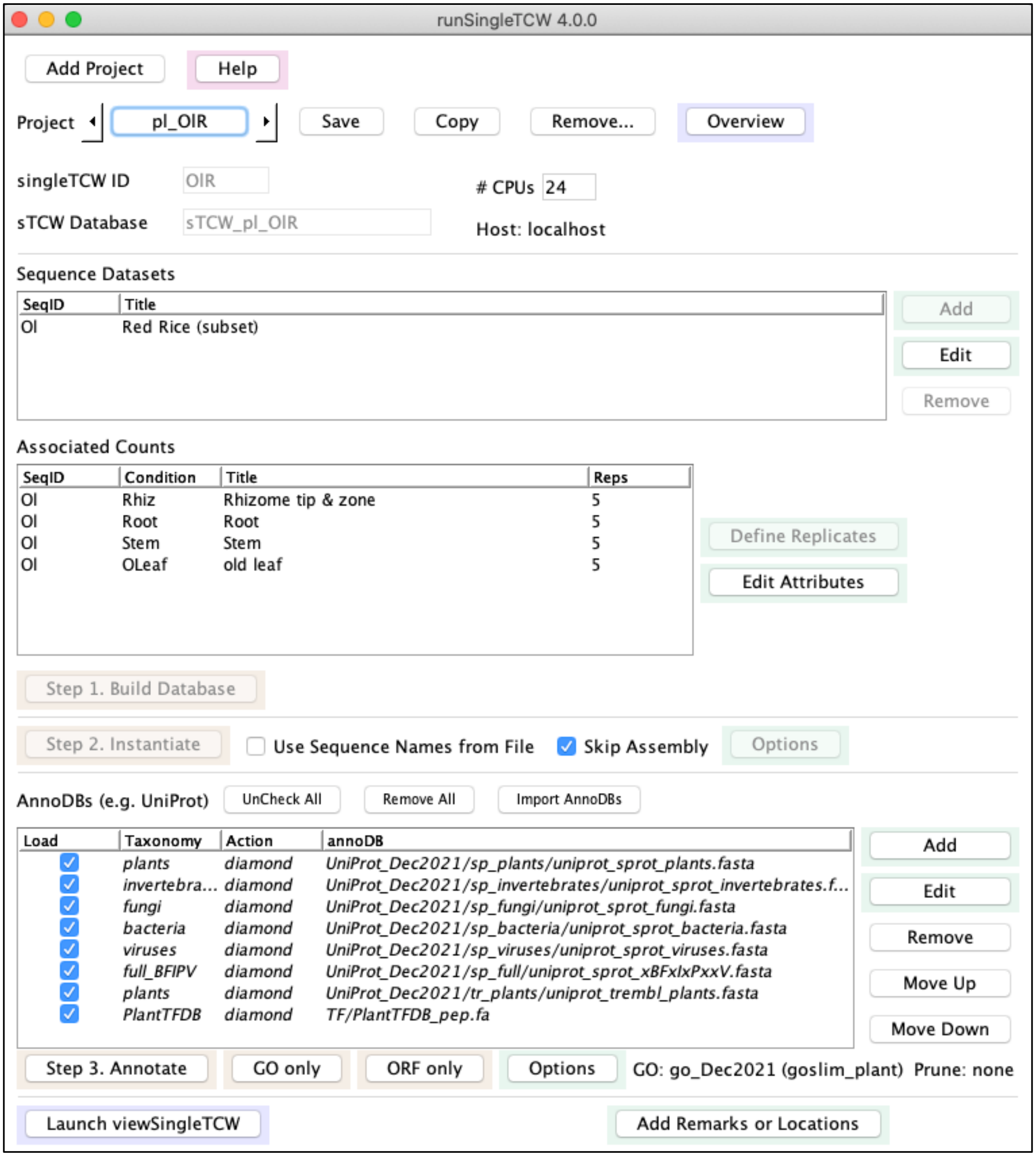

**Figure S3. runSingleTCW: create and annotate a singleTCW database.** The Step 1 and Step 2 buttons are inactive because they have been run. The annoDBs are italicized because they have been loaded into the database; additional annoDBs can be added.

#### 2.1 Annotate

The user defines one or more “annoDBs” (annotation databases), which can be protein or nucleotide FASTA files. All sequences in the sTCWdb are searched against each annoDB file. The results are parsed and the first entry for each sequence-hit pair is loaded into the sTCWdb. The FASTA files are parsed for the description and species of each hit and loaded into the sTCWdb. After all sequence hits are loaded, the Best Bits, Best Anno and other filters are computed. The Best GO is determined after the GOs are loaded.

The Best Bits, Best Anno, Best GO and other filters are computed as follows:

- *Best Bits:* The hits are sorted by {bit-score, is_SwissProt, is_TrEMBL, E-value} where the top hit is assigned. This gives precedence to UniProt hits, as they may have GOs and other annotations.
- *Best Anno:* Using the sorted list, the first one that is considered informative is assigned the Best Anno, where the uninformative rules are described in the next section.
- *Best GO*: The best hit that has GO information. The list is sorted by {bit-score, is_BestBits, is_BestAnno, E-value} and the top hit is assigned. This gives precedence to the Best Bit or Best Anno assigned earlier.
- *Other filters*: Other assignments are made in order to filter hits when viewing a sequence, where the filters are: best hits, best per annoDB, representative set of hits with distinct regions, unique species, and unique descriptions.

##### 2.1.1 Uninformative annotation

The “International Protein Nomenclature Guidelines”[1] suggests that when there is no known domain or motif, the description should be “hypothetical protein”, “uncharacterized protein, “protein GS” or “GS protein” (GS = gene symbol or gene protein). However, there are other description such as “expressed protein” that TCW treats as uninformative. The algorithm first removes any text within brackets, changes the description to lower-case, then marks the description as uninformative if any of the following are true about the description:

1. Contains "uncharacterized protein", "hypothetical protein", "putative uncharacterized", "predicted protein", "whole genome shotgun", "scaffold", "unnamed protein product".
2. Starts with “ORF”, "low quality protein" or "cDNA clone".
3. Equals "expressed protein".
4. Has only 1 or 2 words where the first word contains a mix of letters and digits, and the last word (if it exists) is “protein”, “partial”, “(fragment)” or “scaffold” (e.g. Figure S4 row 7).

These rules are ad hoc and were written from viewing plant and invertebrate results. The rules can easily be changed in the source file “util/methods/BestAnno.java”.

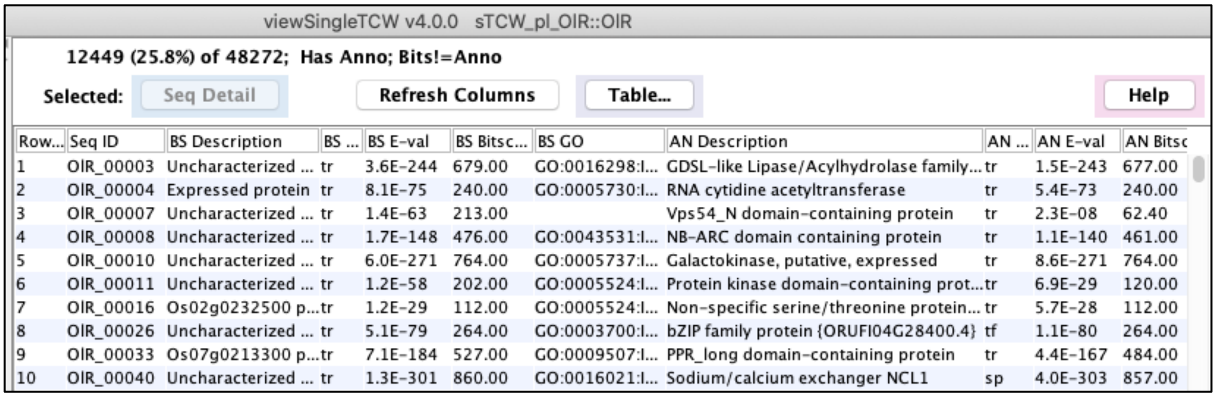

| Row... | Seq ID | BS Description | BS ... | BS E-val | BS Bitsc... | BS GO | AN Description | AN ... | AN E-val | AN Bitsc |
|---|---|---|---|---|---|---|---|---|---|---|
| 1 | OIR_00003 | Uncharacterized ... | tr | 3.6E-244 | 679.00 | GO:0016298:I... | GDSL-like Lipase/Acylhydrolase family... | tr | 1.5E-243 | 677.00 |
| 2 | OIR_00004 | Expressed protein | tr | 8.1E-75 | 240.00 | GO:0005730:I... | RNA cytidine acetyltransferase | tr | 5.4E-73 | 240.00 |
| 3 | OIR_00007 | Uncharacterized ... | tr | 1.4E-63 | 213.00 | | Vps54_N domain-containing protein | tr | 2.3E-08 | 62.40 |
| 4 | OIR_00008 | Uncharacterized ... | tr | 1.7E-148 | 476.00 | GO:0043531:I... | NB-ARC domain containing protein | tr | 1.1E-140 | 461.00 |
| 5 | OIR_00010 | Uncharacterized ... | tr | 6.0E-271 | 764.00 | GO:0005737:I... | Galactokinase, putative, expressed | tr | 8.6E-271 | 764.00 |
| 6 | OIR_00011 | Uncharacterized ... | tr | 1.2E-58 | 202.00 | GO:0005524:I... | Protein kinase domain-containing prot... | tr | 6.9E-29 | 120.00 |
| 7 | OIR_00016 | Os02g0232500 p... | tr | 1.2E-29 | 112.00 | GO:0005524:I... | Non-specific serine/threonine protein... | tr | 5.7E-28 | 112.00 |
| 8 | OIR_00026 | Uncharacterized ... | tr | 5.1E-79 | 264.00 | GO:0003700:I... | bZIP family protein {ORUFI04G28400.4} | tf | 1.1E-80 | 264.00 |
| 9 | OIR_00033 | Os07g0213300 p... | tr | 7.1E-184 | 527.00 | GO:0009507:I... | PPR_long domain-containing protein | tr | 4.4E-167 | 484.00 |
| 10 | OIR_00040 | Uncharacterized ... | tr | 1.3E-301 | 860.00 | GO:0016021:I... | Sodium/calcium exchanger NCL1 | sp | 4.0E-303 | 857.00 |

**Figure S4. Seq Table of BS Bits != Best Anno.** Results of the Sequence Filter showing sequences where the BS hit is not the same as the AN (BS=Best Bits, AN=Best Anno). The BS annotations are considered uninformative, though many have GO annotations (column BS GO).

#### 2.1.2 Prune Hits

Same alignment. Figure S5 shows the results for a sequence from an un-pruned sTCWdb. The highlighted set have identical alignments. RunSingleTCW has an option to prune hits based on identical alignments, which would remove all the hits but the first that have identical alignments.

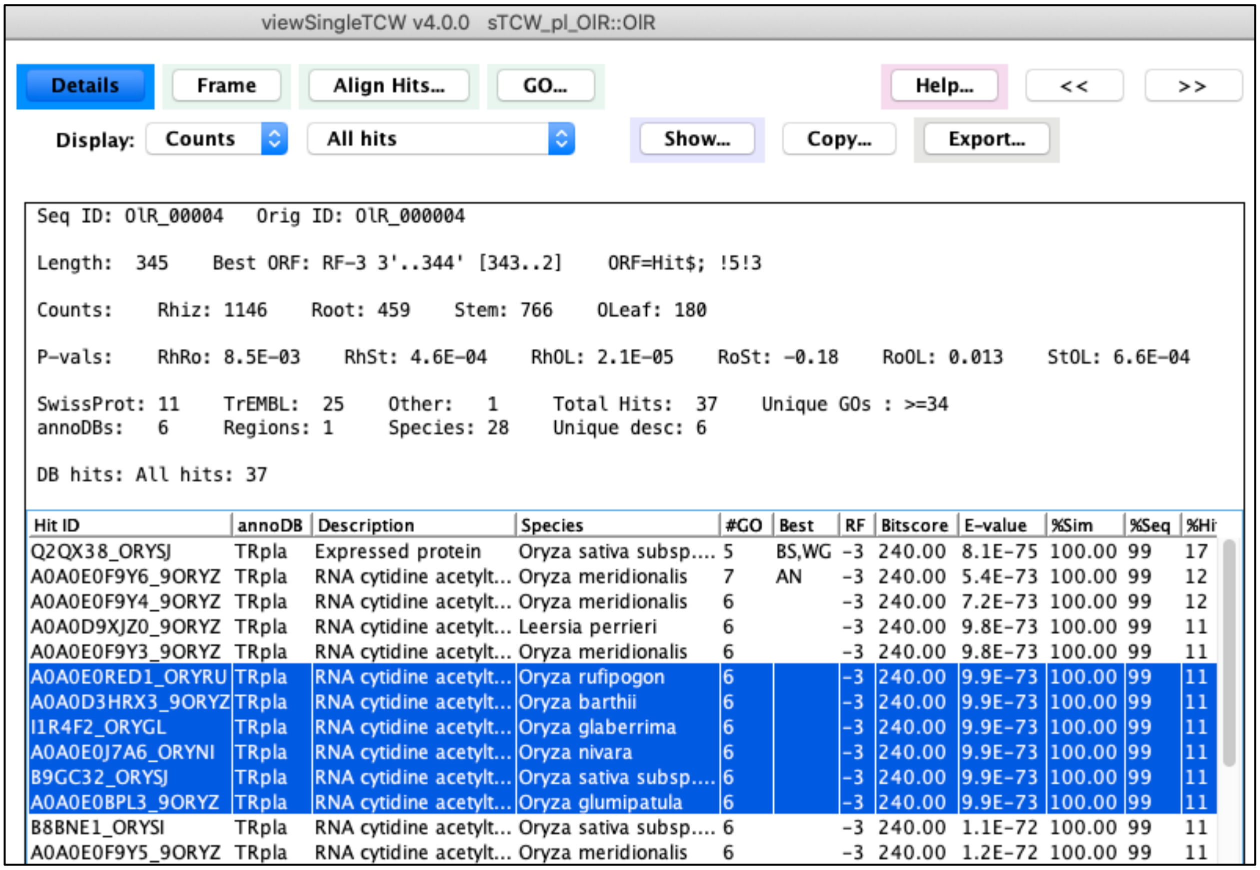

**Figure S5. Seq Detail with all hits.** This view shows all information available for a sequence The hits table is set to show All hits, where BS=Best Bits, AN=Best Anno, WG=With GOs. The highlighted hits have the same alignment.

Similar description. As illustrated in Figure S5, there are many redundant descriptions. RunSingleTCW has an option to prune all descriptions that are the similar for each annoDB; the comparison is not case-sensitive and does not include any text at the end that is within brackets. The description with the best bit-score and E-value is retained; when there is a tie, then the one with the most GOs is retained. Figure S6 shows the same transcript as Figure S5 but with pruned descriptions; the un-pruned has 37 hits and the pruned has 12. There are the same number of "Unique desc" indicating that the only information lost is the list of species that share the description. The pruned set still has some identical descriptions because they are pruned per annoDB, e.g. SPpla (plants), Spinv (invertebrates), SPfun (fungi) and SPful (full) all have "RNA cytidine actyltransferase".

SwissProt: 7    TrEMBL: 4    Other: 1    Total Hits: 12    Unique GOs : >=33
annoDBs: 6    Regions: 1    Species: 11    Unique desc: 6

DB hits: All hits: 12

| Hit ID | annoDB | Description | Species | #GO | Best | RF | Bitscore | E-value |
|---|---|---|---|---|---|---|---|---|
| Q2QX38_ORYSJ | TRpla | Expressed protein | Oryza sativa subsp.... | 5 | BS,WG | -3 | 240.0 | 8.1E-75 |
| A0A0E0F9Y6_9ORYZ | TRpla | RNA cytidine acetyltransferase | Oryza meridionalis | 7 | AN | -3 | 240.0 | 5.4E-73 |
| A0A811RMN9_9POAL | TRpla | Hypothetical protein | Miscanthus lutariori... | --- | | -3 | 239.0 | 1.6E-72 |
| A0A5J9VLR2_9POAL | TRpla | Uncharacterized protein | Eragrostis curvula | 6 | | -3 | 239.0 | 2.9E-72 |
| TCP_1661 | TFpla | TCP family protein {ltr_sc004426.1_g00... | Ipomoea trifida | --- | | -3 | 222.0 | 1.5E-67 |
| NT101_ARATH | SPpla | RNA cytidine acetyltransferase 1 | Arabidopsis thaliana | 6 | | -3 | 215.0 | 3.3E-66 |
| NT102_ARATH | SPpla | RNA cytidine acetyltransferase 2 | Arabidopsis thaliana | 6 | | -3 | 206.0 | 4.9E-63 |
| NAT10_ACHAM | SPpla | RNA cytidine acetyltransferase | Achlya ambisexualis | 5 | | -3 | 170.0 | 5.4E-53 |
| NAT10_DICDI | SPinv | RNA cytidine acetyltransferase | Dictyostelium discoi... | 6 | | -3 | 161.0 | 2.0E-47 |
| NAT10_YEAST | SPfun | RNA cytidine acetyltransferase | Saccharomyces cer... | 11 | | -3 | 157.0 | 1.3E-45 |
| NAT10_HUMAN | SPful | RNA cytidine acetyltransferase | Homo sapiens | 21 | | -3 | 174.0 | 3.8E-51 |
| TMCA_THEPD | SPful | tRNA(Met) cytidine acetyltransferase TmcA | Thermofilum pendens | 6 | | -3 | 43.5 | 2.1E-05 |

**Figure S6. Seq Detail with pruned hits.** The same as Figure S5 except with pruned descriptions per annoDB.

Figure S7A shows the Overview of AnnoDBs with no pruning and Figure S7B shows it with description pruning. All columns to the right of Rank=1 have the same numbers in both Overviews. The ones to the left include all hits, where the Best Bits (BITS) and Best Anno (ANNO) are almost the same, but the UNIQUE and TOTAL are much smaller for the pruned. The total sequence hits is reduced by 64%; in contrast, the alignment pruning reduction is 11% .

A. No prune

Hit Statistics:
Sequences with hits 36,510 (75.6%)    Bases covered by hit 26,087,478 (50.1%)
Unique hits 719,247    Total bases 52,081,922
Total sequence hits 1,537,910

annoDBs (Annotation databases): 8 (see Legend below)

| ANNODB | ONLY | BITS | ANNO | UNIQUE | TOTAL | AVG %SIM | Rank =1 | HAS HIT | (%Seqs) | AVG %SIM | COVER >=50 | COVER >=90 |
|---|---|---|---|---|---|---|---|---|---|---|---|---|
| SP-plants | 52 | 2,300 | 3,843 | 27,934 | 229.4k | 50.9 | | 21,249 | (44.0%) | 67.2 | 35.2% | 6.1% |
| SP-invertebrates | 17 | 186 | 214 | 10,168 | 79,446 | 42.1 | | 9,280 | (19.2%) | 44.7 | 16.0% | 0.2% |
| SP-fungi | 10 | 53 | 72 | 15,139 | 85,929 | 41.1 | | 8,912 | (18.5%) | 43.2 | 14.5% | <0.1% |
| SP-bacteria | 7 | 23 | 46 | 32,873 | 74,617 | 40.8 | | 5,394 | (11.2%) | 41.0 | 11.8% | 0% |
| SP-viruses | 0 | 0 | 3 | 696 | 8,936 | 32.3 | | 1,583 | ( 3.3%) | 35.3 | 2.3% | 0% |
| SP-full_BFIPV | 5 | 34 | 116 | 31,817 | 140.9k | 42.3 | | 10,623 | (22.0%) | 44.2 | 14.7% | 0.2% |
| TR-plants | 11,764 | 33,632 | 31,037 | 563.8k | 800.2k | 84.2 | | 36,349 | (75.3%) | 90.7 | 47.3% | 16.4% |
| TF-plantTFDB | 53 | 282 | 1,179 | 36,777 | 118.4k | 55.4 | | 13,463 | (27.9%) | 58.2 | 11.4% | 2.1% |

B. Description prune

Hit statistics:
Sequences with hits 36,510 (75.6%)    Bases covered by hit 26,045,367 (50.0%)
Unique hits 181,156    Total bases 52,081,922
Total sequence hits 546,669

Annotation databases (annoDBs): 8 (see Legend below)

| ANNODB | ONLY | BITS | ANNO | UNIQUE | TOTAL | AVG %SIM | Rank =1 | HAS HIT | (%Seqs) | AVG %SIM | COVER >=50 | COVER >=90 |
|---|---|---|---|---|---|---|---|---|---|---|---|---|
| SP-plants | 52 | 2,349 | 3,853 | 23,048 | 191.6k | 49.7 | | 21,249 | (44.0%) | 67.6 | 35.2% | 6.1% |
| SP-invertebrates | 17 | 189 | 218 | 6,993 | 54,051 | 39.4 | | 9,280 | (19.2%) | 45.1 | 16.0% | 0.2% |
| SP-fungi | 10 | 53 | 72 | 8,454 | 44,308 | 38.8 | | 8,912 | (18.5%) | 43.6 | 14.5% | <0.1% |
| SP-bacteria | 7 | 23 | 46 | 6,511 | 18,170 | 37.1 | | 5,394 | (11.2%) | 41.4 | 11.8% | 0% |
| SP-viruses | 0 | 0 | 3 | 414 | 5,163 | 32.8 | | 1,583 | ( 3.3%) | 35.8 | 2.3% | 0% |
| SP-full_BFIPV | 5 | 37 | 118 | 14,607 | 45,751 | 40.2 | | 10,623 | (22.0%) | 44.7 | 14.7% | 0.2% |
| TR-plants | 11,764 | 33,542 | 31,010 | 115.2k | 147.8k | 85.3 | | 36,349 | (75.3%) | 91.0 | 46.8% | 16.4% |
| TF-plantTFDB | 53 | 317 | 1,190 | 5,949 | 39,807 | 45.7 | | 13,463 | (27.9%) | 58.6 | 11.4% | 2.1% |

**Figure S7. Overview of annoDBs**. (A) With no pruning (the same as Figure 3 in the manuscript). (B) With description prune.

Figure S8A shows the Overview for the GO statistics with no pruning and Figure S8B shows it with description pruning. There are 4% less GOs for the pruned set. However, in the pruned set there are 6% more sequences where the Seq best hit has GOs; this is due to the algorithm selecting the hit with GOs when there is a tie.

A. No prune

```
Gene Ontology Statistics:
   Unique GOs           25,509              Unique hits with GOs  1,030,863  (78.9%)
   Sequences with GOs   32,455  (67.2%)     Seq best hit has GOs     25,390  (52.6%)
   Has goslim_plant         97

   biological_process   17,074  (66.9%)     is_a                     41,366
   molecular_function    6,055  (23.7%)     part_of                   4,212
   cellular_component    2,380  ( 9.3%)
```

B. Description prune

```
Gene ontology statistics:
   Unique GOs           24,585              Unique hits with GOs  134,929  (74.7%)
   Sequences with GOs   32,059  (66.4%)     Seq best hit has GOs   27,022  (56.0%)
   Has goslim_plant         97

   biological_process   16,495  (67.1%)     is_a                   39,753
   molecular_function    5,809  (23.6%)     part_of                 4,059
   cellular_component    2,281  ( 9.3%)
```

**Figure S8. Overview**. (A) GOs with no pruning. (B) GOs with description prune.

Typically, it would be advisable to use the description prune option as it saves memory resulting in faster queries, and still shows all relevant information. The only reason that is it not the default is that it removes information, which is not done without the user's consent.

##### 2.1.3 Multi-frame and stop codons in hits

The BLAST or DIAMOND results for a sequence may have hits to multiple frames or have stop codons within the hit region of a sequence. The multi-frame NnR transcripts were explored as follows: using the Basic Sequence panel, the TCW option was selected which searches the TCW Remark column, and 'Multi' was entered for the substring (Figure S9). This resulted in the 3497 transcripts with multiple hits.

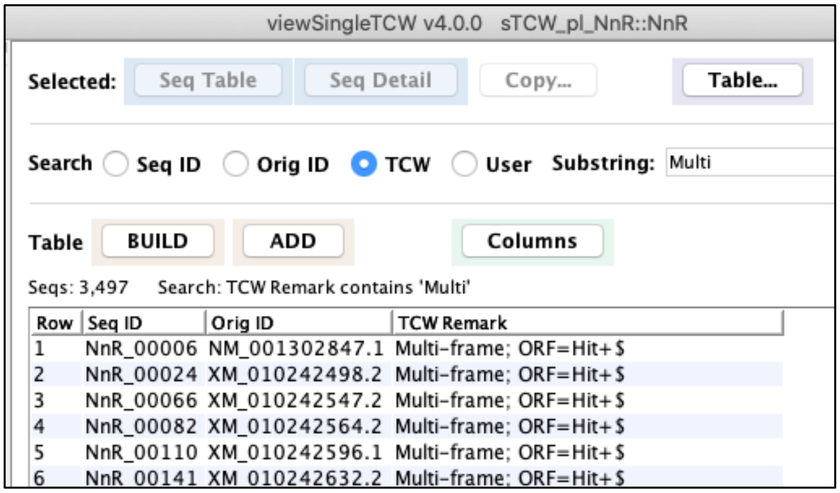

**Figure S9. Multiple frame hits.** The Basic Sequence table, which shows transcripts that have hits to multiple frames.

Using TCW to browse through the multi-frame transcripts, it shows that they tend to have one or more good hits to one frame and one or more poor hits to any other frames; for example, Figure S10A shows a sequence with a good hit to frame 1 and a poor hit to frame 3 (Display: Distinct Regions), and Figure S10B shows the alignment to the two frames (Align Hits…, Selected Hits).

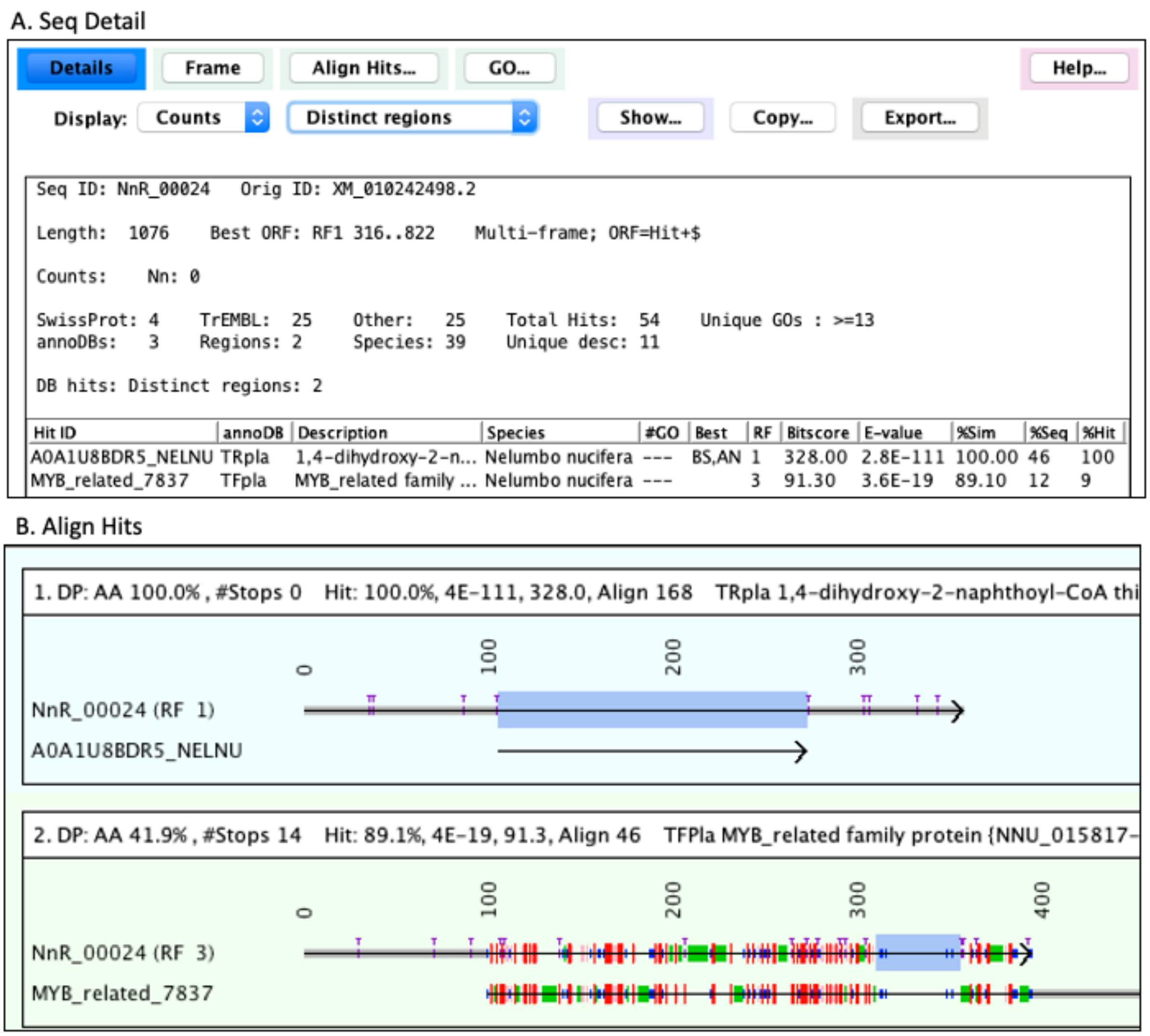

**Figure S10. Multiple frame hits for a sequence.** (A) Seq Detail for a transcript with hits to frame 1 and 3. (B) The alignment of two hits in different frames, where the blue highlighted region is the hit region.

To explore hits with stop codons in the sequence hit region, the "multi" was replaced with "stop" in the Basic Sequence panel, which resulted in 2315 transcripts. Figure S11 shows an example of a near perfect hit with one stop codon in the hit region. The alignment in Figure S11A was computed with dynamic programming (DP), which could give a different alignment than DIAMOND; therefore, to compare the DIAMOND alignment with the DP, the sequence was copied (using the Seq Detail panel Copy.. option), and searched against the TrEMBL plants database by using the Find Hits option; the section of the alignment with the stop codon is shown in Figure S11B. To further confirm the stop, the transcript sequence was entered into the online UniProt site (www.uniprot.org) to be blasted against the UniProt plant database, which also confirmed the stop codon (Figure S11C).

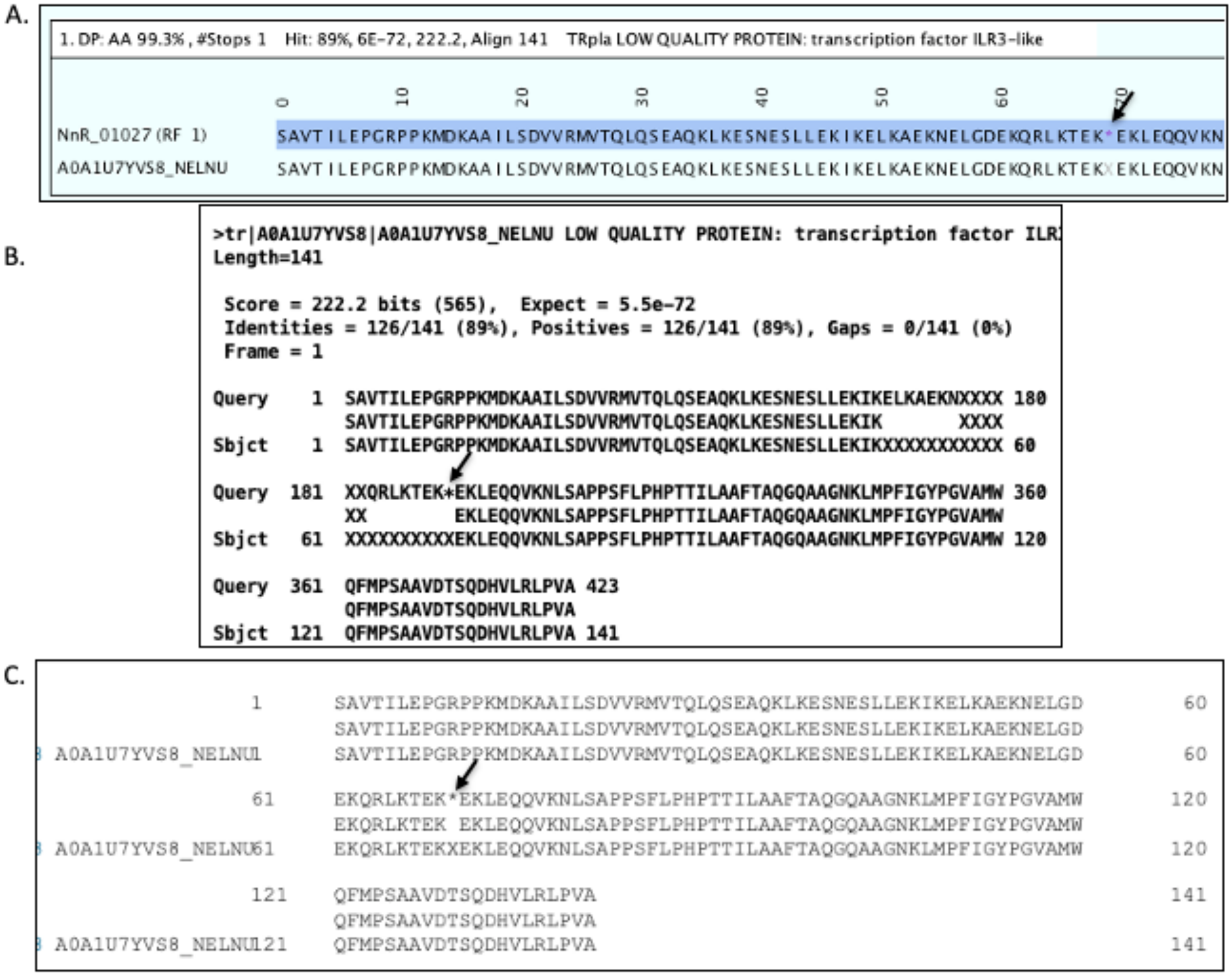

**Figure S11. Sequence region of hit with stop codon.** (A) TCW DP alignment, which shows the stop codon before location 70. (B) DIAMOND alignment using Find Hits, where the output confirms the stop codon. (C) UniProt website alignment, where the BLAST output confirms the stop codon.

#### 2.1.4 Search parameter defaults

RunSingleTCW provides the ability to search with BLAST [2] or DIAMOND [3] for finding the annotation hits. The default search program is DIAMOND since it is fast and freely available. Table 1 shows the parameters that TCW sets to override the respective search program defaults.

**Table 1:** Defaults for BLAST 2.12.0+ and DIAMOND v2.0.11.149

| | BLASTP | | DIAMOND | |
|---|---|---|---|---|
| **Parameter** | **Program defaults** | **TCW defaults** | **Program defaults** | **TCW defaults** |
| evalue | 10 | 0.0001 | 0.001 | - |
| max-hsps | Number found | 1 | 1 | - |
| max-target-seqs | 500 | 25 | 25 | - |
| masking | False (soft) | - | true | false |

To provide an idea of the different hit results between BLAST and DIAMOND, and DIAMOND with and without masking, these were compared as follows: 8871 transcripts were exported from the DIAMOND annotated sTCW_OlR that had a SP-plant hit with E-value ≤ 1E-60 and similarity ≥ 50% (found with Basic

AnnoDB Hit). The transcripts were used to create sTCW_test and annotated with the 3 parameters sets shown in Table 2.

**Table 2: Overview results from BLAST and different DIAMOND parameters**

| Parameters | Unique | Total | Total Avg %Sim | #Seqs with hit | Best Hit (Rank=1) | | | ORF Exact | ORF Multi-frame |
|---|---|---|---|---|---|---|---|---|---|
| | | | | | Avg %Sim | Cover[b] ≥ 50 | Cover[b] ≥ 90 | | |
| BLAST[a] | 23,794 | 132,270 | 51.5 | 8,869 | 79.3 | 71.7 | 12.5 | 726 | 585 |
| DIAMOND | | | | | | | | | |
| --masking 1 | 22,884 | 112,653 | 54.5 | 8,859 | 78.8 | 72.1 | 13.4 | 1,030 | 603 |
| --masking 0 | 23,119 | 115,976 | 53.8 | 8,871 | 79.1 | 73.9 | 14.2 | 1,120 | 619 |

[a] Uses BLAST TCW defaults as shown in Table 1. If sTCW_OlR had been annotated with a BLAST hit file, it is likely that it would have sequence hits that DIAMOND does not find.
[b] Cover ≥ N% is the percent of sequence-hits where the best hit has similarity ≥ N% and hit coverage ≥ N%.

There are more DIAMOND hits without masking compared to with masking, which increases the probable true positives (90 more exact hits) but also increases the possible false positives (16 more multi-frames). The user should experiment with the parameters to find the best for their dataset.

#### 2.2 ORF finding algorithm

The TCW ORF finder is run after the annoDB hits are added to the database. The runSingleTCW Options panel allows the user to set N and M to define a *good hit*, and whether to use alternative start sites (ATG is the default). The algorithm is as follows:

1. If the sequence best hit has a *good hit* (E-value < N or similarity ≥ M), its frame is automatically used. In order to determine the best coordinates:
   a. If the hit region starts and ends exactly at an ATG and stop codon, the hit coordinates are used.
   b. Otherwise, the assigned stop codon is the first downstream stop codon if one exists, else the end of the sequence is used. For the ATG codon, candidates ORFs are created from all upstream possibilities. The best candidate is selected using the sort rules below.
2. Otherwise, the best ORF for the sequence is found by first finding the best ORF per frame and then the best of the 6-frame ORFs using the sort rules.

Sort Rules: Select the best ORF as follows:

1. If the ORF has a *good hit*, select it.
2. Compute lgLen= log ratio of $length_1$ and $length_2$ ($length_1$>$length_2$) and
   lgMK= log ratio of $Markov_1$ and $Markov_2$ ($Markov_1$>$Markov_2$).
   a. If the lgLen > N and at least one ORF has length ≥ 100, select the longest length ORF.
   b. If the lgMK > M and at least one ORF has Markov ≥ 10.0, select the best Markov ORF.
   c. If the lgLen > N, select the longest length ORF.
   d. If the lgMK > M, select the best Markov ORF.
3. If one of the ORFs has both ATG and stop codons and the other does not, select the ORF with ends.
4. Select the ORF with a poor hit or the longest ORF.

Heuristics: ORF finding is complicated by sequences with stop codons in the hit region or have hits to multiple frames. Since the resulting ORF cannot have an in-frame stop codon, the assigned ORF coordinates are the longest hit region that does not contain a stop. TCW will only use the hit information of a multi-frame sequence if the best hit and sequence have coverage ≥ 40% and similarity ≥ 40%. Due to this heuristic, not all sequences with a hit use the hit frame; however, the hit frame may still be used if it passes the sort rules.

5th –order Markov model: The code from TransDecoder v5.5.0 [4] was translated from Perl to Java to be used within TCW. The TCW Markov algorithm is trained on the 2000 longest unique hit regions, where the hit regions come from the *good hits*.

The ORF finder enters information into the TCW Remark to provide information about the ORF, for example, a remark of "!Lg" means it is not the longest ORF (the TCW remark can be searched with Basic Sequence). The ORF finder outputs a set of files as listed in Table 3. It also writes statistics to the log file, as shown in Figure S12.

**Table 3. ORF finder output files.**

| File Name | Description |
|---|---|
| allGoodORFs.pep.fa | ORF for the 6-frames per sequence that pass one of the following: (i) selected ORF, (ii) ORF ≥ 900bp, and (iii) ORF with good Markov score[1]. |
| allGoodORFs.scores.txt | The scores for the allGoodORFs.pep.fa. |
| bestORFs.pep.fa | The translated CDS for the best ORF per sequence. |
| bestORFs.cds.fa | The nucleotide CDS for the best ORF per sequence. |
| bestORFs.scores.txt | The scores for the best ORF per sequence. |
| scoreMarkov.txt | The 5th order Markov model used for training. |
| scoreCodon.txt | The codon usage matrix. |

[1] The Markov good score as defined by TransDecoder, which is described below.

```
ORF Stats:   Average length 587
      Has Hit              36,505  (75.6%)    Both Ends     12,388  (25.7%)    Multi-frame     7,569  (15.7%)
      Is Longest ORF       29,425  (61.0%)    ORF>=300      24,164  (50.1%)    Stops in Hit    4,660   (9.7%)
      Markov Best Score    34,515  (71.5%)    ORF=Hit       15,212  (31.5%)    >=9 Ns in ORF    602    (1.2%)
      All of the above     21,162  (43.8%)      with Ends    3,446   (7.1%)

   Additional ORF info                        For seqs with hit   36,510  (75.6%)    ORF=Hit with Ends   3,446   (7.1%)
    One End              36,133  (74.9%)       Both Ends           9,322  (25.5%)     ORF>=300           3,113  (90.3%)
    Markov Good Frame    36,107  (74.8%)       Markov Good Frame  30,011  (82.2%)     Markov Good Frame  3,290  (95.5%)
    ORF=Hit              15,212  (31.5%)       Markov Best Score  28,408  (77.8%)     Markov Best Score  3,198  (92.8%)
    ORF~Hit               5,641  (11.7%)       Is Longest ORF     23,834  (65.3%)     Is Longest ORF     2,623  (76.1%)
    ORF>Hit              14,806  (30.7%)       Longest & Markov   21,162  (58.0%)     Longest & Markov   2,540  (73.7%)
      with Ends           2,504   (5.2%)       Not hit frame                    5     Sim>=90            3,203  (92.9%)

   Frame: 3(17.5%)  2(16.6%)  1(17.5%)  -1(16.4%)  -2(15.3%)  -3(16.8%)
```

**Figure S12. ORF summary for sTCW_OlR.** The ORF summary in the file logs/anno.log. The top portion is also in the Overview.

In Figure S12, the 'ORF=Hit' is the number of ORFs that use the hit coordinates and the 'ORF=Hit with Ends' is the number that also have an ATG and stop codon (Exact hit). There are two references to Markov:

- Markov Best Score refers to the ORF having the highest Markov score compared to the ORFs found for the other five frames, e.g. frame 1 below has the best Markov score.

```
Length: 576   ORF=Hit
>  1 ORF:  516 (   40 ..  555 )  Markov: 24.29  Codon: 24.33   #Stops  2  EV,WG 3E-91,100%
  -2 ORF:  264 (  461 ..  198 )  Markov: -2.88' Codon:  4.88   #Stops  6
   3 ORF:  111 (    3'..  113 )  Markov: -4.42' Codon:  0.27   #Stops 15
   2 ORF:  135 (  242 ..  376 )  Markov: -4.58' Codon:  0.15   #Stops  7
  -1 ORF:   72 (  405 ..  334 )  Markov: -3.82' Codon:  3.93   #Stops 10
  -3 ORF:   54 (  481 ..  428 )  Markov: -5.59' Codon:  1.97   #Stops 14
```

- Markov Good Frame refers to a score that is positive for the first frame and is better than the score of the other five frames of the ORF, e.g. the following has a good Markov score, where the 6 scores are for offsets 0, +1, +2, -0, -1, -2.

```
Markov   24.29  -39.24  -72.41  -40.11  -15.48  -79.90 (Pos & best) Good score
```

#### 2.2.1 Compare TCW ORF-finder with TransDecoder v5.5.0

The purpose of these two software packages is different, so they have different approaches. A salient purpose for TransDecoder (TD) is to output all potential ORFs, whereas the main purpose for TCW is to select the best ORF per sequence. TD has support for analyzing transcripts that have genome sequence, whereas TCW targets non-model organisms. The following are the major differences in computing ORFs:

1. TCW computes the 5$^{th}$-order Markov model from the 2000 longest unique protein hit regions. TD finds all ORFs over a minimum length (default 300nt) and computes the 5$^{th}$-order Markov model from the 500 longest unique ORFs.
2. TCW compares the transcripts to the annoDBs. TD produces a longest_orfs.cds file of all candidate ORFs, which the user compares to the protein databases.
3. TCW assigns the start and stop coordinates in relation to the hit coordinates whereas TD does not use the hit information when selecting the start and stop of the ORF.

TCW and TD will have slightly different Markov scores since TCW uses the hit regions to train and TD uses the longest ORFs. The scores are shown in the TCW bestORFs.scores.txt file and the TD longest_orfs.cds.scores file. When given the longest_orfs.cds file as input to the TCW ORF finder (i.e. to be used instead of the hit regions), there is still a slight difference because Java has more precision than Perl; the following shows an example of the Markov scores for the 6-frames of an ORF:

```
TCW  77.45  -19.07  -11.84   36.84   8.28  -34.82
TD   78.55  -19.17  -11.34   37.15   5.78  -36.28
```

To compare the ORF results of the two programs, datasets were created from the Osj (*Oryza sativa*) and NnR (*Nelumbo nucifera)* sequences. Each dataset had 5000 sequences with a *perfect hit* to TR-plants (i.e. 100% similarity, 100% hit coverage, ATG and stop codons, no multi-frame); this makes the assumption that these are the correct ORFs. The datasets characteristics are shown under the "TR-plants" columns in Table 4, where the second column shows how many of the 5000 per dataset also have a SP-plants hit.

TCW was used to build a database from each set of 5000 sequences; they were annotated with SP-plants using DIAMOND. TransDecoder was used to analyze the datasets with the option of using hit results, where the hit results were produced by using DIAMOND with the TCW defaults to compare longest_orfs.pep file to SP-plants. The datasets created have the characteristics shown in Table 4.

**Table 4. Characteristics of test datasets with 5000 sequences with perfect TR-plant hits**

| | **TR-plants[1]** | | | | **TCW[2]/TD[3] SP-plants** | | | | | |
|---|---|---|---|---|---|---|---|---|---|---|
| | **Perfect TR hit** | **Has SP Hit** | **Not Longest ORF** | **Not Best Markov** | | **#Seqs** | **%With SP Hit** | **Avg %Sim[4]** | **Cover >=50[4]** | **Stop Codon in Hit** |
| **Set1 Osj** | 5000 | 5000 | 768 | 28 | TCW | 5000 | 100.0 | 66.6 | 100.0% | 55 |
| | | | | | TD | 13,950 | 35.8 | 66.6 | 99.6% | -- |
| **Set2 Osj** | 5000 | 3000 | 1001 | 58 | TCW | 5000 | 70.1 | 55.0 | 53.8% | 48 |
| | | | | | TD | 13,866 | 25.2 | 55.1 | 53.8% | -- |
| **Set3 NnR** | 5000 | 3000 | 136 | 367 | TCW | 5000 | 74.4 | 60.0 | 69.2% | 98 |
| | | | | | TD | 8006 | 46.2 | 60.0 | 69.2% | -- |

[1] The sequences selected from sTCW_Osj (Set1 and Set2) and sTCW_NnR (Set3).

[2] Hit results from the 5000 NT sequence file against SP-plants, used by TCW.

[3] Hit results from the longest_orfs.pep against SP-plants, used by TD. The longest_orfs_pep is produced by TD from the input NT sequences, and may have multiple translated frames per NT sequence.

[4] Rank=1: the Avg %Sim and Cover>=50 as defined for the Overview.

The predicted ORFs results from TCW and TD were compared to the results from the perfect TR_plants ORFs (assumed correct), where the results are shown in Table 5. The TransDecoder does not call an ORF for every sequence, where that number is shown in the "None" column; statistics were not computed for TCW when there was no TransDecoder ORF.

**Table 5. Counts of TCW and TD differences[1(2)] from the correct TR-plant set**

| | | *None* | *Frame* | *Start* | *End* | *Olap* |
|---|---|---|---|---|---|---|
| **Set1** | | | | | | |
| | **TCW** | 0 (0) | 0 (0) | 311 (0) | 0 (0) | 0 (0) |
| | **TD** | 11 (11) | 9 (7) | 879 (0) | 0 (0) | 0 (0) |
| **Set2** | | | | | | |
| | **TCW** | 0 (0) | 21 (21) | 262 (64) | 0 (0) | 2 (0) |
| | **TD** | 84 (84) | 19 (17) | 746 (246) | 0 (0) | 0 (0) |
| **Set3** | | | | | | |
| | **TCW** | 0 (0) | 10 (10) | 167 (4) | 0 (0) | 6 (2) |
| | **TD** | 97 (97) | 22 (22) | 397 (87) | 0 (0) | 3 (2) |
| **Total** | | | | | | |
| | **TCW** | 0 | 31 | 740 | 0 | 8 |
| | **TD** | 192 | 50 | 2022 | 0 | 3 |

[1] *None*: No ORF called. *Frame*: wrong frame. *Start* and *End*: have the correct frame, but wrong start or end. *Olap*: does not overlap the correct coordinates.
[2] The number in parenthesis is the number of incorrect ORFs that did **not** have a hit.

The biggest difference is in the start coordinate, which is not surprising since there is a choice between upstream ATGs, the end of the sequence if there is no intervening stop codons, or the coordinate before the first upstream stop. The reason that TCW gets many of the start coordinates correct is that when there is a similar exact SwissProt sequence, it uses the corresponding coordinates. Conversely, it sometimes gets it wrong because the SwissProt hit has a different start codon then the TrEMBL hit; for example, Figure S13 shows where the TR protein has a different start coordinate than the SP protein. The "Olap" column shows the downside of using the hit directly, as 6 of the 8 TCW wrong overlap coordinates has a SwissProt hit in a different region of the sequence from the TrEMBL hit. Though the TR-plants perfect hits are assumed to be correct, the protein may have been created from the transcripts that they hit; therefore, there could be cases were the TCW or TD results are actually the correct ones.

For the TCW results, the TCW training set was used. However, the training set can make a difference, so it was run again on Set3 with the TD training set (file longest_orfs.cds.top_500_longest). The following is the number of incorrect results with the TD training set, and the number in parenthesis compares the TCW training set:

Frame: 18 (10) Start: 178 (167) Olap: 8 (6)

In summary, for this dataset, the TCW training set provided better results.

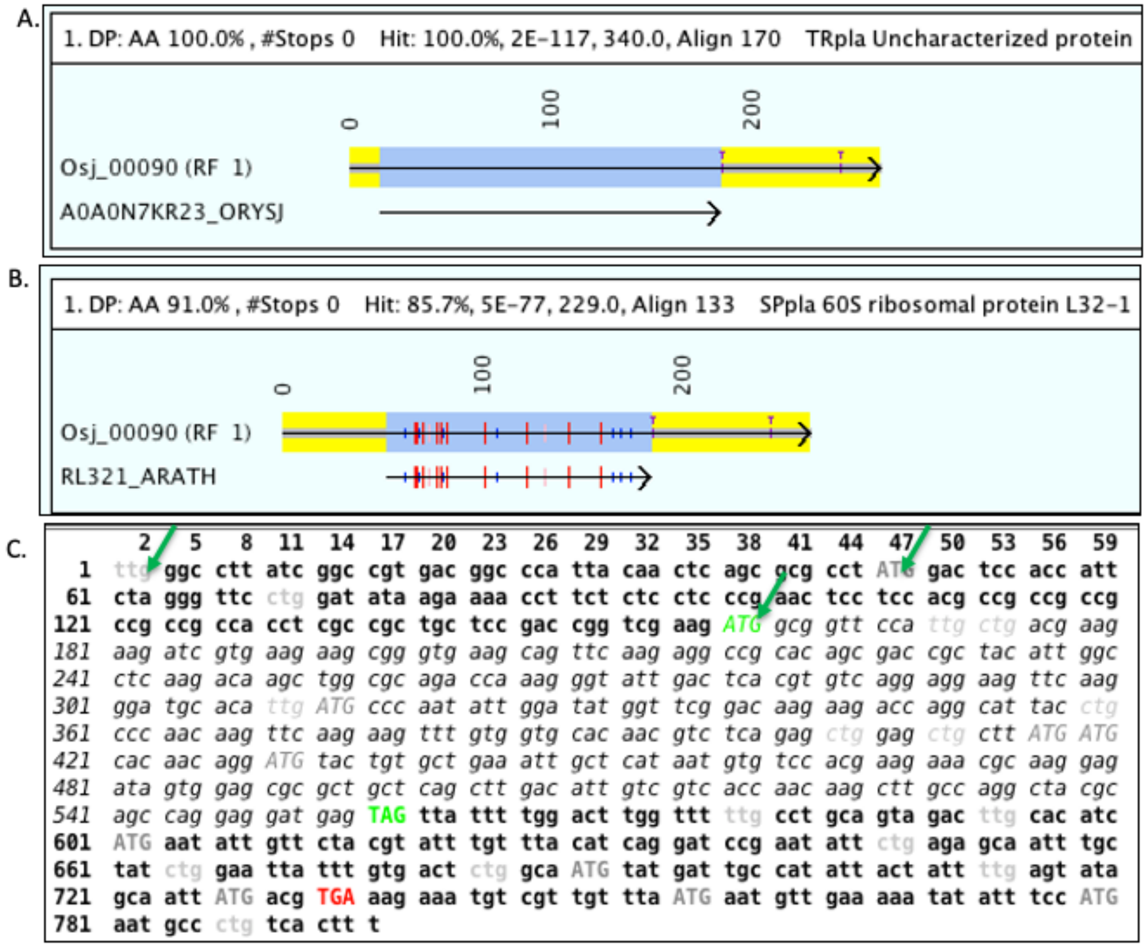

**Figure S13. Different start coordinates.** (A) TCW alignment to a TrEMBL hit. The blue highlighted region is the hit region and the yellow regions are the UTRs. (B) TCW alignment to a SwissProt hit. (C) TCW frame view of the ORF, which starts at position 159 (3rd arrow). The italics is the location of the SwissProt hit. The TrEMBL hit starts the ORF at the ATG a position 47 (2nd arrow). TransDecoder started the ORF at the beginning of the sequence (1st arrow). All three datasets used the same stop codon.

***Reproduce comparison***

- Using viewSingleTCW, create subsets:
  - Set1: Use sTCW_Osj. Basic Hit Filter: Rank=1, AnnoDBs: SP-plants, 1E-30, %Sim ≥ 50, %Hit ≥ 50. Display the results in the Seq Table.
  - Set2: Use sTCW_Osj. Seq Filter: BestBits, AnnoDBs: TR-plants, %Sim ≥ 100, %Hit 100%. Reverse the order of sequences in the Seq Table based on seqID.
  - Set3: Use sTCW_NnR. Same as set2, but do not reverse the order.

  For all three, export the table with columns:
  `#Seq-ID TCW-Remarks #SwissProt BS-Hit-ID BS-DB-type BS-Taxo BS-%Sim BS-%HitCov`

- To create setN.fa (N=1,2,3): Input the exported table into the script found in the TCW source code scripts/V4subsetORF.java. It filters for "ORF=Hit+", not “Multi”, BS-DBtype=”TR”, BS-Taxo=”plants”, BS-Sim=100, BS-HitCov=100 (these last 4 are redundant for set2 and set3). For set2 and set3, include 2000 sequences that have #SwissProt=0. The output is a file of sequences seqN.fa.

- For the TCW Results: Create a sTCWdb with input seqN.fa (N=1,2,3). Annotate with SP-plants using DIAMOND with default parameters. With viewSingleTCW, export a file named tcwSet.tsv with the following columns:
  `Seq-ID  #SwissProt  BS-Frame  BS-Start  BS-End`

- For the TD results: the following commands were executed (N=1,2,3):
  1. `>./TransDecoder.LongOrfs -t setN.fa`
  2. Create a sTCWdb with input longest_orfs.pep. Annotate with SP_plant using DIAMOND with default parameters. Move the resulting search file hitResults/tdN_SPpl.tab to the TransDecoder directory.
  3. `>./TransDecoder.Predict --retain_blastp_hits tdN_SPpl.tab --single_best_only -t setN.fa`
  4. `>grep “>”setN.fna.transdecoder.cds >TD.results`

  The setN.fna.transdecoder.cds header lines were saved to the TD.result file since they contain information about the ORF.

- For the (probably) correct results: For sTCW_Osj and sTCW_NnR, the files trFullOsj.tsv and trFullNnR.tsv were created by exporting the Seq Table with the following columns:
  `Seq-ID  SeqLen  BS-Frame  BS-Start  BS-End`

- The comparison script can be found in the TCW source code file scripts/V4cmpORF.java. It compares the TCW and TD ORF results with the TR-plants ORFs, which outputs a file of the differences and a file of remarks. Using runSingleTCW, the remarks were loaded in sTCW_setN as User Remarks (see §2.5 below); viewSingleTCW was used to inspect the results.

In order to get the exact same results, it would be necessary to use the UniProt SwissProt and TrEMBL plant proteins from Dec-2021.

#### 2.3 GC content

The GC content per sequence is the count of the G and C bases in the sequence divided by the length of the sequence. The number of N’s in a nucleotide sequence is also computed. These numbers are stored in the database for each sequence for display in viewSingleTCW. For the overview, the total percent GC and the CpG O/E [5] is computed for the 5’UTR, 3’UTR and CDS.

#### 2.4 GO levels

Figure S14A shows the QuickGO [6] graph for GO:0000166 and Figure S14B shows the corresponding TCW view. In the TCW view, the levels are shown in the first column. As discussed in the manuscript, levels are not a valid GO relation since a GO can be on multiple levels, however, it provides a convenient way for users to understand a GO terms approximate location in the graph.

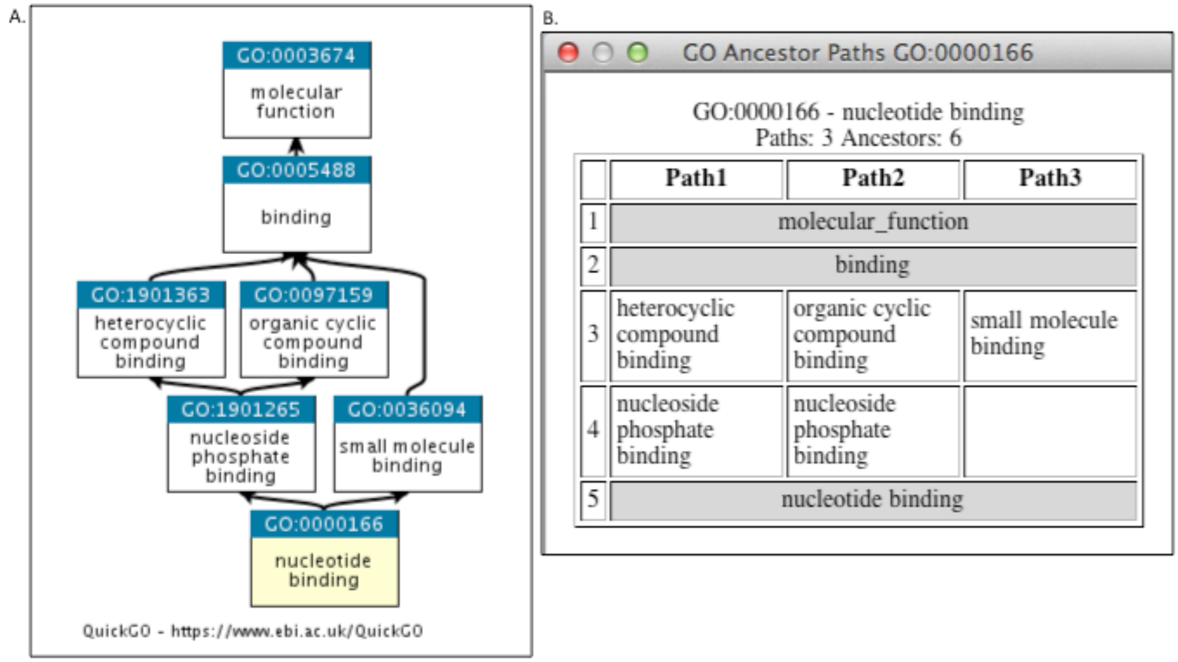

**Figure S14. The paths of a GO term.** (A) The QuickGO view of GO:0000166. (B) The TCW view of GO:000166. TCW assigns level 5 to GO:0000166 since it has maximum level.

***To reproduce the Figure S14 graphs:***

- Figure S14A: link https://www.ebi.ac.uk/QuickGO/term/GO:0000166.
- Figure S14B: From the Basic GO Annotation panel, enter GO:0000166 in the Basic GO search box, select BUILD, select the GO term from table, select Show Ancestor Paths from the Show… dropdown.

#### 2.5 Load Remarks and Locations

At the bottom of the runSingleTCW interface (Figure S3) is the Load Remarks and Locations option. Using this feature, the user can load locations (i.e. chromosome, start, end), which are entered into the sTCWdb as columns that can be viewed by viewSingleTCW in the Sequence table. The user can also load a file that contains one or more rows of sequence names followed by a remark to be associated with the sequence as a User Remark. These can be queried and viewed in the Basic Sequence panel and viewed as a column in the Sequence table. If there are different types of external data to be added, the user should enter the data in "keyword:value" format, then a given type of data can be queried by its keyword in Basic Sequence. This simple approach is a powerful way to keep external information associated with each sequence; the user has complete control of the content of the remarks.

### 3 runDE: differential expression and GO enrichment

Figure S15 shows the interface for computing differential expression. The fourth option of the EXECUTE section allows the user to input the p-values, which would be necessary if the desired method is not coded in the R language. After computing all desired DE p-values, the Select p-value column can be used to select a single DE column or all of the columns (All p-value columns); then GO enrichment script can be run on the selected column(s).

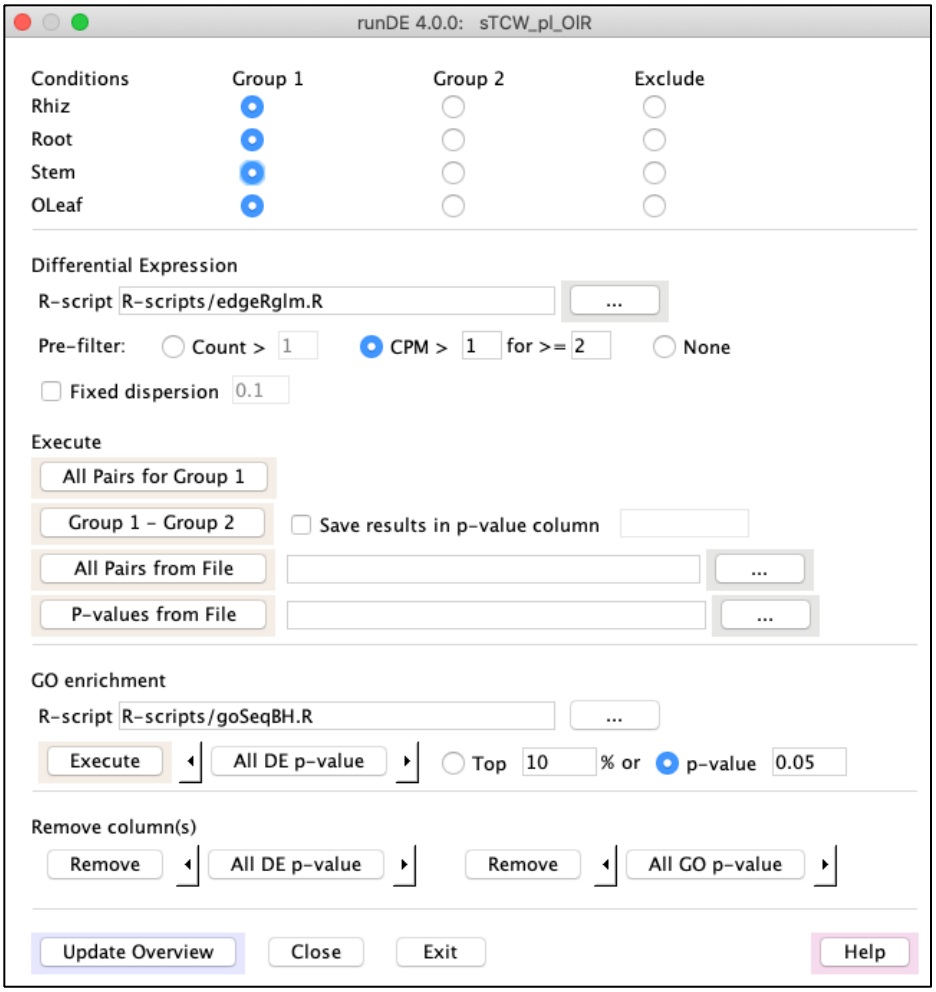

**Figure S15. The runDE graphical interface.** For the sTCW_OlR, these four tissues were selected and All Pairs for Group 1 was executed, which runs all pairs and automatically creates a column name for each.

#### 3.1 R scripts for Differential Expression

The TCW runDE program writes the appropriate data to R data structures (i.e. a matrix of sequence names and replicate counts for both conditions, etc.), then runs the R script, which uses the data from the structures and puts the results into another structure, which TCW reads and enters into the sTCWdb. R scripts are supplied for edgeR [7] and DESeq2 [8], which provides a template to modify or create new R scripts.

#### 3.2 R scripts for GO enrichment

GO enrichment runs a user-supplied script in a similar manner as discussed above for DE. A script is supplied for GOseq. Before GOseq can be executed, the DE p-values must be computed and the GOs associated with the sequences must be in the sTCWdb. GOseq takes as input the sequences that are significantly DE between conditions based on a given p-value cutoff and the list of GOs assigned to each sequence. To be more specific, TCW creates the binary vector of all sequences that have at least one hit with GOs where the $i^{th}$ value will be 1 if the $i^{th}$ transcript has DE p-value < cutoff (default 0.05), otherwise, it is 0. TCW also creates N lists of GO terms where the $i^{th}$ list is all direct and indirect GOs associated with the $i^{th}$ sequence. These two data structures are used by the GOseq script, which produces a p-value for each GO term.

### 4 Results using REVIGO and WeGO

The OlR dataset was queried for all levels of the GOs that had a GOseq p-value < 1E-06 for RhRo (rhizome-root), RhSt (rhizome-stem) or RhOl (rhizome-leaf), which resulted in 162 GOs. The input to REVIGO [9] was the GOs and their DE transcript count (see S1 Suppl §2.1.6). Each ontology was reduced separately, where the cellular process was reduced to 30 GO terms as shown in Figure S16. WeGO [10] displays the its level 3 GO terms, as shown in Figure S17.

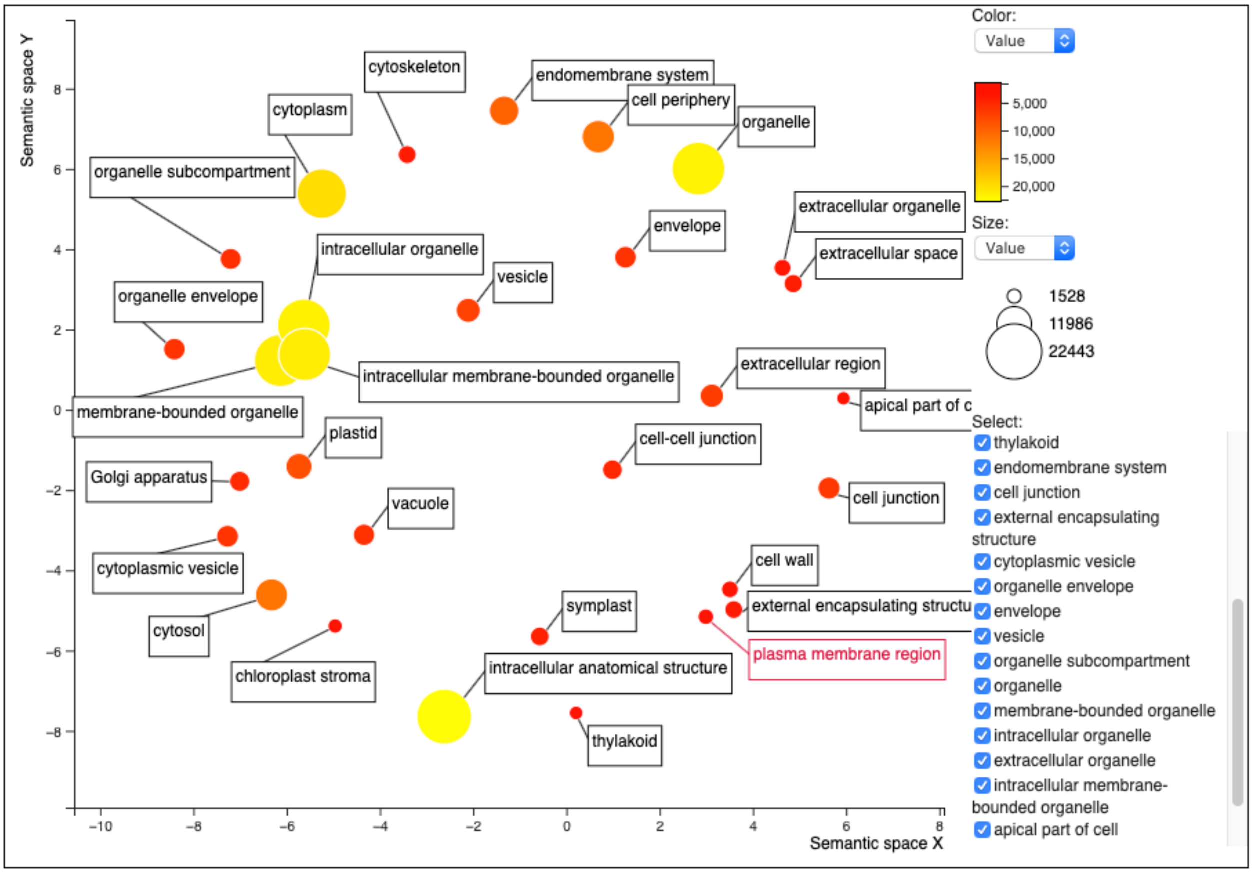

**Figure S16. REVIGO.** Cellular component enriched GOs for rhizomes.

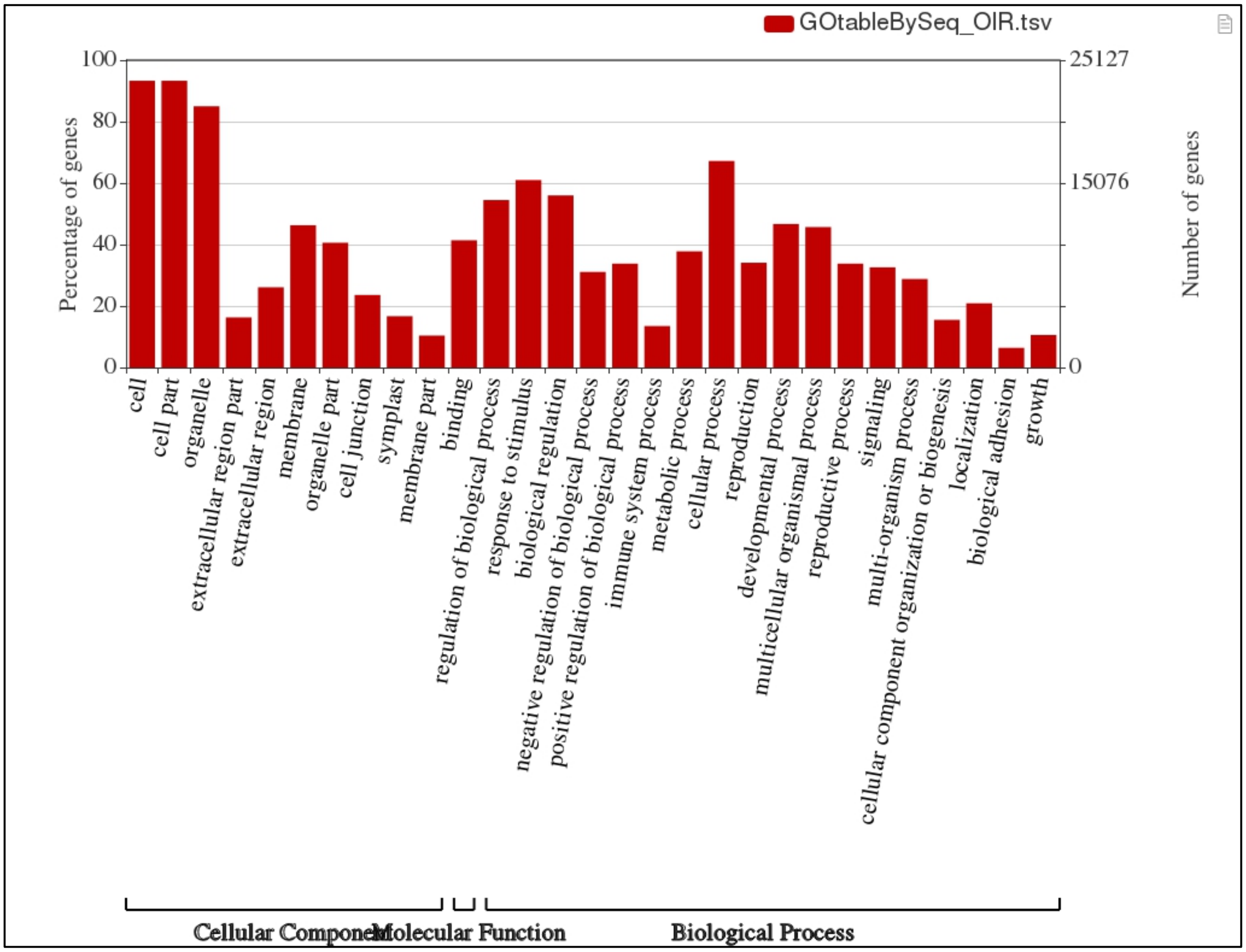

**Figure S17. WeGO.** Biological process over-represented level 3 GOs for rhizomes.

***To reproduce these graphs:***

- Basic GO Annotation:
  - Enrich: checkbox, cutoff 1E-06, P-value RhRo, RhSt, RhOL with Any, #Seqs DE.
  - BUILD.
- For REVIGO (Figure S16):
  1. Select columns GO ID and #Seqs. Select Export table from the Table… dropdown.
  2. Enter the resulting file into http://revigo.irb.hr
  3. On the REVIGO interface, set:
     a. "How large…" to "Small".
     b. "the values associated with GO terms represent" with "higher is better".
     c. "Species" to "Oryza Sativa".
  4. The GO database and mappings were from November 16, 2021
  5. When the results are displayed, select "Cellular Process" and "Scatterplot&Table". A snapshot of the results was used.
- For WeGO (Figure S17):
  1. Select dropdown option SeqID with GO from Export….
  2. Enter the resulting file into http://wego.genomics.org.cn.
  3. Use "Native format". The latest GO at the time of this writing was Nov 2018.
  4. On the resulting page, select "Graph". Export the image.
