## Supplementary material for "Transcriptome computational workbench (TCW): analysis of single and comparative transcriptomes": S3 Suppl

### Build a multiTCW database

**Sections**

The purpose of this supplement is to provide additional information about the algorithms and parameters. ViewMultiTCW is used to demonstrate the results, where Figure 1 shows the interface features.

mTCW_pl and mTCW_rhi are referred to in this document. The mTCW_pl was discussed in the main text. The mTCW_rhi was created using the OlR subset and the original NnR dataset [1], which has 26,285 sequences and the same expression counts files as the OlR datasets, i.e. rhizome, root, stem and old leaf.

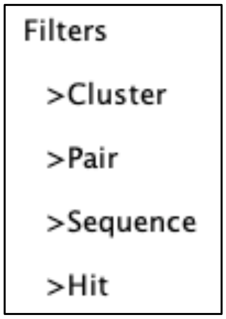

**Figure S1. The panels shown on the left of viewMultiTCW**.

Cluster Filter provides filters on clusters, which results in the Cluster Table. A cluster row can be selected to view the MSA, sequences, or pairs.

Pair Filter provides filters on pairs, which results in the Pair Table. A pair row can be selected to view the pairwise alignment, sequences or clusters.

Sequence Filter provides filters on sequences, which results in the Sequence Table. A sequence row can be selected to view the details, clusters or pairs. A set of rows can be selected to view the MSA.

Hit Filter provides limited filters on hits, which results in the Hit Table. A hit row can be selected to view the hit sequences, or pair alignment of hit protein with the hit Sequences.

### 1 runMultiTCW interface

The runMultiTCW interface is shown in Figure S2. Step 1 loads the sTCWdbs. Step 2 runs the self-search and loads the pairs. Step 3 computes the clusters. Step 4 adds statistics to the pairs and clusters.

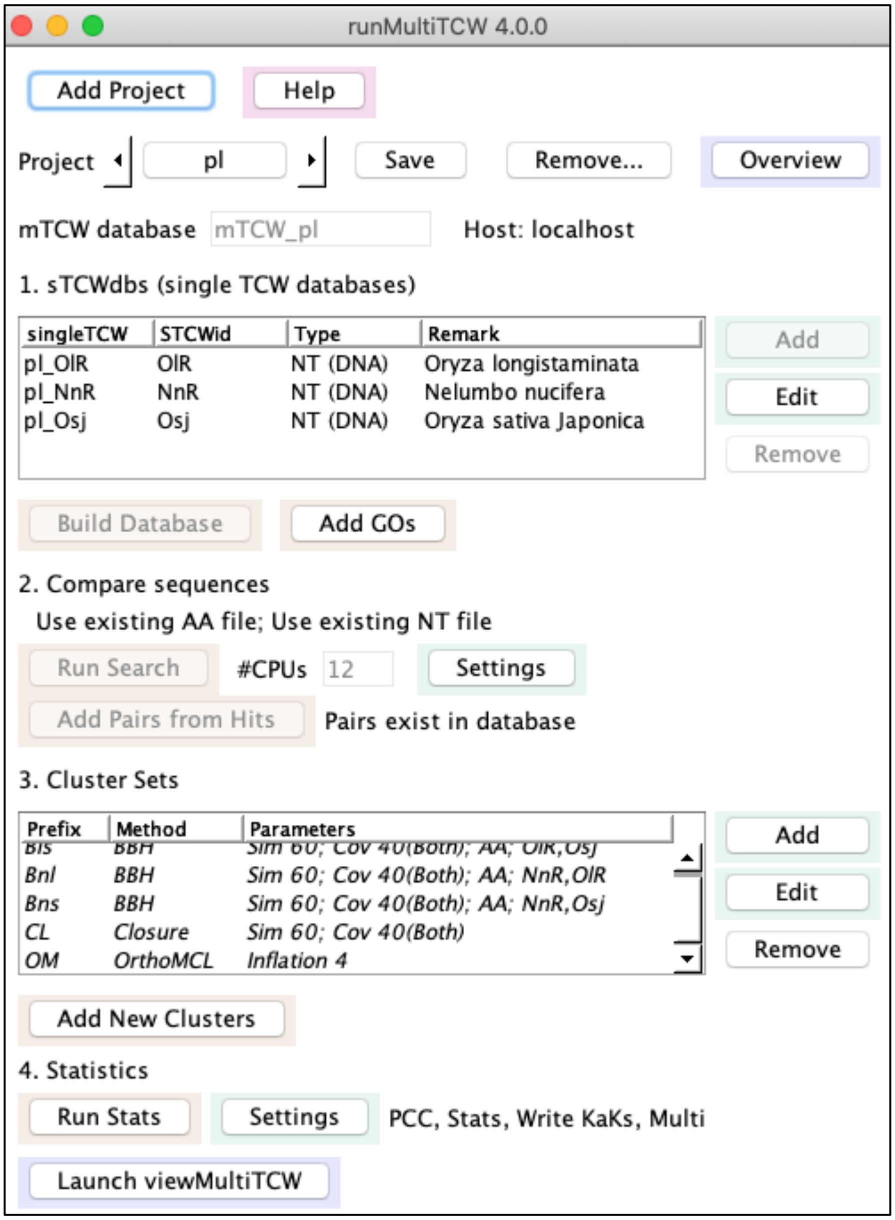

**Figure S2. The runMultiTCW main interface.** The disabled buttons indicate that the function has been executed. The italics clusters indicate they are in the database. Additional clusters can be added, which is why the Add New Clusters is not disabled.

### 2 Build Database and Add GOs

For each sTCWdb listed in runMultiTCW, only the top 5 annotations from each annoDB are loaded, along with the Best Anno and Best GO if they are not in the top 5. For nucleotide sTCWdbs, the ORF coordinates and translated ORFs are loaded. The GC and CpG content are computed for the 5'UTR, CDS, 3'UTR and full nucleotide sequence, as discussed in §6. If there are expression levels for the input sTCWdbs, the TPM and DE values are loaded, as discussed in §7. The Add GOs function transfers the GO information associated with the hits in the database from the sTCWdbs; this can be run anytime after the Build Database.

### 3 Search and add pairs

The Run Search runs the AA (amino acid) and optional NT (nucleotide) self-search on a file containing all sequences in the database. For the AA search, either DIAMOND [2,3] or BLAST[4] may be used. The NT search uses BLAST.

#### 3.1 Search parameters for amino acid sequences

Moreno-Hagelsieb and Latimer [5] showed that the optimal BBH hits were obtained with the BLAST parameters '-soft_masking true -use_sw_tback', where the second parameter causes BLAST to compute locally optimal Smith-Waterman alignments; these parameters are used for the TCW BLAST defaults (however, sometimes the -use_sw_tback causes blastp to crash).

Table 1 shows the results from 9 DIAMOND runs with different parameters using mTCW_rhi (74,557 sequences) and mTCW_pl (144,745 sequences):

**Table 1. Comparing various DIAMOND parameters.**

| DB | Parameters[1] | | | Pairs | | | | Clusters | | |
|---|---|---|---|---|---|---|---|---|---|---|
| | Mk | Sen | Cov | Total Pairs | % Share Descr | %Similarity | % Coverage | BBH Bnl | Closure | Ortho MCL |
| **mTCW_rhi** | | | | | | | | | | |
| **A** | | | | 339,758 | 83.2 | 47.3 | 56.4 | 4,382 | 11,788 | 11,291 |
| **B** | Y | | | 363,280 | 82.5 | 46.1 | 58.4 | 4,452 | 12,082 | 11,462 |
| ***C*[2]** | *Y* | | *Y* | *279,474* | *83.2* | *45.3* | *70.7* | *4,491* | *12,090* | *11,696* |
| **D** | | Y | | 444,180 | 78.7 | 44.8 | 53.2 | 4,383 | 11,836 | 11,218 |
| **E** | Y | Y | | 465,151 | 77.8 | 43.8 | 55.6 | 4,498 | 12,127 | 11,362 |
| **F** | Y | Y | Y | 360,532 | 79.0 | 43.1 | 68.9 | 4,534 | 12,122 | 11,680 |
| **mTCW_pl** | | | | | | | | | | |
| **B** | Y | | | 1,277,703 | 84.9 | 56.4 | 71.2 | 5,028 | 26,566 | 20,039 |
| ***C*[2]** | *Y* | | *Y* | *1,056,941* | *85.8* | *56.3* | *79.8* | *5,049* | *26,613* | *20,607* |
| **F** | Y | Y | Y | 1,212,949 | 82.7 | 54.2 | 78.3 | 5,058 | 26,658 | 20,480 |

[1] Mk = –masking; Sen = –sensitive; Cov = –query-cover 25 –subject-cover 25; Y = the parameter was used.
[2] TCW DIAMOND defaults.

For DIAMOND, the TCW defaults are '–masking 0 –query-cover 25 –subject-cover 25' (parameters *C*) which produce less pairs with better coverage, and the number of clusters is similar to the other sets.

By default, DIAMOND sets masking to 1 (on). With masking on, DIAMOND can miss short good hits, e.g. a perfect alignment of two sequences with 27 AA was missed with masking on and found when masking off. Of course, with masking off there will be more false positives, but they will be filtered in the clustering.

The user should experiment to find the best parameters for their dataset; however, there is no perfect set of parameters. The search parameters can be changed using the Settings panel.

Reproduce comparison: The Table 1 results are printed in the log files.

#### 3.2 Add Pairs from Hits

Add Pairs from Hits loads all hits and associated information from the AA and NT tabular files into the database. The bidirectional best hits are recorded in the database for the AA and NT hits when the respective file is loaded. The algorithm for assigning the 'best description' is the same as for clusters (see §4.5), with the exception that if there is no shared hit, the majority HitID is assigned the value '*NoShare'.

### 4 Compute clusters

#### 4.1 BBH

BBH (bidirectional best hit), also called RBH (reciprocal best hit), computes pairs where they are each others best hit. The TCW BBH has two user-defined parameters: the %Similarity (identity; default 60%) and the %Coverage (alignment-length/sequence-length; default 40%), where the coverage requirement can be applied to either or both sequences of a pair (default both). As stated above, the BBH pairs were computed when the AA pairs were loaded from the hit file. The BBH algorithm queries the database for the pairs marked as AA-BBH and filters them according to the similarity and coverage parameters.

##### 4.1.1 Compare Galaxy and TCW BBH

Galaxy BLAST_rbh [6] computes BBH. The TCW BBH and Galaxy BLAST_rbh.py v0.1.11 were compared, which showed the following minor algorithmic differences:

1. TCW rounds the similarity or coverage value, then compares it to the cutoff whereas Galaxy does not. Galaxy uses the BLAST qcovhsp optional column for the coverage whereas TCW uses (alignment-length/sequence-length), where alignment-length the BLAST align column.
2. TCW first selects the BBH pairs, then filters on similarity and coverage. Galaxy first filters on similarity and coverage, then selects the BBH pairs.
3. TCW arbitrarily selects a pair if there is a tie based on E-value and bit-score whereas Galaxy does not select any pair in this case; however, it does have an option to remove redundant sequences which would subsequently result in a BBH pair.
4. TCW performs a complete self-search of all sequences. Galaxy BLASTs the first dataset file against the second dataset file and vice versus. This results in slightly different scores.

With the exception of these differences, the programs produce the same results.

TCW and Galaxy agreed on 800 pairs, where TCW had 853 pairs and Galaxy had 850 pairs. Most of the differences came when A→B→C⇔D in TCW (where → implies one-way best hit and ⇔ is bidirectional); that is, due to the differences stated above, TCW and Galaxy will occassionally identify different bidirectional pairs. The benefit of the Closure method is that it brings ABCD into the same cluster.

Reproduce comparison:

- The Galaxy tar file was downloaded from: https://toolshed.g2.bx.psu.edu/repository?repository_id=d5dd1c5d2070513e.
- Viewing the mTCW_rhi database with viewMultiTCW, the pairs that were in the Closure cluster were viewed and the AA sequences written to file. A script was written to read the file and create an NNU.fa and OlR.fa file of the first 2000 paired sequences.
- TCW:
  - Two singleTCW databases were created from the NNU.fa and OlR.fa sequences.
  - A multiTCW database was created from the two singleTCW. The search program was blastp with no parameters (as Galaxy uses). The BBH clusters were created using Similarity cutoff of 70 and Coverage cutoff of 50 (as Galaxy uses).
  - viewMultiTCW was used to view all pairs along with the BBH column, and the Table... function was used to export the table into a file called PairTable.xls.

- Galaxy: The following command was executed in the Galaxy directory: `./BLAST_rbh.py -a prot -t BLASTp -o galaxy.tsv NNU.fa OlR.fa`
- The TCW PairTable.xls and Galaxy galaxy.tsv files were compared with V4cmpBBH.java in the TCW scripts directory.

#### 4.2 Closure

The Closure algorithm seeds the clusters with the BBH marked pairs that pass the user-supplied parameters, and then pulls in similar pairs. It uses the same two parameters as the BBH cluster algorithm (%Similarity and %Coverage). It requires that all sequences in a cluster to have a search hit and pass the parameters with all other sequences in the cluster.

When Closure is run using the same parameters that were used with BBH, it provides an expansion of the BBH clusters, i.e. showing what other sequences align to the BBH pairs. If run with the coverage parameters set to zero, it performs poorly as it will include very short sequences embedded in long sequences.

#### 4.3 Best Anno hitID or description

The method has 4 parameters: (i) use Best Anno (best annotation) hitID or description, (ii) %Similarity (default 20), (iii) %Coverage (default 20), and (iv) All/Any (default all). If the hit description is used, a cluster can contain multiple different hitIDs. To compare descriptions:

- If the Best Anno description is un-informative (e.g. contains 'uncharacterized protein'), then the sequence can only be part of a cluster with the same hitID.
- Before comparing the descriptions, each is converted to lower case, and if there is text at the end starting with '{', it is removed.

Clusters are formed as follows (the terms 'protein hit' or 'pair hit' indicates a DIAMOND/BLAST result):

- All sequences share the same best hitID or best description.
- Each sequence in the cluster has a protein hit with %similarity ≥N and %coverage ≥M to the representative cluster protein.
- A sequence must have a pair hit with %similarity ≥N and %coverage ≥M to all/any other sequences in the cluster.

This was not used in the mTCW_pl database, as it does not perform as well as Closure or OrthoMCL.

#### 4.4 OrthoMCL

OrthoMCL [7] is designed to find distantly related orthologous sequences using a Markov Cluster algorithm. OrthoMCL requires multiple steps including creating a MySQL database and running BLAST. Using TCW to run OrthoMCL, TCW creates the MySQL database, uses the hits file generated with Run Search as input to OrthoMCL, runs all steps, and enters the results into the mTCWdb. OrthoMCL has one parameter called 'inflation', where the default is set to 4 (the OrthoMCL default is 1.5, but 4 provides tighter clusters). OrthoMCL can result in sequences that do not have a hit with one or more other sequences in a given cluster.

OrthoMCL was downloaded from https://orthomcl.org/common/downloads/ and the mcl program was downloaded from http://www.micans.org/mcl/src/mcl-latest.tar.gz.

#### 4.5 Annotation of clusters

The algorithm for assigning a description and representative hitID to the cluster is as follows:

1. A best hit list is created as follows: the best description (Best Anno) from each sequence is modified and then added to the list if it does not already exist. The descriptions are modified by removing the leading words such as 'putative', converting to lower case, then taking the first two words.
2. All hits for all sequences are checked against the modified descriptions of the best hit list, and the sequences that have at least one hit with the same two first words are counted (numHasHit).

3. The hit list is sorted on {is_good_annotation, numHasHit, E-value, is_SwissProt, number of GOs}. The first hit from the sorted list is assigned as the cluster annotation.
4. If all the sequences in a cluster do not have a hit, the cluster is given a hitID of '*Novel'.

The percentage of cluster sequences that have the description/hitID is stored in the database per cluster.

### 5 Run Stats

#### 5.1 Pairs alignments

For all pairs in clusters that have a hit, each pair is aligned using dynamic programming of the amino acid sequences and the results are mapped to the codons of the sequences, which provides a *codon-based alignment*. The codon-based statistics, listed in Table 2 of the manuscript, are computed on the overlapping regions of the alignment. These statistics can be queried and viewed (e.g. Figure S3 shows some statistics for a pair).

```
CROP:    249 (   83)   Calign:     82    Codon exact:     78 (95.1%)   Amino exact:       79
Full:    249 (   83)   GapOpen:     1    Synonymous:       1 ( 1.2%)   Substitution >0:   1
Hang:      0 (    0)   Gaps:        3    Nonsynonymous:    3 ( 3.7%)   Substitution<=0:   2

OlR_01296     1  atg gag gct tca cgc aag gtg ttc tcg gcc atg ctt ctc atg gtg ctg ctg ctg ctt gca    60   60

Osj_15230     1  atg gag gct tca cgc aag gtg ttc tcg gcc atg ctt ctc atg gtg --- ctg ctg ctt gca    57
-------
OlR_01296    61  gcc acc ggt gag atg ggc ggg ccg gtg atg gtg gcg gag gct cgg acg tgc gag tcg cag   120  120
                       |
Osj_15230    58  gcc act ggt gag atg ggc ggg ccg gtg atg gtg gcg gag gct cgg acg tgc gag tcg cag   117
-------
OlR_01296   121  agc cac cgg ttc aag ggc ccg tgc gcc cac aag gag aac tgc gcc agc gta tgc aac acg   180  180
                                                      *       *
Osj_15230   118  agc cac cgg ttc aag ggc ccg tgc gcc cgc aag gcg aac tgc gcc agc gta tgc aac acg   177
-------
OlR_01296   181  gag ggc ttc ccc gac ggc tac tgc cac ggc atc cgc cgc cgc tgc atg tgc acc aag ccc   240  240
                                                          *
Osj_15230   178  gag ggc ttc ccc gac ggc tac tgc cac ggc gtc cgc cgc cgc tgc atg tgc acc aag ccc   237
-------
OlR_01296   241  tgc ccc tga                                                                       249  300

Osj_15230   238  tgc ccc tga                                                                       246
-------

LEGEND:
        | = Synonymous codon
        * = Nonsynonymous codon
```

**Figure S3. AA, CDS, NT alignment.** Text view of the CDS alignment with synonymous/nonsynonymous shown.

<u>Reproduce Figure S3:</u> From a pair alignment, select AA,CDS,UTR from the Pairwise... option, then select CDS... and show with the defaults.

The aligned sequences are written to files with the gaps removed for input to the KaKs_Calculator [8] and a runKaKs script is written for the user to execute from the command line, followed by using runMultiTCW to read the results into the database. The aligned sequences are split into multiple files, and there is one line per file in the runKaKs script to run the KaKs_Calculator:

```
/Users/cari/TCW/Ext/mac/KaKs_Calculator  -i oTCW1.awt -o iTCW1.tsv -m YN &
```

The YN is the method that can be changed by the user before executing the script. Alternatively, the user can provide one or more KaKs files for input to TCW.

#### 5.2 Multi alignments

The MAFFT [9] program is executed on all clusters of size > 2 to compute the multiple sequence alignment (MSA); occasionally it fails, in which case, MUSCLE [10] is run. For clusters of size 2, the dynamic programming results from the pair alignment are used. The alignment, consensus sequence length and the standard deviation of all aligned sequences are entered into the database to be viewed with viewMultiTCW, as shown in Figure S4.

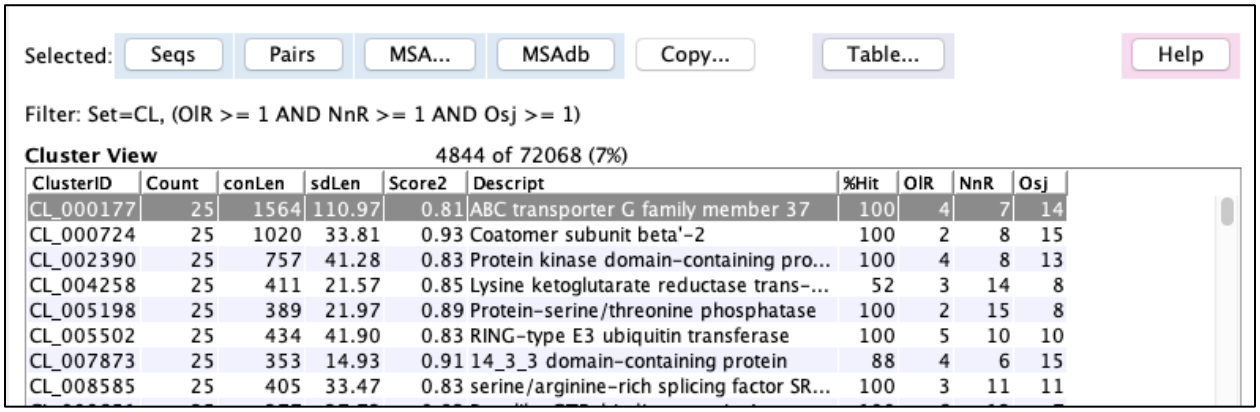

**Figure S4. Cluster table.** The MSAdb displays the precomputed MSA; the MSA... dropdown has other options like computing the MSA with all sequences and the hit sequence. The conLen is the consensus length and the sdLen is the standard deviation of the sequence lengths from the conLen.

By default, the MSA are scored with sum-of-pairs and Wentropy [11], which are referred to as Score1 and Score2. The user can substitute either Score1 or Score2 with methods from MstatX [12], which is included in the TCW package; alternatively, they can provide a file of their own scores or add a statistic to MstatX.

The Wentropy methods was translated from the MstatX C++ code to Java for use within TCW. The cluster score is the 1-(average column score).

The sum-of-pairs score compares each two characters in the column, where there are 22 possible characters (20 amino acids, gap '-', and leading/trailing space ' '). The comparison scores are: ($aa_1$,$aa_2$) is the BLOSUM68 score, (aa,'-') is -4, (aa,' ') is -1, ('-','-') is 0, (' ',' ') is 0. The cluster score is the sum of the columns scores divided by the number of comparisons.

### 6 GC, CpG and Ts/Tv

For each sequence, the CpG Obs/Exp [13] is computed for the 3'UTR, CDS, 5'UTR sequences. The average GC and CpG are computed for the full nucleotide sequence. The overall percent GC and CpG Obs/Exp is computed for the Overview as shown in Figure S5A.

The transition(ts)/transversion(tv) ratio is computed for each aligned pair along with the percent SNPs that are transitions and percent SNPs that are transversions. Using the CDS alignment, the Jaccard index is computed for the GC content, CpG-NT (may cross codon boundaries) and the CpG-Cd (may not cross codon boundaries). Figure S5B and S5C shows the Overview of these results for the BBH OlR-Osj and BBH NnR-Osj pairs.

A. Datasets

| | Average Lengths 5UTR | CDS | 3UTR | %GC 5UTR | CDS | 3UTR | CpG O/E 5UTR | CDS | 3UTR |
|---|---|---|---|---|---|---|---|---|---|
| OlR | 216.5 | 566.6 | 295.9 | 44.0% | 50.1% | 40.3% | 0.761 | 0.772 | 0.616 |
| NnR | 332.3 | 1372.5 | 362.2 | 42.4% | 44.3% | 37.3% | 0.691 | 0.465 | 0.394 |
| Osj | 401.4 | 1260.0 | 539.2 | 49.2% | 52.6% | 40.9% | 0.885 | 0.851 | 0.614 |

B. BBH OlR-Osj

| | Pos1 | Pos2 | Pos3 | Total | | GC | CpG-Nt | CpG-Cd |
|---|---|---|---|---|---|---|---|---|
| Transition | 12.7% | 11.0% | 29.9% | 53.5% | Both | 48.6% | 8.3% | 4.5% |
| Transversion | 13.7% | 10.8% | 22.0% | 46.5% | Either | 50.8% | 9.7% | 5.2% |
| ts/tv | 0.92 | 1.02 | 1.36 | 1.15 | Jaccard | 0.96 | 0.86 | 0.87 |

C. BBH NnR-Osj

| | Pos1 | Pos2 | Pos3 | Total | | GC | CpG-Nt | CpG-Cd |
|---|---|---|---|---|---|---|---|---|
| Transition | 11.1% | 7.6% | 29.3% | 48.0% | Both | 35.2% | 2.1% | 1.4% |
| Transversion | 13.6% | 8.6% | 29.9% | 52.0% | Either | 59.8% | 13.3% | 7.1% |
| ts/tv | 0.82 | 0.88 | 0.98 | 0.92 | Jaccard | 0.59 | 0.16 | 0.19 |

**Figure S5. Overview of the GC, CpG and Ts/Tv.**

Reproduce Figure S5: (A) The top part of the mTCW_pl overview. (B-C) These were obtained as follows: (i) From Pairs Filter, select Cluster Set OlR-Osj. (ii) View Filtered Pairs. (iii) On the resulting table, select option Show Table Stats from the Table.... dropdown. The pop-up display shows the statistics shown. Repeat i-iii for NnR-Osj.

The text pair alignment has an option of showing the CpG sites, as shown in Figure S6. The alignment can also be viewed with annotation synonymous/non-synonymous, amino acid characters, degenerate or ts/tv.

CDS codon align

```
                                          All CpG              By Codon
CROP:     90 (   30)    GC Both:       38    CpG Both:        2    CpG Both:        1
Full:    330 (  110)    GC Either:     56    CpG Either:     14    CpG Either:      9
Hang:    240 (   80)    GC Jaccard: 0.679    CpG Jaccard: 0.143    CpG Jaccard: 0.111

OlR_09567     1  aac aag cgc gtg tgc gat gag gtc gcg atc atc ccg tcc aag cgg atg cgc aac aag atc      60   60
                          x                       x           x   x       |       x
NnR_42769    25  aac aag aag atg tta gag gag gtc gcc att att cct tcg aag cgc ctg ccc aac aag atc      84
-------
OlR_09567    61  gcg ggt tac atc acg cac ttg atg aag cgc                                              90  120
                  x               x                   x
NnR_42769    85  atc gga ttt tgt acc cac ctc atg aag tga                                             114
-------

LEGEND:
        | = both codons have CpG
        x = only one codon has CpG
Note: CpG crossing codon boundaries are not shown.
```

**Figure S6. CDS alignment showing codon-based CpG content.**

Reproduce Figure S6: From a pair, select 5UTR,CDS,3UTR or AA,CDS,UTR from the Pairwise... dropdown, select CDS... which will result in a pop-up of options, select option CpG.

### 7 TPM, DE and PCC

The DE analysis in multiTCW is only effective if the expressions profiles have the same conditions. For input into mTCW, they need to be given the same column names for both the counts and the DE, as shown in Figure S7.

**Counts**

| | Rhiz | Root | Stem | OLeaf |
|---|---|---|---|---|
| NNU | 153,384,951 | 146,651,258 | 146,994,359 | 140,443,622 |
| OlR | 132,552,736 | 127,328,342 | 141,332,137 | 149,461,466 |

**Differential Expression (p-value < 0.001)**

| | RhRo | RhSt | RhOL | RoSt | RoOL | StOL |
|---|---|---|---|---|---|---|
| NNU | 9k | 9k | 8k | 12k | 9k | 3k |
| OlR | 12k | 21k | 22k | 13k | 12k | 8k |

**Figure S7. The TPM and DE from the mTCW_rhi overview.**

The only feature available for the expression profiles is the Pearson Correlation Coefficient (PCC), which is applied to the TPM values for each pair of sequences that have a hit score. Figure S8A shows the results of setting the Pairs Filter to show the BBH pairs with PCC ≥ 0.8. Figure S8B shows a pair with their TPM values. For clusters, there is a column showing the percentage of pairs in the cluster with PCC ≥ 0.8 (not shown).

A. Filter: PCC>=0.8 AND In(BB)

**Pair View** 967 of 389136 (<1%)

| Row # | SeqID1 | SeqID2 | PCC | Descript | AAeval | Align | %Cov1 | %Cov2 |
|---|---|---|---|---|---|---|---|---|
| 1 | NNU_008809 | OlR_06890 | 1.000 | probable aquaporin TIP-type R... | 5.2E-95 | 555 | 73.71 | 100.54 |
| 2 | NNU_010125 | OlR_26010 | 0.9999 | iron-sulfur assembly protein Is... | 1.7E-62 | 309 | 60.23 | 90.35 |
| 3 | NNU_013224 | OlR_47041 | 0.9999 | Expansin | 1.4E-88 | 570 | 59.75 | 100.53 |
| 4 | NNU_016024 | OlR_01012 | 0.9999 | psbP domain-containing protei... | 2.3E-56 | 321 | 51.20 | 100.00 |
| 5 | NNU_002178 | OlR_12920 | 0.9999 | protein CONSERVED IN THE GR... | 3.0E-147 | 966 | 103.21 | 102.88 |
| 6 | NNU_025699 | OlR_45065 | 0.9998 | Na_H_Exchanger domain-conta... | 0.0 | 2496 | 103.10 | 102.21 |
| 7 | NNU_020519 | OlR_40026 | 0.9998 | | 3.0E-72 | 540 | 80.36 | 100.00 |
| 8 | NNU_019581 | OlR_19118 | 0.9997 | endoplasmic reticulum oxidore... | 1.2E-249 | 1491 | 107.58 | 101.64 |
| 9 | NNU_012116 | OlR_07154 | 0.9997 | peroxisomal membrane protei... | 6.5E-112 | 699 | 100.00 | 104.48 |
| 10 | NNU_025150 | OlR_29178 | 0.9996 | Nitrate reductase [NADH] 1 | 0.0 | 2307 | 84.88 | 102.53 |

B. Filter: Pair #103044

**Sequence View for Pairs** 2 of 74957 (<1%)

| SeqID | Descript | Rhiz | Root | Stem | OLeaf |
|---|---|---|---|---|---|
| NNU_010125 | iron-sulfur assembly protein IscA, chloroplastic | 15.25 | 14.24 | 40.41 | 89.91 |
| OlR_26010 | Fe-S_biosyn domain-containing protein | 15.58 | 11.79 | 65.20 | 164.00 |

**Figure S8. PCC of TPM.** (A) OlR-NNU pairs with PCC ≥ 0.8. The AAeval is the E-value for the AA alignment between the sequences. The %Cov1 and %Cov2 are the coverage, which can be greater than 100% due to gaps. (B) The TPM values for a pair with PCC ≥ 0.8.
